# Longitudinal Gut Microbiome Dynamics in Human CTLA-4 pathway Deficiencies: A One-Year Interventional Metagenomic Study

**DOI:** 10.64898/2026.05.18.726014

**Authors:** Máté Krausz, Bei Zhao, Pavla Mrovecova, Michele Proietti, Bodo Grimbacher

## Abstract

**Background:** CTLA-4 haploinsufficiency (CHAI) and LRBA deficiency cause severe immune dysregulation including enteropathy. Abatacept, a CTLA-4-immunoglobulin fusion protein, targets the underlying pathway defect, but its impact on the gut microbiome remains undefined.

**Methods:** We performed longitudinal shotgun metagenomics (MetaPhlAn4/HUMAnN3) on stool samples from patients enrolled in the ABACHAI clinical trial, collected at pre-treatment baseline and months 3, 6, and 12. Healthy individuals from the same household served as controls. Compositional and functional microbiome changes were analyzed using linear mixed-effects models and MaAsLin3, and correlated with organ-specific CHAI Morbidity Scores.

**Results:** At baseline, patients showed significantly reduced alpha diversity (Shannon index, p=0.0029) and distinct community composition (PERMANOVA p=0.0001) compared to healthy controls, characterised by enrichment of oral-associated taxa (*Veillonella, Streptococcus, Lacrimispora*) and depletion of butyrate-producing commensals (*Ruminococcus, Oscillibacter, Dysosmobacter*). Functionally, the baseline metagenome exhibited broad reductions in amino acid and SCFA biosynthesis alongside enrichment of purine salvage and folate pathways. During treatment, beta diversity shifted significantly with treatment duration (Aitchison PERMANOVA R^2^=0.103, p=0.015), with within-patient community turnover peaking at month 6 (Δ=0.216, p=0.006). Longitudinal analyses demonstrated progressive decreases in disease-enriched taxa (*Veillonella, Lacrimispora*) and recovery of commensals (*Collinsella, Adlercreutzia*). FDR-significant reductions in microbial folate and purine biosynthesis pathways were observed over the treatment course. Gut CHAI domain severity correlated inversely with butyrate-producer abundance and positively with oral taxon enrichment.

**Conclusion:** In CTLA-4 pathway insufficiency patients, abatacept therapy is associated with an improvement of enteropathy and a progressive, measurable gut microbiome restructuring, positioning microbiome dynamics as a candidate biomarker of treatment response in this monogenic immune dysregulation disorder.

## Introduction

Cytotoxic T-lymphocyte-associated protein 4 (CTLA-4) is an inhibitory checkpoint of immune activation, therefore a protein with a central role regulating the immune homeostasis. CTLA-4 is constitutively expressed on regulatory T cells (Tregs) and gets upregulated on conventional T cells upon activation.(1) CTLA-4 outcompetes the co-stimulatory receptor CD28 for their shared ligands CD80 and CD86 on antigen-presenting cells (APCs), thereby regulating T-cell proliferation. Through the process of transendocytosis, CTLA-4 physically removes its ligands from the surface of APCs; the process itself is dependent on the lipopolysaccharide-responsive and beige-like anchor protein (LRBA) (2–4). Homozygous deficiency of *Ctla4* in mice is lethal due to fulminant lymphoproliferation, underscoring the non-redundant role of this checkpoint in immune regulation (1). Heterozygous loss of function mutations in humans (in contrast to mice, where - at least under SPF conditions - no obvious phenotype can be detected) lead to a disease called CTLA-4 (haplo)insufficiency with autoimmune infiltration (CHAI). These mutations lead to impaired CTLA-4 expression or function, resulting in Treg dysfunction, hyperactivation of effector T cells, and lymphocytic infiltration of non-lymphoid organs (1,2). In the largest published cohort of 133 mutation carriers from 54 families, the clinical penetrance was estimated at approximately 67%. The clinical spectrum of affected patients includes hypogammaglobulinemia (84%), lymphoproliferation (73%), autoimmune cytopenia (62%), and respiratory (68%), gastrointestinal (59%), and neurological (29%) involvement (5). Notably, disease manifestation can range from asymptomatic carrier status to life-threatening multiorgan infiltration, even among carriers of the same mutation within the same family, indicating the influence of additional genetic (like HLA), epigenetic, and environmental modifiers on disease expression (6–8). Gastrointestinal (GI) involvement is one of the most common and a clinically highly significant manifestation of CTLA-4 insufficiency. The GI involvement is characterized by lymphocytic infiltration of the intestinal mucosa, leading to chronic diarrhea, and malabsorption. Additional gastrointestinal features include celiac-like disease, inflammatory bowel disease-like phenotypes, and early-onset Crohn’s disease. As LRBA regulates CTLA-4 turnover, its absence leads to increased CTLA-4 removal from the cell surface, exacerbating immune activation. Hence, the phenotype of LRBA deficiency is very similar to CTLA-4 insufficiency: both conditions share similar clinical features, with LRBA deficiency being generally more severe.

Current therapeutic strategies for treating CTLA-4-associated immune dysregulation include immunosuppressive agents, such as corticosteroids, sirolimus, mycophenolate mofetil or cyclosporine. Treatment with abatacept (a CTLA-4-immunoglobulin fusion protein) represents a targeted approach, as abatacept address the underlying pathway defect. In the largest therapeutic analysis to date, abatacept ameliorated disease severity across multiple organ domains, particularly gastrointestinal and pulmonary involvement (9). The ABACHAI clinical trial was designed to prospectively evaluate the safety and efficacy of subcutaneous abatacept in this population (10). The trial demonstrated a favorable safety profile for abatacept in this indication with moderate efficacy, supporting its role as a targeted immunomodulatory therapy (manuscript submitted).

The gut microbiome is being increasingly recognized as one of the central modulators of immune homeostasis (11). Gut dysbiosis has been implicated in the pathogenesis of multiple immune-mediated diseases, like rheumatoid arthritis (12), systemic lupus erythematodes (13) or inflammatory bowel disease (14). In the context of inborn errors of immunity (IEI), disruption of the delicate host-microbe balance is of particular relevance, as patients harbor genetically determined immune defects that directly affect microbial control at mucosal surfaces. The most well studied is the connection between disease and microbiome amongst IEIs is in common variable immunodeficiency (CVID) (15–17). In CVID, decreased alpha diversity, enrichment of potentially pathogenic taxa such as *Enterococcus*, and depletion of short-chain fatty acid-producing commensals have been linked to systemic inflammation and immune dysregulation (16,18). Importantly, IgA deficiency — usually present in many antibody deficiency syndromes, including CVID as well — disrupts mucosal immunoglobulin-mediated microbial selection, promoting dysbiosis (16,19). These observations collectively suggest that impaired adaptive immunity shapes a permissive microbial milieu that may, in turn, perpetuate or exacerbate immune dysregulation.

The relationship between CTLA-4 and the gut microbiome has been extensively studied in the oncological setting, where immune checkpoint inhibitors (ICI), blocking CTLA-4 (like ipilimumab) are used for cancer immunotherapy. Vétizou *et al*. showed that the antitumor efficacy of CTLA-4 blockade depends on distinct *Bacteroides* species, establishing a direct link between CTLA-4 signaling, gut microbiota composition, and systemic immune responses (20,21). Subsequent work has shown that immune checkpoint inhibitor-related colitis is associated with microbiota-dependent activation of CD4+ T cells and selective depletion of peripherally induced Tregs in the gut (21). These findings are directly relevant to inborn errors of the CTLA-4 pathway, where a genetically determined reduction of CTLA-4 mimics the ICI-induced blockade, but chronically.

Despite the close connection of the CTLA-4 insufficiency pathology to the gut, the intestinal microbiome in this condition remained unexplored until recently. In the first and currently only published study, we - together with our collaborators from the NIH - have characterized the intestinal microbiome and metabolome in two geographically distinct cohorts of patients with CTLA-4 deficiency (NIH, United States, n = 32; CCI, Freiburg, Germany, n = 25) (22). We found that the genera *Veillonella* and *Streptococcus* are disease-specific biomarkers that distinguished CTLA-4 patients from both healthy controls and patients with CVID. In addition, our studies showed that disease severity and gastrointestinal involvement were associated with further microbiome and metabolome alterations. In addition, we found that treatment with abatacept and/or sirolimus partially restored microbial diversity and reversed several dysbiotic features, suggesting that microbiome remodeling may accompany clinical improvement. However, this study was cross-sectional in design and thus could not capture the temporal dynamics of microbiome changes during treatment.

Here we present a longitudinal shotgun metagenomic analysis of the gut microbiome in patients with CTLA-4 haploinsufficiency undergoing treatment with abatacept. We have prospectively collected stool samples from nine participants of the ABACHAI trial (10) at up to four time points over the one-year long treatment period (pre-treatment, months 3, 6, and 12). Using shotgun metagenomics processed through we characterized both compositional and functional changes in the gut microbiome and compared these to organ-specific disease severity measured by the CHAI-Morbidity Score (10). To our knowledge, this represents the first study to track gut microbial dynamics prospectively during targeted immunomodulatory therapy in a monogenic immune dysregulation disorder.

## Methods

### Sample collection

Stool samples from patients and healthy individuals were collected, as described before (15). Patients being treated as part of the ABACHAI clinical trial at the CCI Outpatient clinic, Medical Center, University of Freiburg were recruited to participate in the study after ethical approval by the local ethics committee of the University of Freiburg (protocol no. 466/18). The ABACHAI clinical trial was a phase II prospective interventional clinical trial. Patient during their trial visits received pre-prepared packages including stool collection tubes (Invitek) and a lifestyle questionnaire about diet, environment, current symptoms and medications as well a brief clinical history. The samples were shipped by the patients to our laboratory via the German post. Upon arrival, samples were aliquoted and frozen at -20 °C and stored at -80 °C on the long term. From the 20 participants of the ABACHAI trial only 10 patients provided baseline, and 9 patients follow up stool samples, all of them being treated at the Freiburg trial site. Samples were collected longitudinally: a baseline sample before trial inclusion, as well as follow up samples at month 3, 6, 12 of the trial. Not all patients provided a sample at each visit; the analyzed sample are summarized in Table 1.

**Table 1.** Summary of analyzed sample information. The column assigned visit date refers to the factorial analysis, where -1 referst to the baseline visit, 3, 6 and 12 to the respective study visit month.

| SampleID | Patient ID | Group | Time difference to baseline [months] | Assigned date | Abatacept | Sex | Patient group | Antibiotics | Antibiotics Info | Probiotics | Probiotic Info |
| --- | --- | --- | --- | --- | --- | --- | --- | --- | --- | --- | --- |
| 01/001-1 | 01/001 | patient | 0 | -1 | 0 | Female | CTLA4 | 1 | Cotrimoxazole | 0 | - |
| 01/001-2 | 01/001 | patient | 3.4825462 | 3 | 1 | Female | CTLA4 | 1 | Cotrimoxazole | 0 | - |
| 01/001-3 | 01/001 | patient | 9.9219713 | 6 | 1 | Female | CTLA4 | 1 | Cotrimoxazole | 0 | - |
| 01/001-4 | 01/001 | patient | 13.174538 | 12 | 1 | Female | CTLA4 | 1 | Cotrimoxazole | 0 | - |
| 01/004-1 | 01/004 | patient | -2.299795 | -1 | 0 | Male | CTLA4 | 0 | - | 0 | - |
| 01/004-2 | 01/004 | patient | 6.8665298 | 6 | 1 | Male | CTLA4 | 0 | - | 0 | - |
| 01/006-1 | 01/006 | patient | -0.328542 | -1 | 0 | Female | CTLA4 | 0 | - | 0 | - |
| 01/006-2 | 01/006 | patient | 3.9096509 | 3 | 1 | Female | CTLA4 | 0 | - | 0 | - |
| 01/006-3 | 01/006 | patient | 6.3408624 | 6 | 1 | Female | CTLA4 | 0 | - | 0 | - |
| 01/006-4 | 01/006 | patient | 11.827515 | 12 | 1 | Female | CTLA4 | 0 | - | 0 | - |
| 01/009-1 | 01/009 | patient | -3.12115 | -1 | 0 | Female | CTLA4 | 0 | - | 0 | - |
| 01/009-2 | 01/009 | patient | 6.5051335 | 6 | 1 | Female | CTLA4 | 0 | - | 0 | - |
| 01/009-3 | 01/009 | patient | 5.7494867 | 3 | 1 | Female | CTLA4 | 0 | - | 0 | - |
| 01/009-4 | 01/009 | patient | 12.648871 | 12 | 1 | Female | CTLA4 | 0 | - | 0 | - |
| 01/010-1 | 01/010 | patient | -4.041068 | -1 | 0 | Male | CTLA4 | 0 | - | 0 | - |
| 01/010-2 | 01/010 | patient | 3.1211499 | 3 | 1 | Male | CTLA4 | 0 | - | 0 | - |
| 01/010-3 | 01/010 | patient | 6.6694045 | 6 | 1 | Male | CTLA4 | 0 | - | 0 | - |
| 01/011-1 | 01/011 | patient | -1.774127 | -1 | 0 | Female | CTLA4 | 1 | Levofloxacin | 0 | - |
| 01/011-2 | 01/011 | patient | 3.3511294 | 3 | 1 | Female | CTLA4 | 0 | - | 0 | - |
| 01/012-1 | 01/012 | patient | -0.525667 | -1 | 0 | Male | CTLA4 | 0 | - | 0 | - |
| 01/012-2 | 01/012 | patient | 3.8439425 | 3 | 1 | Male | CTLA4 | 0 | - | 0 | - |
| 01/012-3 | 01/012 | patient | 8.0492813 | 6 | 1 | Male | CTLA4 | 0 | - | 0 | - |
| 01/012-4 | 01/012 | patient | 13.240246 | 12 | 1 | Male | CTLA4 | 0 | - | 0 | - |
| 01/013-1 | 01/013 | patient | -1.445585 | -1 | 0 | Female | CTLA4 | 0 | - | 0 | - |
| 01/016-1 | 01/016 | patient | -0.197125 | -1 | 0 | Female | LRBA | 0 | - | 0 | - |
| 01/016-2 | 01/016 | patient | 7.8850103 | 6 | 1 | Female | LRBA | 0 | - | 0 | - |
| 01/016-3 | 01/016 | patient | 12.517454 | 12 | 1 | Female | LRBA | 0 | - | 1 | Mutaflor |
| 05/1489-1 | 05/1489 | control | - | -1 | -1 |  | control | 0 | - | 0 | - |
| 05/244-1 | 05/244 | control | - | -1 | -1 |  | control | 0 | - | 0 | - |
| 05/245-1 | 05/245 | control | - | -1 | -1 |  | control | 0 | - | 0 | - |
| 05/259-1 | 05/259 | control | - | -1 | -1 |  | control | 0 | - | 0 | - |
| 05/266-1 | 05/266 | control | - | -1 | -1 |  | control | 0 | - | 0 | - |
| 05/4379-1 | 05/4379 | control | - | -1 | -1 |  | control | 0 | - | 0 | - |
| 05/4468-1 | 05/4468 | control | - | -1 | -1 |  | control | 1 | Ciprofloxacin (3 days) | 0 | - |
| 05/4470-1 | 05/4470 | control | - | -1 | -1 |  | control | 0 | - | 0 | - |
| 05/558-1 | 05/558 | control | - | -1 | -1 |  | control | 0 | - | 0 | - |
| 05/573-1 | 05/573 | control | - | -1 | -1 |  | control | 0 | - | 0 | - |
| 05/6964-1 | 05/6964 | control | - | -1 | -1 |  | control | 0 | - | 0 | - |
| 05/7782-1 | 05/7782 | control | - | -1 | -1 |  | control | 1 | Penicillin 4 weeks before sampling | 0 | - |
| 05/7796-1 | 05/7796 | control | - | -1 | -1 |  | control | 0 | - | 0 | - |
| 05/7801-1 | 05/7801 | control | - | -1 | -1 |  | control | 0 | - | 0 | - |
| 05/7950-1 | 05/7950 | control | - | -1 | -1 |  | control | 0 | - | 0 | - |
| 05/8061-1 | 05/8061 | control | - | -1 | -1 |  | control | 0 | - | 0 | - |
| 05/8442-1 | 05/8442 | control | - | -1 | -1 |  | control | 0 | - | 0 | - |

### Shotgun metagenomics

For shotgun metagenomics DNA was extracted using the AllPrep PowerFecal Pro DNA/RNA kit following the manufacturer’s instructions. DNA quantity and integrity were assessed using Nanodrop ND2000 (Thermo Fisher Scientific, USA). The extracted DNA samples were shipped to Novogene (Cambridge, United Kingdom), where DNA library preparation and sequencing was performed.

A total amount of 1 μg DNA per sample was used as input. Sequencing libraries were generated using NEBNext® Ultra™ DNA Library Prep Kit for Illumina (NEB, USA) following manufacturer’s recommendations and index codes were added to attribute sequences to each sample. Briefly, the DNA sample was fragmented by sonication to a size of 350bp, then DNA fragments were end-polished, A-tailed, and ligated with the full-length adaptor for Illumina sequencing with further PCR amplification. At last, PCR products were purified (AMPure XP system, Beckman Coulter) and libraries were analysed for size distribution by Agilent2100 Bioanalyzer and quantified using real-time PCR. The clustering of the index-coded samples was performed on a cBot Cluster Generation System according to the manufacturer’s instructions. After cluster generation, the library preparations were sequenced on an Illumina NovaSeq 6000 platform and paired-end reads were generated with an average depth of 73,8 million pairedl⍰end reads per sample.

For analysis of the bacteria, reads were initially processed with kneaddata v0.12.0. The trimmed forward and reverse reads merged and handed off for further processing with Humann v3.9 (23) and Metaphlan v4.1.1 (24,25) with database version mpa_vJun23_CHOCOPhlAnSGB_202307. Downstream statistical analysis was performed in R (v4.5.2) using various packages (vegan (26), MaAsLin3 (27), lmertest (28), ggplot2 (29)).

## Results

### Cohort characteristics and sampling overview

From the 20 ABACHAI trial participants nine patients (eight CTLA-4 and one LRBA) provided altogether 27 stool samples. Eight of the trial participants’ household controls provided a baseline control sample, which cohort was expanded with additional 9 samples (household controls of other, non-trial participating CTLA4 patients). Due to rare trial population and limited sample availability, antibiotic use was not excluded: patient 01/001 was on continuous prophylactic cotrimoxazole therapy due to lymphopenia; patient 01/011 received levofloxacine treatment for a respiratory infection at baseline; 1 healthy control received a 3-day treatment with ciprofloxacin before sampling, another received penicillin 4 weeks prior sampling (Table 1).

### Patients at baseline have a dysbiosis specific for CTLA-4 pathway defects

Comparing baseline samples of the trial participants (9 samples) to the healthy control cohort (17 samples), we observed a significantly decreased alpha diversity (Shannon-index, Mann-Whitney U-Test, p=0.0029, Figure 1A). Beta-diversity testing revealed significant differences between the two groups (PERMANOVA (adonis2), R2= 0.099, F=2.65, p=0.0001). Differences in multivariate dispersion between groups did not reach statistical significance (betadisper ANOVA: F=3.27, p=0.083, permutation test: p=0.083) indicating that the observed clustering primarily reflects a shift in community composition rather than systematically increased within-group variability in either cohort. Altogether the patient group exhibited a more heterogenous microbiome composition (Figure 1B). A genus level differential abundance analysis using MaAsLin3 (TSS normalization, log⍰transformed abundances) revealed two significantly different features after multiple testing correction (q<0.1): the genus *Lacrimispora* and an unclassified species⍰level genome bin (GGB9760). The relative abundance of *Lacrimispora* was higher in the patient group (β=3.55, SE=0.65, q=0.027), whereas GGB9760 were strongly depleted in patients (β=−6.42, SE=1.27, q=0.027), with its prevalence association flagged as secondary to the abundance effect. Beyond these FDR⍰significant taxa, several further genera that have been previously implicated in CTLA4-deficiency showed concordant (but nominal) associations, including higher relative abundance of *Veillonella, Streptococcus, Lactobacillus/Limosilactobacillus, Parasutterella*, and *Eggerthella*, and lower relative abundance of *Ruminococcus, Oscillibacter, Dysosmobacter* and *Prevotella*, patients compared to controls (Figure 1C and Supplementary Table 1).

**Figure 1.**
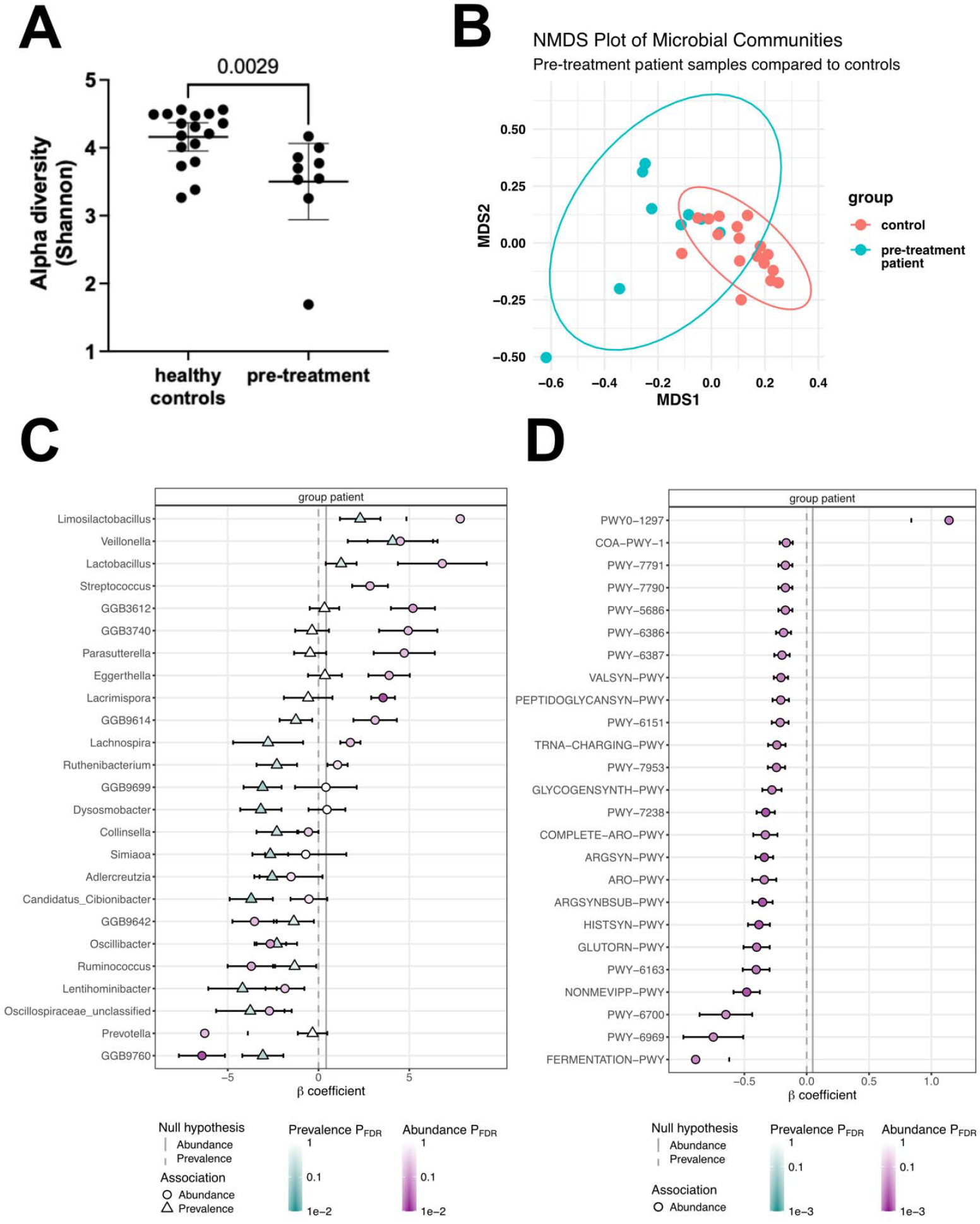
(A) Patients with defects in the CTLA-4 pathway have a significantly decreased alpha diversity (Shannon-index) before starting abatacept treatment (Mann-Whitney U-test). (B) Beta-diversity (Bray-Curtis) analysis and subsequent PERMANOVA-testing shows statistically significant compositional differences between patients and healthy controls before initiating abatacept treatment. (C) Differential abundance testing (MaAsLin3) at genus levels, showing the top 25 genera after q-value ranking. (D) Differentially abundant pathways (MaAsLin3) comparing pre-treatment samples to healthy controls showing the top 25 significant genera after q-value ranking; complete pathway names are listed in Supplementary Table 2.

Functional profiling revealed large differences in microbial metabolic pathways between the patients and healthy controls. Using HUMAnN3 unstratified CPM⍰normalized pathways as an input for MaAsLin3, we identified a coordinated reduction in multiple core biosynthetic and energy metabolism pathways in CTLA-4⍰pathway deficient individuals (Figure 1D, full pathway names listed in Supplementary Table 2). De novo biosynthesis of several amino acids, including arginine, L-ornithine, histidine, valine, tryptophan and aromatic amino acids, was significantly decreased in patients (β ≈ −0.2 to −0.5, q<0.1). We also observed reduced abundance of pathways involved in peptidoglycan precursor synthesis, central carbon metabolism, including mixed acid fermentation and a variant tricarboxylic acid cycle, and several cofactor biosynthesis routes (tetrapyrrole, queuosine, coenzyme A). In contrast, pathways mediating purine deoxyribonucleoside degradation (PWY0⍰1297, β=1.14, q=0.025) and purine nucleotide salvage (PWY66⍰409, β=0.81, q=0.074) were relatively enriched in CTLA4⍰deficient patients, and a bacterial superpathway of tetrahydrofolate biosynthesis (PWY⍰6612) showed a large increase (β=1.04, q=0.096). For most pathways, prevalence was 100% across samples, indicating that these differences reflect quantitative shifts in pathway abundance rather than presence/absence.

### Microbial communities show a shift during abatacept treatment

Alpha diversity (Shannon-index) was analyzed longitudinally across the clinical visits in the trial in 9 patients (27 samples total, Figure 2A). To account for the unbalanced longitudinal design, differences in Shannon-index between treatment timepoints and baseline were assessed using a linear mixed-effects model with patient as a random intercept. Baseline Shannon diversity (mean±SE=3.50±0.18) showed a gradual but not statistically significant (BH-adjusted) increase at month 3 (β=0.27, SE=0.27, p=0.341), month 6 (β =0.42, SE=0.26, p=0.125), and month 12 (β=0.43, SE=0.29, p=0.156). A continuous trend model further supported a positive but non-significant trajectory, estimating a Shannon diversity increase of 0.030 per month from baseline (SE=0.019, p=0.131).

**Figure 2.**
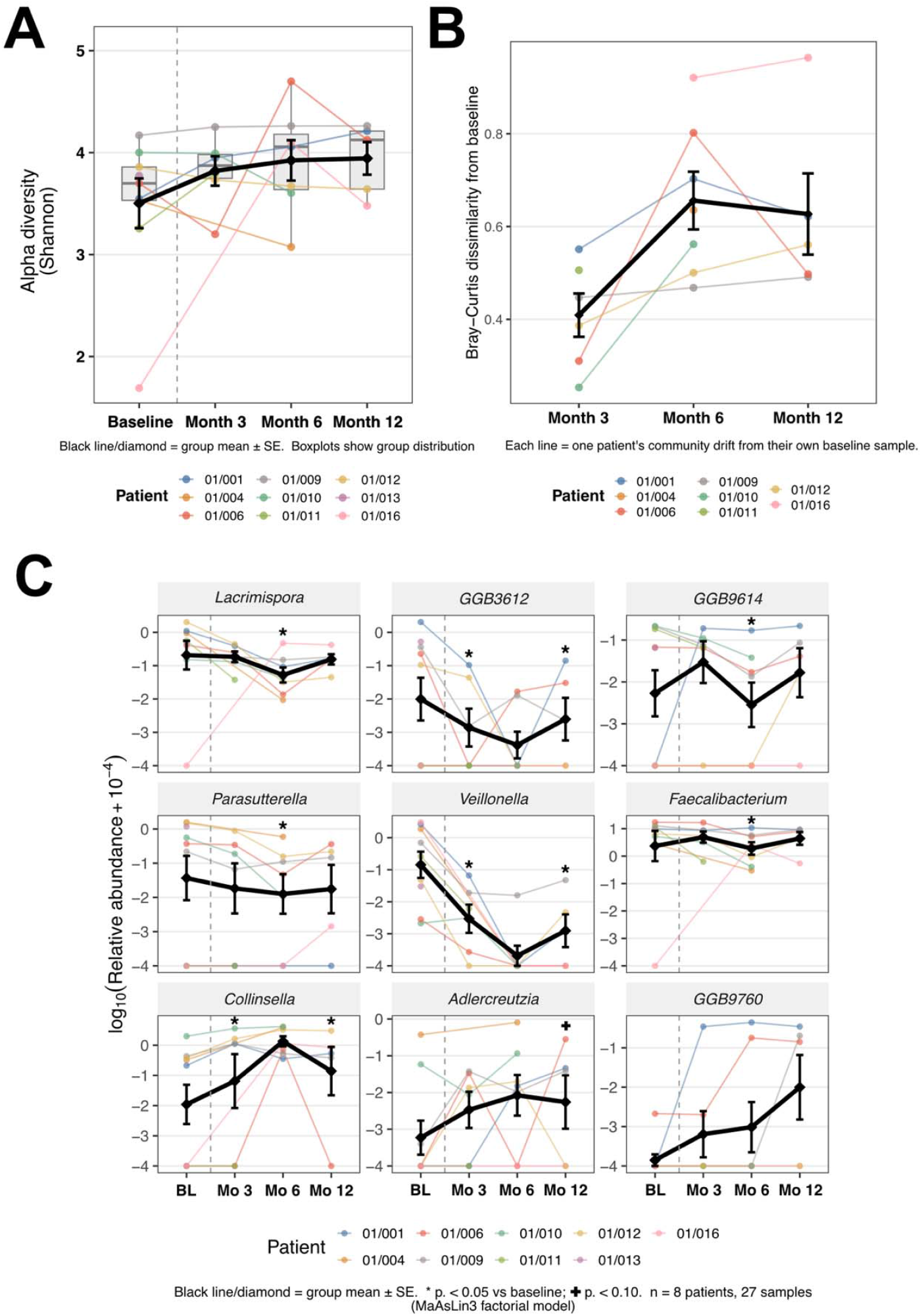
Longitudinal microbiome changes during abatacept treatment in patients with CTLA-4 pathway deficiency. (A) Alpha diversity (Shannon-index) across baseline and on-treatment timepoints. Boxplots show group distribution; colored lines and points represent individual patients; black lines and diamonds represent group mean±SE. (B) Within-patient Bray-Curtis dissimilarity from baseline, used as a measure of individual community turnover during treatment (n=8 patients) (C) Longitudinal trajectory plots for selected candidate genera identified from the baseline analysis and longitudinal mixed-effects modeling. Colored lines and points represent individual patients; black lines and diamonds represent group mean±SE. Asterisks indicate p<0.05 and daggers indicate p<0.10 versus baseline in the factorial MaAsLin3 model; all associations in panel C are exploratory.

Within-patient community turnover was assessed by the Bray-Curtis dissimilarity index between each patient’s sample during the treatment phase compared to their own baseline sample, to measure the treatment-associated individual community drift (Figure 2B). Turnover was modelled using a linear mixed-effects model with on-site visit as a fixed effect and patient as a random intercept. Mean Bray-Curtis dissimilarity from baseline was 0.445 (SE=0.063) at month 3, this increased significantly to 0.661 at month 6 (p=0.006) and remained elevated at month 12 compared to month 3 (p=0.035), though with a slight attenuation relative to the peak at 6 months. A continuous trend model further supported a positive association between time from baseline and community dissimilarity, though this did not reach conventional significance (β=0.019 per month, p=0.081), consistent with the non-monotonic trajectory observed in the factorial model.

### Taxonomic responders to abatacept therapy

To evaluate taxonomical dynamics over the course of abatacept treatment, we performed longitudinal mixed-effects analysis (MaAsLin3, log-transformed relative abundance, random intercept per patient) in 9 treatment-naïve participants. For this, we have pre-selected 25 candidate genera, based on the baseline results. Two complementary analyses were performed: a factorial model comparing each post-baseline timepoint against baseline, and a continuous trend model estimating monotonic change per month from baseline. Given the limited sample size (not all patients provided samples at all given timepoints, Table 1) no taxon reached FDR significance (q<0.1) in either model. At a nominal threshold (p<0.05) applied to a candidate panel derived from baseline differential abundance analysis, both models identified coherent and directionally consistent changes, supporting the hypothesis of progressive microbiome remodeling during treatment with abatacept.

The factorial model revealed two contrasting longitudinal patterns (Figure 2C). First, several taxa enriched in patients at baseline showed decreasing pattern during treatment: *Lacrimispora* (month 6: β=−3.38, p=0.010), GGB3612 (month 3: β=−4.40, p=0.022; month 12: β=−4.23, p=0.034), GGB9614 (month 6: β=−2.53, p=0.018), *Parasutterella* (month 6: β=−2.93, p=0.027), and *Veillonella* (month 3: β=−4.13, p=0.042; month 12: β =−5.54, p=0.031); altogether a partial normalisation of the disease-associated microbiome signature. Second, *Collinsella* — depleted in patients at baseline relative to healthy controls — showed sustained increase across all on-treatment timepoints (month 3: β=1.89, p=0.022; month 6: β=1.67, p=0.053; month 12: β=1.30, p=0.043), suggesting treatment-associated recolonisation of a previously suppressed niche. Additionally, *Faecalibacterium* decreased temporarily at month 6 (β=−2.11, p=0.046) despite showing no difference at baseline.

The continuous trend model, estimating abundance change per month from baseline, confirmed the two strongest normalizing signals: *Veillonella* (β=−0.41/month, p=0.033) and GGB3612 (β=−0.34/month, p=0.041). Additionally, *Adlercreutzia* showed a consistent, gradual increase (β=0.25/month, p=0.051) that was not present in the factorial model, suggesting a slow but sustained colonisation dynamic over the full 12-month period. *Parasutterella* and GGB9760 showed supporting trends in the same direction (p=0.059 and p=0.061, respectively).

### Functional pathway shifts during abatacept treatment

To assess longitudinal changes in the functional capacity of the gut microbiome during abatacept treatment, we performed MaAsLin3 analysis on a candidate panel of 25 pathways identified based on the baseline patient-versus-control analyses, focusing on functional categories including nucleotide metabolism, folate/one-carbon metabolism, short-chain fatty acid fermentation, amino acid biosynthesis, and immune-relevant pathways. Similarly to the taxonomical longitudinal analysis, a linear mixed model with subject as a random intercept to account for repeated measures was used with two complementary modeling strategies: a factorial and a continuous linear model. Both approaches independently identified a progressive reduction in microbial folate biosynthesis (factorial model: month 6: coef≈−0.79 to −0.82, p < 0.04; month 12: coef ≈ −1.07 to −1.11, p < 0.022; linear model: coef≈−0.077 to −0.080 per month, p<0.009, q=0.075) and purine metabolism (factorial model month 12: p < 0.042; linear model: p < 0.029) (Figure 3, Supplementary Tables 3 and 4). No significant shifts were observed in butyrate fermentation, aromatic amino acid biosynthesis, or immune-activating structural pathways. Although results did not uniformly reach FDR-corrected significance at q < 0.1, the convergence of effect direction and magnitude across orthogonal modeling strategies strengthens confidence in these candidate pathway associations.

**Figure 3.**
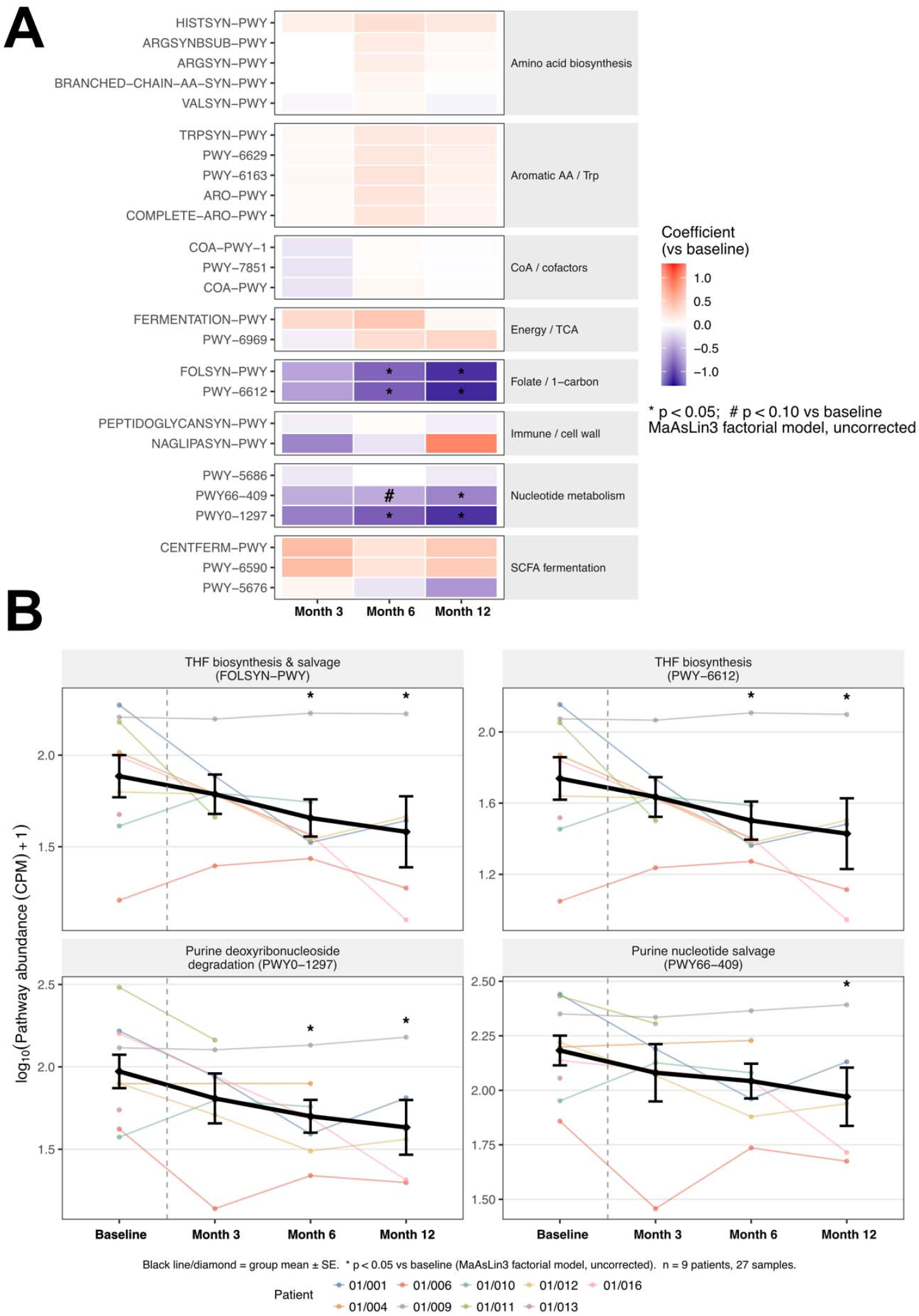
Longitudinal changes in microbial metabolic pathways during abatacept treatment. (A) Heatmap of MaAsLin3 factorial abundance model for the candidate pathway panel across the trial relative to baseline; blue indicates decreased pathway abundance and red indicates increased abundance compared with baseline (B) Longitudinal trajectory plots for the 4 pathways showing the strongest and most consistent treatment-associated changes: FOLSYN-PWY (superpathway of tetrahydrofolate biosynthesis and salvage), PWY-6612 (superpathway of tetrahydrofolate biosynthesis), PWY0-1297 (superpathway of purine deoxyribonucleosides degradation), and PWY66-409 (superpathway of purine nucleotide salvage). Colored lines and points represent individual patients; black lines and diamonds represent group mean±SE. Asterisks indicate nominally significant differences versus baseline in the factorial model.

### Associations Between Gut Microbiome Composition and CHAI Disease Domains

The CHAI score was developed alongside the ABACHAI trial to assess disease activity during abatacept treatment. the score consists of six domains (gut, lung, cytopenia, immune system, lymphoproliferation and skin), each gets scored from 0 (none) to 3 (severe), for overall disease activity the mean of all assessed domains can be used (Figure 4A). To investigate organ-specific associations between microbiome composition and disease severity in CTLA-4 pathway insufficiency, we performed domain-resolved MaAsLin3 analyses across all detected genera (abundance and prevalence models, random intercept per patient), using each of the CHAI sub-domain scores as separate continuous predictors. The central nervous system (CNS) domain was excluded from interpretation due to insufficient observations. No (except one) association reached FDR-corrected significance threshold at q<0.1 across either model type; results are therefore reported at nominal significance (p<0.05, |coef|>1).

**Figure 4.**
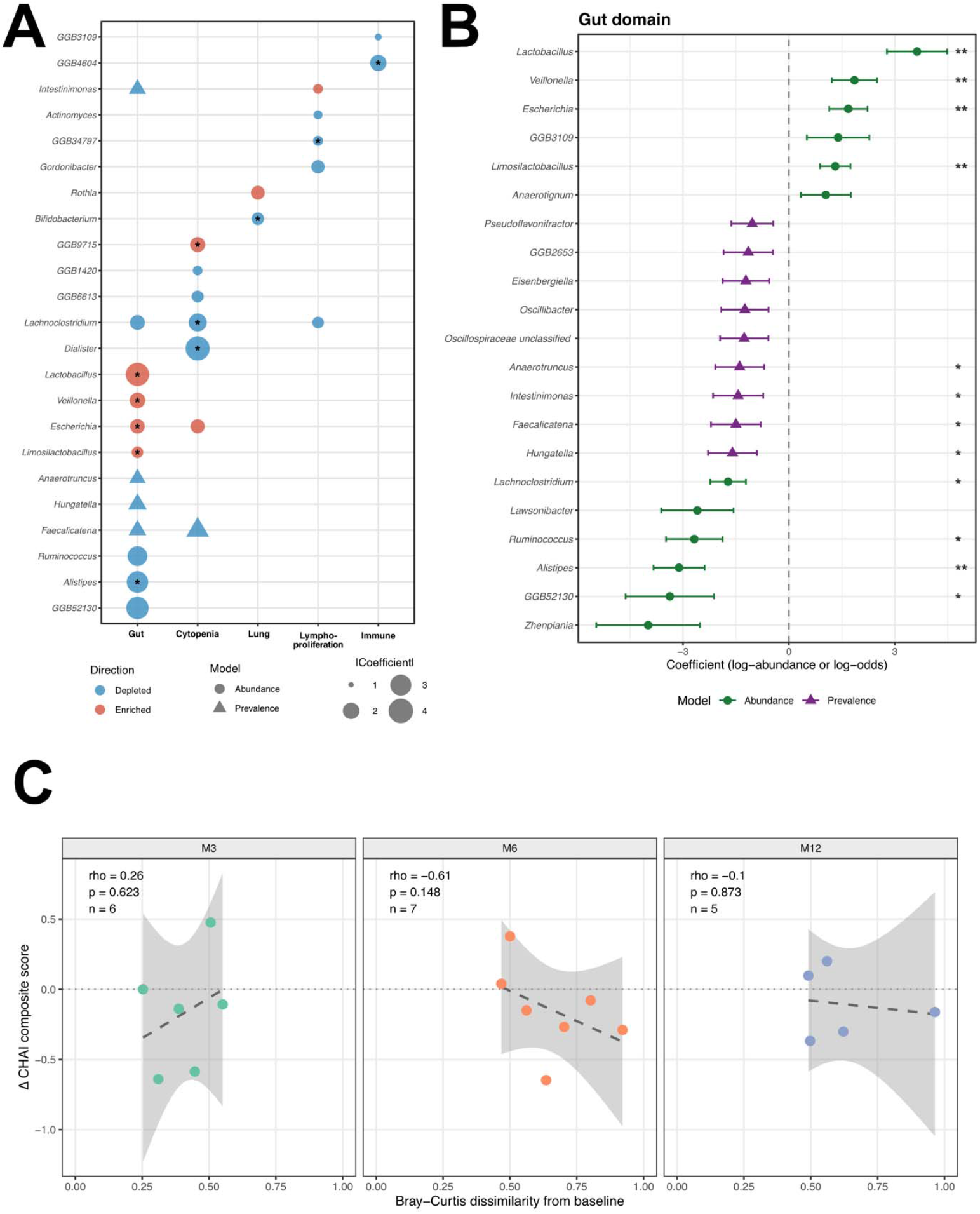
Domain-specific associations between gut microbiome composition and CHAI disease severity scores. (A) Bubble plot showing all genera with nominal associations (p<0.05, |coef|>1) across five CHAI domains. Bubble size represents effect size (|coefficient|); colour indicates direction of association relative to domain severity score; shape distinguishes abundance (circle) and prevalence (triangle) model associations. Asterisks indicate associations at p<0.01; all displayed associations are at p<0.05 with |coef|>1. (B) Forest plot of gut domain associations showing coefficients ± SE for abundance (green) and prevalence (purple) models for features p<0.1. Significance: ** p<0.01, * p<0.05; results are exploratory. (C) Scatter plots showing the Spearman correlation between within-patient Bray-Curtis dissimilarity from baseline (x-axis) and change in CHAI composite score from baseline (ΔCHAI, y-axis) at month 3 (n=6), month 6 (n=7), and month 12 (n=5). Each point represents one patient. ΔCHAI was calculated as the mean of available CHAI domain scores at each timepoint minus the corresponding baseline value; negative values indicate clinical improvement. Dashed lines represent linear regression fits with 95% confidence intervals. Spearman rho and p-values are indicated within each panel.

The gut domain yielded the most coherent and biologically consistent signal. Higher gut CHAI scores were associated with reduced abundance of butyrate-producing commensals, including *Alistipes* (coef=−3.11, p=0.002), *Ruminococcus* (coef=−2.68, p=0.015), and *Lachnoclostridium* (coef=−1.72, p=0.033), and with enrichment of oral and facultative anaerobe taxa, including *Escherichia* (coef=1.68, p=0.003), *Veillonella* (coef=1.86, p=0.003), and *Lactobacillus/Limosilactobacillus* (coef=3.63/1.31, p<0.01). Complementary prevalence model associations further identified reduced colonisation probability for *Hungatella* (coef=−1.60, p=0.021), *Faecalicatena* (coef=−1.50, p=0.033), *Intestinimonas* (coef=−1.44, p=0.043), *Anaerotruncus* (coef=−1.39, p=0.044), and *Oscillibacter* (coef=−1.25, p=0.062) with increasing gut severity, indicating ecological exclusion beyond quantitative abundance reduction (Figure 4B). The directional concordance between the gut domain microbiome-severity associations and the patient-versus-control baseline dysbiosis signature provides internal cross-validation of the candidate taxa identified in the primary differential abundance analysis.

As for the cytopenia domain a signal for *Lachnoclostridium* was identified, which showed inverse abundance associations with the cytopenia domain (coef=−2.31, p=0.0003, q=0.054*) -* the only near FDR-significant clean association across the entire domain analysis - as well as nominal associations (p<0.05) with the gut and lymphoproliferation domains. The convergence of this Lachnospiraceae member across three independent CHAI sub-scores suggests that its depletion may reflect a systemic disease activity association rather than isolated gut involvement. Additional cytopenia-associated depletions included *Dialister* (coef=−3.88, p=0.006) and *Faecalicatena* (prevalence: coef=−2.10, p=0.044). In the lung domain (n=20 samples with lung involvement), exploratory associations included depletion of *Bifidobacterium* (coef=−1.41, p=0.009) and enrichment of *Rothia* (coef=1.57, p=0.045). (Figure 4A and 4B, Supplementary Table 5 and 6).

To directly test whether the magnitude of microbiome restructuring observed during treatment was associated with clinical response, we correlated within-patient Bray-Curtis community turnover with ΔCHAI composite score (average of all domains) at matched timepoints (Figure 4C). Spearman correlation between within-patient Bray-Curtis dissimilarity from baseline and ΔCHAI composite score was assessed separately at each timepoint to avoid conflation with the time-dependent increase in community turnover. At month 6, where community turnover was maximal, a moderate negative correlation was observed (rho=−0.61, p=0.148, n=7), suggesting that patients with greater microbiome restructuring tended toward larger reductions in disease severity. No associations were observed at month 3 (rho=+0.26, p=0.623, n=6) or month 12 (rho=−0.10, p=0.873, n=5). The pooled cross-timepoint correlation was near zero (rho=−0.13, p=0.604, n=18), reflecting the confounding effect of time on Bray-Curtis dissimilarity when timepoints are aggregated. These findings are hypothesis-generating and consistent with a model in which microbiome remodeling and clinical response are partially coupled during active immunomodulation, with the greatest concordance at the timepoint of peak community turnover.

## Discussion

Our study is the first prospective, interventional longitudinal metagenomic characterization of the gut microbiome in patients with CTLA-4 pathway insufficiency undergoing treatment with abatacept. Our results show that patients prior targeted CTLA-4 pathway reconstitution have a distinct, functionally impoverished gut microbiome, and that twelve months of treatment is associated with a measurable, progressive restructuring of the gut microbiome – even though a full recovery remained incomplete within the limited observation window.

The pre-treatment microbiome of ABACHAI trial participants confirmed the dysbiotic profile that our collaborators and we have reported in the currently largest published cross-sectional study of CTLA-4 insufficiency, which characterized two geographically distinct cohorts totaling 57 patients (22). Enrichment of oral-associated taxa (*Veillonella, Streptococcus, Lactobacillus/Limosilactobacillus, Parasutterella*) alongside depletion of butyrate-producing Clostridiales (*Ruminococcus, Oscillibacter, Dysosmobacter, Odoribacter*) were reproduced here in a separate, much smaller cohort. The replication of this signature across another cohort processed with a different pipeline (16S-amplicon vs. metagenomics) supports causality to the underlying immunodeficiency. At the functional level, the baseline metagenome showed a broad reduction in *de novo* amino acid biosynthesis (tryptophan, branched-chain, aromatic), central fermentation and TCA cycle pathways, and cofactor synthesis, mirroring the metabolomic findings from the cross-sectional study (22). Mechanistically, this compositional-functional profile is consistent with the pattern observed in immune checkpoint inhibitor (ICI)-related colitis during anti-CTLA-4 therapy, where a colitogenic microbiome characterized by oral-taxon translocation and SCFA-producer depletion associated with gut toxicity (20,30–32). CTLA-4 haploinsufficiency can thus be considered a chronic, genetically determined model of checkpoint blockade in the gut, in which years of reduced CTLA-4 signaling select for a dysbiotic community long before treatment initiation.

Alpha diversity showed a positive (but non-significant) trend across all timepoints consistent with the variable clinical response observed in the trial (manuscript submitted). Beta diversity analysis provided stronger evidence: treatment duration was a significant predictor of community composition by Aitchison distance – which weights log-ratio shifts uniformly rather than being dominated by abundant taxa – indicates broad, proportional community restructuring rather than shifts in a few dominant species. Within-patient community turnover from baseline peaked at month 6 (Δ=+0.216, p=0.006) and remained elevated at month 12, suggesting that the period of active immune suppression during months 3–6 constitutes the primary window of ecological reorganization, followed by partial stabilization. Notably, the patient with LRBA deficiency (01/016) showed the highest individual community turnover across all timepoints, consistent with the more severe baseline disease phenotype and greater immunological reconstitution potential reported for this genotype. Similar slow, progressive microbiome changes under immunomodulatory therapy have been described in rheumatoid arthritis reinforcing that re-establishment of a healthy commensal community following chronic immune dysregulation is inherently gradual (12,33).

At the taxon level, abatacept treatment was associated with two opposing trajectories. Disease-enriched taxa - notably *Veillonella* (strongest signal), *Lacrimispora, GGB3612*, and *Parasutterella* - showed progressive decreases consistent with partial normalization of the disease-associated microbiome. Depleted taxa at baseline – principally *Collinsella* (sustained increase across all timepoints) and *Adlercreutzia* (gradual increase) – showed treatment-associated recovery. *Collinsella* recovery is of particular interest: this taxon increases in parallel with clinical remission following fecal microbiota transplantation for refractory ICI-related colitis (34), mirroring the gut phenotype of CTLA-4/LRBA insufficiency. The transient nominal decrease in *Faecalibacterium* at month 6, despite no baseline depletion, may reflect a temporary perturbation during peak community remodeling, consistent with the known sensitivity of this obligate anaerobe to niche perturbation during microbiome restructuring. Its partial recovery trajectory toward month 12 supports a model of progressive re-equilibration rather than treatment-induced loss of butyrate-producing capacity.

On a functional level, we observed a progressive reduction in microbial folate and purine biosynthesis over the treatment course. These pathways are biochemically connected through the role of tetrahydrofolate as the obligate one-carbon donor for de-novo purine ring synthesis (35). Gut microbial folate biosynthetic capacity may contribute to intestinal immune tolerance, as folate availability is required for maintenance of colonic Foxp3 regulatory T cells, while microbiota-derived purine nucleosides—particularly inosine—can modulate T-cell responses through adenosine A2A receptor signaling (36–39). The decline in these pathways under abatacept may reflect a diminished microbiome-level compensatory demand for Treg support as the primary CTLA-4 defect is pharmacologically addressed and may simultaneously reduce the availability of immunomodulatory purine metabolites at the mucosal surface. Integration with longitudinal metabolomics and Treg phenotyping is required to resolve the directionality of these changes.

Organ-involvement restricted analyses identified the gut-involvement as having the most coherent microbiome correlates, with depletion of *Alistipes, Ruminococcus*, and *Lachnoclostridium* and enrichment of *Escherichia* and *Veillonella* tracking with higher gut severity scores — a pattern fully concordant with the patient-vs-control baseline signature. *Lachnoclostridium* showed a strong signal in the cytopenia domain and showed nominal associations with gut and lymphoproliferation scores, making it a candidate for a marker of overall disease activity. A moderate negative correlation between community turnover and overall CHAI score improvement at month 6 is hypothesis-generating and warrants prospective validation with larger sample sizes.

The primary limitation of this study is the relatively small sample size, unavoidable considering the rarity of both diseases, leading to restricted statistical power at the taxon level. In addition, the incomplete time-series coverage further temper interpretation. Nonetheless, this study provides the first longitudinal metagenomic framework for treatment monitoring in monogenic immune dysregulation, identifies *Veillonella* suppression and *Collinsella* recovery as candidate microbiome biomarkers of treatment response, and establishes the folate–purine metabolic axis as a mechanistic target for future investigation. Larger prospective cohorts with integrated metabolomics, mucosal immunophenotyping is needed to translate these exploratory, hypothesis-generating results into clinically actionable findings.

## Ethics and clinical trial registration information

Current research was approved by the Ethics Committee of the University of Freiburg (Protocol no. 466/18).

## Acknowledgments

We acknowledge the support of Nils Kolb and Sven Kolb for helping metadata curation. In addition, we thank for the help of the whole ABACHAI trial team, as well the participating patients.

## Author Contributions

BG, MP and MK planned the study; BG and MK recruited and treated patients. BZ and PM performed experiments. MK performed the statistical analyses and wrote the first draft. All authors contributed to the writing of the manuscript and critically revised and approved the final version.

## Declaration of generative AI and AI-assisted technologies in the manuscript preparation process

During the preparation of this work the authors used Perplexity for literature search, text editing as well as support during the post-processing analysis; they then reviewed and edited the text and code and take full responsibility for the content.

