## Supplementary Tables 1-6 for "Longitudinal Gut Microbiome Dynamics in Human CTLA-4 pathway Deficiencies: A One-Year Interventional Metagenomic Study"

### Supplementary Table 1: Differential Abundance Analysis (MaAsLin3) at genus level, Healthy controls vs. Patients at baseline

| feature | metadata | value | name | coef | null_hypothesis | stderr | pval_individual | qval_individual | pval_joint | qval_joint | error | model | N | N_not_zero |
|---|---|---|---|---|---|---|---|---|---|---|---|---|---|---|
| **GGB9760** | group | patient | grouppatient | -6.42019066431558 | 0.420724300002535 | 1.26832230455916 | 0.000196908332843582 | 0.0274648753932116 | 0.000196908332843582 | 0.0275671665981014 | NA | abundance | 26 | 14 |
| **Lacrimispora** | group | patient | grouppatient | 3.5547549211292 | 0.420724300002535 | 0.654688746821889 | 0.000224203064434381 | 0.0274648753932116 | 0.00044835586185466 | 0.0313849103298262 | NA | abundance | 26 | 24 |
| **GGB3612** | group | patient | grouppatient | 5.20009650379714 | 0.420724300002535 | 1.21228882606003 | 0.0027628615718156 | 0.132636594730824 | 0.00551808973956618 | 0.151338487744683 | NA | abundance | 26 | 13 |
| **Oscillibacter** | group | patient | grouppatient | -2.6534775461037 | 0.420724300002535 | 0.869121711691404 | 0.00324597900411083 | 0.132636594730824 | 0.00648142162852653 | 0.151338487744683 | NA | abundance | 26 | 21 |
| **Candidatus_Cibionibacter** | group | patient | grouppatient | -3.70414328264262 | 0 | 1.18930949481236 | 0.00184236549392622 | 0.132636594730824 | 0.00368133667723923 | 0.151338487744683 | NA | prevalence | 26 | 18 |
| **GGB9699** | group | patient | grouppatient | -3.07925473410008 | 0 | 1.04622200395695 | 0.00324824313626507 | 0.132636594730824 | 0.00648593518905785 | 0.151338487744683 | NA | prevalence | 26 | 17 |
| **Ruminococcus** | group | patient | grouppatient | -3.69872485423468 | 0.420724300002535 | 1.30714718901883 | 0.00569335501773827 | 0.172417391959094 | 0.0113542957441185 | 0.198700175522074 | NA | abundance | 26 | 23 |
| **Dysosmobacter** | group | patient | grouppatient | -3.17544833094058 | 0 | 1.13666845661122 | 0.00521176322453495 | 0.172417391959094 | 0.0103963639731613 | 0.198700175522074 | NA | prevalence | 26 | 19 |
| **Simiaoa** | group | patient | grouppatient | -2.65428322481268 | 0 | 0.984867192958462 | 0.00703744456975892 | 0.172417391959094 | 0.0140253635134454 | 0.218172321320262 | NA | prevalence | 26 | 16 |
| **Eggerthella** | group | patient | grouppatient | 3.88771113347635 | 0.420724300002535 | 1.14080797704353 | 0.00896237721163107 | 0.197750937262674 | 0.0178444302179786 | 0.245352627347026 | NA | abundance | 26 | 19 |
| **Adlercreutzia** | group | patient | grouppatient | -2.55055978239855 | 0 | 0.985971074893479 | 0.00968576019245749 | 0.197750937262674 | 0.0192777064344092 | 0.245352627347026 | NA | prevalence | 26 | 18 |
| **Eubacteriaceae_unclassified** | group | patient | grouppatient | -2.72476670777253 | 0 | 1.1170540641457 | 0.0147180528511501 | 0.242013488881385 | 0.029219484622571 | 0.269663801493871 | NA | prevalence | 26 | 20 |
| **Odoribacter** | group | patient | grouppatient | -4.63578753051609 | 0 | 1.89549614575292 | 0.0144577034389776 | 0.242013488881385 | 0.0287063816892257 | 0.269663801493871 | NA | prevalence | 26 | 20 |
| **Oliverpabstia** | group | patient | grouppatient | -2.3191201979155 | 0 | 0.951700375118188 | 0.014817152380493 | 0.242013488881385 | 0.0294147567563192 | 0.269663801493871 | NA | prevalence | 26 | 15 |
| **GGB9642** | group | patient | grouppatient | -3.52317833196253 | 0.420724300002535 | 1.21912635209528 | 0.0251585547457429 | 0.256070713757921 | 0.0496841566145913 | 0.269663801493871 | NA | abundance | 26 | 7 |
| **Lactobacillus** | group | patient | grouppatient | 6.81726470126762 | 0.420724300002535 | 2.4459767055852 | 0.0264952877648046 | 0.256070713757921 | 0.0522885752558694 | 0.269663801493871 | NA | abundance | 26 | 12 |
| **Oscillospiraceae_unclassified** | group | patient | grouppatient | -2.70360555784683 | 0.420724300002535 | 1.22786476789743 | 0.022237147704308 | 0.256070713757921 | 0.0439798046705927 | 0.269663801493871 | NA | abundance | 26 | 22 |
| **Parasutterella** | group | patient | grouppatient | 4.72143141056262 | 0.420724300002535 | 1.68381364578054 | 0.0220207318987902 | 0.256070713757921 | 0.043556551164222 | 0.269663801493871 | NA | abundance | 26 | 19 |
| **Prevotella** | group | patient | grouppatient | -6.26565552695906 | 0.420724300002535 | 2.37364226366548 | 0.0172312362863081 | 0.256070713757921 | 0.0341655570686616 | 0.269663801493871 | NA | abundance | 26 | 13 |
| **Streptococcus** | group | patient | grouppatient | 2.83676462170468 | 0.420724300002535 | 0.984596991035946 | 0.026359833512628 | 0.256070713757921 | 0.026359833512628 | 0.269663801493871 | NA | abundance | 26 | 26 |
| **Faecalibacillus** | group | patient | grouppatient | -2.09987815923056 | 0 | 0.963293132684478 | 0.0292652244294767 | 0.256070713757921 | 0.0576739954980458 | 0.269663801493871 | NA | prevalence | 26 | 19 |
| **Gemmiger** | group | patient | grouppatient | -2.09987815923056 | 0 | 0.963293132684478 | 0.0292652244294764 | 0.256070713757921 | 0.0576739954980453 | 0.269663801493871 | NA | prevalence | 26 | 19 |
| **GGB34797** | group | patient | grouppatient | -2.03320255166375 | 0 | 0.931898824816764 | 0.0291254299541916 | 0.256070713757921 | 0.0574025692383668 | 0.269663801493871 | NA | prevalence | 26 | 14 |
| **GGB9730** | group | patient | grouppatient | -2.52358373750615 | 0 | 1.1050512259922 | 0.0223903254751718 | 0.256070713757921 | 0.0442793242754595 | 0.269663801493871 | NA | prevalence | 26 | 12 |
| **Intestinibacter** | group | patient | grouppatient | -2.09987815923056 | 0 | 0.963293132684478 | 0.0292652244294766 | 0.256070713757921 | 0.0576739954980455 | 0.269663801493871 | NA | prevalence | 26 | 19 |
| **Lentihominibacter** | group | patient | grouppatient | -4.18510590734803 | 0 | 1.88379941621544 | 0.0263082988152097 | 0.256070713757921 | 0.051924471043869 | 0.269663801493871 | NA | prevalence | 26 | 21 |
| **Sutterella** | group | patient | grouppatient | -2.52358373750615 | 0 | 1.1050512259922 | 0.0223903254751718 | 0.256070713757921 | 0.0442793242754595 | 0.269663801493871 | NA | prevalence | 26 | 12 |
| **GGB3740** | group | patient | grouppatient | 4.9389302724262 | 0.420724300002535 | 1.60472890590273 | 0.039378852664427 | 0.26577865541441 | 0.0772070112916873 | 0.269663801493871 | NA | abundance | 26 | 7 |
| **GGB9614** | group | patient | grouppatient | 3.11780148810185 | 0.420724300002535 | 1.19567304520784 | 0.0418634585981248 | 0.26577865541441 | 0.0819743680304527 | 0.269663801493871 | NA | abundance | 26 | 19 |
| **Lachnospira** | group | patient | grouppatient | 1.75566083493182 | 0.420724300002535 | 0.553984377940127 | 0.0423076226986203 | 0.26577865541441 | 0.0828253104588318 | 0.269663801493871 | NA | abundance | 26 | 24 |
| **Veillonella** | group | patient | grouppatient | 4.50439577288188 | 0.420724300002535 | 1.8028273139997 | 0.040929893583504 | 0.26577865541441 | 0.080184530978251 | 0.269663801493871 | NA | abundance | 26 | 17 |
| **Collinsella** | group | patient | grouppatient | -2.29513865564977 | 0 | 1.1170540641457 | 0.0399142483432504 | 0.26577865541441 | 0.0782353494656941 | 0.269663801493871 | NA | prevalence | 26 | 21 |
| **Eubacterium** | group | patient | grouppatient | -2.29513865564977 | 0 | 1.1170540641457 | 0.0399142483432503 | 0.26577865541441 | 0.078235349465694 | 0.269663801493871 | NA | prevalence | 26 | 21 |
| **GGB9758** | group | patient | grouppatient | -2.28441402400212 | 0 | 1.09914307434828 | 0.0376762957124711 | 0.26577865541441 | 0.0739330881663286 | 0.269663801493871 | NA | prevalence | 26 | 11 |
| **GGB9770** | group | patient | grouppatient | -2.28441402400212 | 0 | 1.09914307434828 | 0.0376762957124714 | 0.26577865541441 | 0.0739330881663291 | 0.269663801493871 | NA | prevalence | 26 | 11 |

| | | | | | | | | | | | | | | |
|---|---|---|---|---|---|---|---|---|---|---|---|---|---|---|
| **Limosilactobacillus** | group | patient | grouppatient | 2.29513865564977 | 0 | 1.1170540641457 | 0.0399142483432502 | 0.26577865541441 | 0.0782353494656938 | 0.269663801493871 | NA | prevalence | 26 | 5 |
| **Oscillibacter** | group | patient | grouppatient | -2.29513865564977 | 0 | 1.1170540641457 | 0.0399142483432501 | 0.26577865541441 | 0.00648142162852653 | 0.151338487744683 | NA | prevalence | 26 | 21 |
| **Ruthenibacterium** | group | patient | grouppatient | -2.29513865564977 | 0 | 1.1170540641457 | 0.0399142483432501 | 0.26577865541441 | 0.0782353494656936 | 0.269663801493871 | NA | prevalence | 26 | 21 |
| **Oscillospiraceae_unclassified** | group | patient | grouppatient | -3.75547785522527 | 0 | 1.88379941621544 | 0.0461995396279828 | 0.282972180221394 | 0.0439798046705927 | 0.269663801493871 | NA | prevalence | 26 | 22 |
| **Flavonifractor** | group | patient | grouppatient | 2.80105850521711 | 0.420724300002535 | 1.1278071629532 | 0.0532163436664443 | 0.313949064624399 | 0.103600708099664 | 0.290081982679058 | NA | abundance | 26 | 22 |
| **Lentihominibacter** | group | patient | grouppatient | -1.84764889371345 | 0.420724300002535 | 1.07028193041846 | 0.053819839649897 | 0.313949064624399 | 0.051924471043869 | 0.269663801493871 | NA | abundance | 26 | 21 |
| **Agathobaculum** | group | patient | grouppatient | 2.17537661488372 | 0.420724300002535 | 0.971044575008725 | 0.0954280145469291 | 0.326409213144366 | 0.181749523133489 | 0.322309761189383 | NA | abundance | 26 | 24 |
| **Alistipes** | group | patient | grouppatient | -2.30447924764639 | 0.420724300002535 | 1.36277925241755 | 0.0625075260918908 | 0.326409213144366 | 0.121107861365653 | 0.302769653414133 | NA | abundance | 26 | 25 |
| **Clostridiaceae_unclassified** | group | patient | grouppatient | -0.815632222638034 | 0.420724300002535 | 0.619512926462653 | 0.080605028601695 | 0.326409213144366 | 0.15471288656751 | 0.321978487862176 | NA | abundance | 26 | 25 |
| **Coprococcus** | group | patient | grouppatient | -0.643602361600731 | 0.420724300002535 | 0.513129410254453 | 0.0794687443391423 | 0.326409213144366 | 0.152622207351445 | 0.321978487862176 | NA | abundance | 26 | 25 |
| **Faecalicatena** | group | patient | grouppatient | -2.27035908106739 | 0.420724300002535 | 1.37233863473138 | 0.0687989699644462 | 0.326409213144366 | 0.132864641660724 | 0.321907204865078 | NA | abundance | 26 | 22 |
| **Lachnospiraceae_unclassified** | group | patient | grouppatient | -1.11088083174158 | 0.420724300002535 | 0.789373427791927 | 0.0808132604466696 | 0.326409213144366 | 0.155095737829318 | 0.321978487862176 | NA | abundance | 26 | 23 |
| **Limosilactobacillus** | group | patient | grouppatient | 7.80509870657096 | 0.420724300002535 | 2.96395184716104 | 0.0892610222102559 | 0.326409213144366 | 0.0782353494656938 | 0.269663801493871 | NA | abundance | 26 | 5 |
| **Megasphaera** | group | patient | grouppatient | -4.39959560975597 | 0.420724300002535 | 2.21192224827419 | 0.0827701474489697 | 0.326409213144366 | 0.158689397589215 | 0.321978487862176 | NA | abundance | 26 | 7 |
| **Paraprevotella** | group | patient | grouppatient | 2.17742125238868 | 0.420724300002535 | 0.825929114668946 | 0.0900816619548857 | 0.326409213144366 | 0.172048618089217 | 0.322309761189383 | NA | abundance | 26 | 8 |
| **Barnesiella** | group | patient | grouppatient | -1.67490664994316 | 0 | 0.89627928999797 | 0.0616599760865849 | 0.326409213144366 | 0.119517999522172 | 0.302769653414133 | NA | prevalence | 26 | 18 |
| **Butyricimonas** | group | patient | grouppatient | -1.50450759996222 | 0 | 0.863709383217398 | 0.0815234667916348 | 0.326409213144366 | 0.156400857945543 | 0.321978487862176 | NA | prevalence | 26 | 15 |
| **Escherichia** | group | patient | grouppatient | 1.53738932031307 | 0 | 0.91263569520397 | 0.0920735505720502 | 0.326409213144366 | 0.175669562429156 | 0.322309761189383 | NA | prevalence | 26 | 14 |
| **GGB3005** | group | patient | grouppatient | -4.08370304422165 | 0 | 2.47237415991193 | 0.0985889051946244 | 0.326409213144366 | 0.0985889051946244 | 0.281682586270357 | NA | prevalence | 26 | 9 |
| **GGB3033** | group | patient | grouppatient | -4.08370304422165 | 0 | 2.47237415991193 | 0.0985889051946245 | 0.326409213144366 | 0.0985889051946245 | 0.281682586270357 | NA | prevalence | 26 | 9 |
| **GGB33469** | group | patient | grouppatient | -4.31349060066882 | 0 | 2.47358895475108 | 0.0811907139073936 | 0.326409213144366 | 0.0811907139073936 | 0.269663801493871 | NA | prevalence | 26 | 10 |
| **GGB33512** | group | patient | grouppatient | -4.31349060066882 | 0 | 2.47358895475109 | 0.0811907139073935 | 0.326409213144366 | 0.0811907139073935 | 0.269663801493871 | NA | prevalence | 26 | 10 |
| **GGB3653** | group | patient | grouppatient | -1.50450759996222 | 0 | 0.863709383217398 | 0.08152346679163 5 | 0.326409213144366 | 0.156400857945543 | 0.321978487862176 | NA | prevalence | 26 | 15 |
| **GGB9345** | group | patient | grouppatient | -4.08370304422165 | 0 | 2.47237415991193 | 0.0985889051946245 | 0.326409213144366 | 0.0985889051946245 | 0.281682586270357 | NA | prevalence | 26 | 9 |
| **GGB9453** | group | patient | grouppatient | -4.08370304422165 | 0 | 2.47237415991193 | 0.0985889051946243 | 0.326409213144366 | 0.0985889051946243 | 0.281682586270357 | NA | prevalence | 26 | 9 |
| **GGB9509** | group | patient | grouppatient | -4.55266031417286 | 0 | 2.47621991217464 | 0.0659810329776004 | 0.326409213144366 | 0.0659810329776004 | 0.269663801493871 | NA | prevalence | 26 | 11 |
| **GGB9522** | group | patient | grouppatient | -4.55266031417287 | 0 | 2.47621991217465 | 0.0659810329776006 | 0.326409213144366 | 0.0659810329776006 | 0.269663801493871 | NA | prevalence | 26 | 11 |
| **GGB9602** | group | patient | grouppatient | -4.08370304422164 | 0 | 2.47237415991193 | 0.0985889051946248 | 0.326409213144366 | 0.0985889051946248 | 0.281682586270357 | NA | prevalence | 26 | 9 |
| **GGB9608** | group | patient | grouppatient | -4.55266031417286 | 0 | 2.47621991217464 | 0.0659810329776004 | 0.326409213144366 | 0.0659810329776004 | 0.269663801493871 | NA | prevalence | 26 | 11 |
| **GGB9635** | group | patient | grouppatient | -1.82780414168496 | 0 | 1.09640647935638 | 0.0954972602661773 | 0.326409213144366 | 0.181874793814009 | 0.322309761189383 | NA | prevalence | 26 | 9 |
| **GGB9708** | group | patient | grouppatient | -4.55266031417286 | 0 | 2.47621991217464 | 0.0659810329776003 | 0.326409213144366 | 0.0659810329776003 | 0.269663801493871 | NA | prevalence | 26 | 11 |
| **Lachnospiraceae_unclassified** | group | patient | grouppatient | -3.30479623205722 | 0 | 1.89549614575292 | 0.0812464338097808 | 0.326409213144366 | 0.155095737829318 | 0.321978487862176 | NA | prevalence | 26 | 23 |
| **Lawsonibacter** | group | patient | grouppatient | -3.30479623205722 | 0 | 1.89549614575292 | 0.0812464338097808 | 0.326409213144366 | 0.155891884612755 | 0.321978487862176 | NA | prevalence | 26 | 23 |
| **Mogibacterium** | group | patient | grouppatient | -1.82780414168496 | 0 | 1.09640647935638 | 0.0954972602661775 | 0.326409213144366 | 0.181874793814009 | 0.322309761189383 | NA | prevalence | 26 | 9 |
| **Senegalimassilia** | group | patient | grouppatient | -4.08370304422165 | 0 | 2.47237415991193 | 0.0985889051946242 | 0.326409213144366 | 0.0985889051946242 | 0.281682586270357 | NA | prevalence | 26 | 9 |
| **Slackia** | group | patient | grouppatient | -1.53738932031307 | 0 | 0.91263569520397 | 0.09207355057205 | 0.326409213144366 | 0.175669562429156 | 0.322309761189383 | NA | prevalence | 26 | 12 |
| **Turicibacter** | group | patient | grouppatient | -1.53738932031307 | 0 | 0.91263569520397 | 0.09207355057205 | 0.326409213144366 | 0.175669562429156 | 0.322309761189383 | NA | prevalence | 26 | 12 |
| **Veillonella** | group | patient | grouppatient | 4.08370304422165 | 0 | 2.47237415991193 | 0.0985889051946248 | 0.326409213144366 | 0.080184530978251 | 0.269663801493871 | NA | prevalence | 26 | 17 |

| | | | | | | | | | | | | | | |
|---|---|---|---|---|---|---|---|---|---|---|---|---|---|---|
| **Enterocloster** | group | patient | grouppatient | 2.89899628345633 | 0.420724300002535 | 1.42274637320781 | 0.105818838842147 | 0.332379686106743 | 0.200440051030393 | 0.342214721271403 | NA | abundance | 26 | 18 |
| **Brotolimicola** | group | patient | grouppatient | -1.84445703248171 | 0 | 1.13666845661122 | 0.104656349736653 | 0.332379686106743 | 0.198359747933105 | 0.342214721271403 | NA | prevalence | 26 | 22 |
| **Faecalicatena** | group | patient | grouppatient | -1.84445703248172 | 0 | 1.13666845661122 | 0.104656349736653 | 0.332379686106743 | 0.132864641660724 | 0.321907204865078 | NA | prevalence | 26 | 22 |
| **Romboutsia** | group | patient | grouppatient | -1.84445703248172 | 0 | 1.13666845661122 | 0.104656349736653 | 0.332379686106743 | 0.198359747933105 | 0.342214721271403 | NA | prevalence | 26 | 22 |
| **GGB4552** | group | patient | grouppatient | -2.26332014168628 | 0.420724300002535 | 1.39778997815678 | 0.117214776372439 | 0.343411231004011 | 0.220690248944837 | 0.359263195956711 | NA | abundance | 26 | 7 |
| **GGB9715** | group | patient | grouppatient | -2.22261690791666 | 0.420724300002535 | 1.54565448738077 | 0.11333258309392 | 0.343411231004011 | 0.213820891797101 | 0.352175586489342 | NA | abundance | 26 | 16 |
| **Ellagibacter** | group | patient | grouppatient | -3.85688071835166 | 0 | 2.47237415991194 | 0.11876207683137 | 0.343411231004011 | 0.11876207683137 | 0.302769653414133 | NA | prevalence | 26 | 8 |
| **GGB3304** | group | patient | grouppatient | -3.85688071835166 | 0 | 2.47237415991193 | 0.118762076831369 | 0.343411231004011 | 0.118762076831369 | 0.302769653414133 | NA | prevalence | 26 | 8 |
| **Hydrogenoanaerobacterium** | group | patient | grouppatient | -1.33974362304598 | 0 | 0.859701565610698 | 0.119142671980983 | 0.343411231004011 | 0.224090367675199 | 0.360605189362389 | NA | prevalence | 26 | 17 |
| **Methanobrevibacter** | group | patient | grouppatient | -3.85688071835166 | 0 | 2.47237415991193 | 0.11876207683137 | 0.343411231004011 | 0.11876207683137 | 0.302769653414133 | NA | prevalence | 26 | 8 |
| **Monoglobus** | group | patient | grouppatient | -3.85688071835166 | 0 | 2.47237415991193 | 0.11876207683137 | 0.343411231004011 | 0.11876207683137 | 0.302769653414133 | NA | prevalence | 26 | 8 |
| **Brotolimicola** | group | patient | grouppatient | 2.45797548739637 | 0.420724300002535 | 1.2519390885117 | 0.127197477967725 | 0.357848071344328 | 0.198359747933105 | 0.342214721271403 | NA | abundance | 26 | 22 |
| **Collinsella** | group | patient | grouppatient | -0.556745302489149 | 0.420724300002535 | 0.553979689012085 | 0.129044762109661 | 0.357848071344328 | 0.0782353494656941 | 0.269663801493871 | NA | abundance | 26 | 21 |
| **Erysipelatoclostridium** | group | patient | grouppatient | 1.8629270584852 | 0.420724300002535 | 0.891798884817442 | 0.14127872627083 | 0.357848071344328 | 0.262597774044952 | 0.391103067726525 | NA | abundance | 26 | 18 |
| **GGB9480** | group | patient | grouppatient | -3.38316732211156 | 0.420724300002535 | 0.733194510606077 | 0.129664088180031 | 0.357848071344328 | 0.242515400596504 | 0.380598595198154 | NA | abundance | 26 | 3 |
| **Agathobaculum** | group | patient | grouppatient | -2.77610128035518 | 0 | 1.92752367891039 | 0.149798838437826 | 0.357848071344328 | 0.181749523133489 | 0.322309761189383 | NA | prevalence | 26 | 24 |
| **Anaerobutyricum** | group | patient | grouppatient | -2.77610128035518 | 0 | 1.92752367891039 | 0.149798838437826 | 0.357848071344328 | 0.277157984878331 | 0.393487153673676 | NA | prevalence | 26 | 24 |
| **Erysipelatoclostridium** | group | patient | grouppatient | 1.59801658523779 | 0 | 1.09914307434828 | 0.145980978232091 | 0.357848071344328 | 0.262597774044952 | 0.391103067726525 | NA | prevalence | 26 | 18 |
| **Eubacteriales_unclassified** | group | patient | grouppatient | -2.77610128035518 | 0 | 1.92752367891039 | 0.149798838437826 | 0.357848071344328 | 0.277157984878331 | 0.393487153673676 | NA | prevalence | 26 | 24 |
| **Evtepia** | group | patient | grouppatient | -1.59801658523779 | 0 | 1.09914307434828 | 0.145980978232091 | 0.357848071344328 | 0.270651510458584 | 0.393487153673676 | NA | prevalence | 26 | 8 |
| **GGB13489** | group | patient | grouppatient | -3.62709316190449 | 0 | 2.47358895475109 | 0.142558905011677 | 0.357848071344328 | 0.142558905011677 | 0.321907204865078 | NA | prevalence | 26 | 7 |
| **GGB87445** | group | patient | grouppatient | -3.62709316190449 | 0 | 2.47358895475108 | 0.142558905011677 | 0.357848071344328 | 0.142558905011677 | 0.321907204865078 | NA | prevalence | 26 | 7 |
| **GGB9342** | group | patient | grouppatient | -3.62709316190449 | 0 | 2.47358895475108 | 0.142558905011677 | 0.357848071344328 | 0.142558905011677 | 0.321907204865078 | NA | prevalence | 26 | 7 |
| **GGB9627** | group | patient | grouppatient | -1.30760176386589 | 0 | 0.909337991204204 | 0.150442250401901 | 0.357848071344328 | 0.278251630097814 | 0.393487153673676 | NA | prevalence | 26 | 11 |
| **GGB9634** | group | patient | grouppatient | -3.62709316190449 | 0 | 2.47358895475109 | 0.142558905011677 | 0.357848071344328 | 0.142558905011677 | 0.321907204865078 | NA | prevalence | 26 | 7 |
| **GGB9747** | group | patient | grouppatient | -3.62709316190449 | 0 | 2.47358895475109 | 0.142558905011677 | 0.357848071344328 | 0.142558905011677 | 0.321907204865078 | NA | prevalence | 26 | 7 |
| **Lachnospira** | group | patient | grouppatient | -2.77610128035518 | 0 | 1.92752367891039 | 0.149798838437826 | 0.357848071344328 | 0.0828253104588318 | 0.269663801493871 | NA | prevalence | 26 | 24 |
| **Lactobacillus** | group | patient | grouppatient | 1.24786408211557 | 0 | 0.850579276318447 | 0.142355497281933 | 0.357848071344328 | 0.0522885752558694 | 0.269663801493871 | NA | prevalence | 26 | 12 |
| **Phocaeicola** | group | patient | grouppatient | -2.77610128035518 | 0 | 1.92752367891039 | 0.149798838437826 | 0.357848071344328 | 0.27715798487833 | 0.393487153673676 | NA | prevalence | 26 | 24 |
| **GGB9614** | group | patient | grouppatient | -1.2452785978204 | 0 | 0.89627928999797 | 0.164715167717928 | 0.384335391341831 | 0.0819743680304527 | 0.269663801493871 | NA | prevalence | 26 | 19 |
| **Megasphaera** | group | patient | grouppatient | 1.2452785978204 | 0 | 0.89627928999797 | 0.164715167717927 | 0.384335391341831 | 0.158689397589215 | 0.321978487862176 | NA | prevalence | 26 | 7 |
| **GGB4567** | group | patient | grouppatient | -3.38792344840044 | 0 | 2.47621991217465 | 0.171254616937499 | 0.388494269904513 | 0.171254616937499 | 0.322309761189383 | NA | prevalence | 26 | 6 |
| **GGB6606** | group | patient | grouppatient | -3.38792344840044 | 0 | 2.47621991217465 | 0.1712546169375 | 0.388494269904513 | 0.1712546169375 | 0.322309761189383 | NA | prevalence | 26 | 6 |
| **GGB9176** | group | patient | grouppatient | -3.38792344840044 | 0 | 2.47621991217465 | 0.171254616937499 | 0.388494269904513 | 0.171254616937499 | 0.322309761189383 | NA | prevalence | 26 | 6 |
| **Evtepia** | group | patient | grouppatient | -0.493787788034658 | 0.420724300002535 | 0.534553114669812 | 0.177133089497293 | 0.398143182815016 | 0.270651510458584 | 0.393487153673676 | NA | abundance | 26 | 8 |
| **Bilophila** | group | patient | grouppatient | 1.95475617224214 | 0.420724300002535 | 1.12867510725082 | 0.207000424054682 | 0.45185666623089 | 0.371151672550545 | 0.514467664921548 | NA | abundance | 26 | 15 |
| **GGB9261** | group | patient | grouppatient | -3.13127993055332 | 0 | 2.48076076833312 | 0.206867593166369 | 0.45185666623089 | 0.206867593166369 | 0.344779321943948 | NA | prevalence | 26 | 5 |
| **GGB9524** | group | patient | grouppatient | -3.13127993055332 | 0 | 2.48076076833312 | 0.206867593166369 | 0.45185666623089 | 0.206867593166369 | 0.344779321943948 | NA | prevalence | 26 | 5 |

| | | | | | | | | | | | | | | |
|---|---|---|---|---|---|---|---|---|---|---|---|---|---|---|
| **GGB9715** | group | patient | grouppatient | -1.05382597679423 | 0 | 0.837728236129266 | 0.208407360343227 | 0.45185666623089 | 0.213820891797101 | 0.352175586489342 | NA | prevalence | 26 | 16 |
| **Dialister** | group | patient | grouppatient | -1.30980320793351 | 0.420724300002535 | 1.31353870983656 | 0.217688593482972 | 0.458215340507567 | 0.387988863233349 | 0.532533733849695 | NA | abundance | 26 | 15 |
| **Romboutsia** | group | patient | grouppatient | -0.804171303545303 | 0.420724300002535 | 0.925859886106991 | 0.216862540973495 | 0.458215340507567 | 0.198359747933105 | 0.342214721271403 | NA | abundance | 26 | 22 |
| **GGB4552** | group | patient | grouppatient | -1.35884687173375 | 0 | 1.1050512259922 | 0.218821203426062 | 0.458215340507567 | 0.220690248944837 | 0.359263195956711 | NA | prevalence | 26 | 7 |
| **GGB9642** | group | patient | grouppatient | -1.35884687173375 | 0 | 1.1050512259922 | 0.218821203426062 | 0.458215340507567 | 0.0496841566145913 | 0.269663801493871 | NA | prevalence | 26 | 7 |
| **Akkermansia** | group | patient | grouppatient | 2.2094590461352 | 0.420724300002535 | 1.40095703265146 | 0.228384420520222 | 0.474187991758087 | 0.404609397504086 | 0.54995452087934 | NA | abundance | 26 | 16 |
| **Candidatus_Gastranaerophilale** | group | patient | grouppatient | 5.21880997617163 | 0.420724300002535 | 3.69792686483842 | 0.235918561768244 | 0.481667063610166 | 0.416179555749692 | 0.554906074332923 | NA | abundance | 26 | 9 |
| **Segatella** | group | patient | grouppatient | -1.0807794379959 | 0 | 0.909337991204204 | 0.234622955164917 | 0.481667063610166 | 0.414197979239516 | 0.554906074332923 | NA | prevalence | 26 | 10 |
| **GGB1495** | group | patient | grouppatient | -2.84536228430117 | 0 | 2.48826692682863 | 0.252826209667345 | 0.495539370947996 | 0.252826209667345 | 0.380598595198154 | NA | prevalence | 26 | 4 |
| **GGB6544** | group | patient | grouppatient | -2.84536228430117 | 0 | 2.48826692682863 | 0.252826209667345 | 0.495539370947996 | 0.252826209667345 | 0.380598595198154 | NA | prevalence | 26 | 4 |
| **GGB9712** | group | patient | grouppatient | -2.84536228430117 | 0 | 2.48826692682862 | 0.252826209667345 | 0.495539370947996 | 0.252826209667345 | 0.380598595198154 | NA | prevalence | 26 | 4 |
| **GGB9737** | group | patient | grouppatient | -2.84536228430117 | 0 | 2.48826692682863 | 0.252826209667345 | 0.495539370947996 | 0.252826209667345 | 0.380598595198154 | NA | prevalence | 26 | 4 |
| **Parolsenella** | group | patient | grouppatient | -2.84536228430117 | 0 | 2.48826692682863 | 0.252826209667345 | 0.495539370947996 | 0.252826209667345 | 0.380598595198154 | NA | prevalence | 26 | 4 |
| **Parabacteroides** | group | patient | grouppatient | -1.31576208077968 | 0 | 1.18930949481236 | 0.268586146203614 | 0.518138628503035 | 0.465033774474719 | 0.608455405854772 | NA | prevalence | 26 | 23 |
| **Ruminococcus** | group | patient | grouppatient | -1.31576208077968 | 0 | 1.18930949481236 | 0.268586146203614 | 0.518138628503035 | 0.0113542957441185 | 0.198700175522074 | NA | prevalence | 26 | 23 |
| **Phascolarctobacterium** | group | patient | grouppatient | 2.69901903690545 | 0.420724300002535 | 1.90242269265657 | 0.272906000339479 | 0.522359141274785 | 0.471334315657667 | 0.610988927704383 | NA | abundance | 26 | 9 |
| **Adlercreutzia** | group | patient | grouppatient | -1.5126917567602 | 0.420724300002535 | 1.73356019892278 | 0.284640296391111 | 0.532352824361259 | 0.0192777064344092 | 0.245352627347026 | NA | abundance | 26 | 18 |
| **GGB51647** | group | patient | grouppatient | -1.93570506443328 | 0.420724300002535 | 1.7935759040634 | 0.283016953937165 | 0.532352824361259 | 0.485935311658459 | 0.621432461557186 | NA | abundance | 26 | 5 |
| **Schaalia** | group | patient | grouppatient | -0.387494449715621 | 0.420724300002535 | 0.691705899180748 | 0.284645795882959 | 0.532352824361259 | 0.488268362652075 | 0.621432461557186 | NA | abundance | 26 | 25 |
| **Clostridium** | group | patient | grouppatient | -0.286885844118987 | 0.420724300002535 | 0.594504529032003 | 0.286946691101012 | 0.53259044939203 | 0.286946691101012 | 0.401725367541417 | NA | abundance | 26 | 26 |
| **Vescimonas** | group | patient | grouppatient | -0.910115570923219 | 0 | 0.859701565610698 | 0.289763176865759 | 0.533774273173766 | 0.49556365506418 | 0.625035240621489 | NA | prevalence | 26 | 18 |
| **GGB9635** | group | patient | grouppatient | -3.12477428892935 | 0.420724300002535 | 3.28546075430769 | 0.316498375821423 | 0.538297658014675 | 0.181874793814009 | 0.322309761189383 | NA | abundance | 26 | 9 |
| **Ruthenibacterium** | group | patient | grouppatient | 1.04699197451946 | 0.420724300002535 | 0.558012988517994 | 0.322309372364306 | 0.538297658014675 | 0.0782353494656936 | 0.269663801493871 | NA | abundance | 26 | 21 |
| **Alistipes** | group | patient | grouppatient | -2.02907657653555 | 0 | 2.02251633158107 | 0.315743338514994 | 0.538297658014675 | 0.121107861365653 | 0.302769653414133 | NA | prevalence | 26 | 25 |
| **Anaerostipes** | group | patient | grouppatient | -2.02907657653555 | 0 | 2.02251633158107 | 0.315743338514994 | 0.538297658014675 | 0.531792821213394 | 0.631975142887608 | NA | prevalence | 26 | 25 |
| **Blautia** | group | patient | grouppatient | -2.02907657653555 | 0 | 2.02251633158107 | 0.315743338514994 | 0.538297658014675 | 0.531792821213394 | 0.631975142887608 | NA | prevalence | 26 | 25 |
| **Clostridiaceae_unclassified** | group | patient | grouppatient | -2.02907657653555 | 0 | 2.02251633158107 | 0.315743338514994 | 0.538297658014675 | 0.15471288656751 | 0.321978487862176 | NA | prevalence | 26 | 25 |
| **Coprococcus** | group | patient | grouppatient | -2.02907657653555 | 0 | 2.02251633158107 | 0.315743338514994 | 0.538297658014675 | 0.152622207351445 | 0.321978487862176 | NA | prevalence | 26 | 25 |
| **Faecalibacterium** | group | patient | grouppatient | -2.02907657653555 | 0 | 2.02251633158107 | 0.315743338514994 | 0.538297658014675 | 0.531792821213394 | 0.631975142887608 | NA | prevalence | 26 | 25 |
| **Fusicatenibacter** | group | patient | grouppatient | -2.02907657653555 | 0 | 2.02251633158107 | 0.315743338514994 | 0.538297658014675 | 0.531792821213394 | 0.631975142887608 | NA | prevalence | 26 | 25 |
| **GGB3363** | group | patient | grouppatient | -1.1022033538871 | 0 | 1.11518922404327 | 0.322978594808805 | 0.538297658014675 | 0.54164201691294 | 0.631975142887608 | NA | prevalence | 26 | 6 |
| **GGB4566** | group | patient | grouppatient | -1.1022033538871 | 0 | 1.11518922404327 | 0.322978594808804 | 0.538297658014675 | 0.541642016912938 | 0.631975142887608 | NA | prevalence | 26 | 6 |
| **Mediterraneibacter** | group | patient | grouppatient | -2.02907657653555 | 0 | 2.02251633158107 | 0.315743338514994 | 0.538297658014675 | 0.531792821213394 | 0.631975142887608 | NA | prevalence | 26 | 25 |
| **Roseburia** | group | patient | grouppatient | -2.02907657653555 | 0 | 2.02251633158107 | 0.315743338514994 | 0.538297658014675 | 0.531792821213394 | 0.631975142887608 | NA | prevalence | 26 | 25 |
| **Schaalia** | group | patient | grouppatient | -2.02907657653555 | 0 | 2.02251633158107 | 0.315743338514995 | 0.538297658014675 | 0.488268362652075 | 0.621432461557186 | NA | prevalence | 26 | 25 |
| **Acidaminococcus** | group | patient | grouppatient | 2.28170933695969 | 0.420724300002535 | 1.61429380885782 | 0.336561908873176 | 0.551454730705003 | 0.559849899241996 | 0.644068642287184 | NA | abundance | 26 | 5 |
| **Dorea** | group | patient | grouppatient | 0.041515226786598 | 0.420724300002535 | 0.273967342770806 | 0.337625345329594 | 0.551454730705003 | 0.56125981685026 | 0.644068642287184 | NA | abundance | 26 | 24 |
| **Bilophila** | group | patient | grouppatient | -0.797182458947579 | 0 | 0.82418432662414 | 0.333425033885632 | 0.551454730705003 | 0.371151672550545 | 0.514467664921548 | NA | prevalence | 26 | 15 |

| | | | | | | | | | | | | | | |
|---|---|---|---|---|---|---|---|---|---|---|---|---|---|---|
| **Anaerotignum** | group | patient | grouppatient | 1.72197732658862 | 0.420724300002535 | 1.31942151256514 | 0.344079084277828 | 0.558274010914357 | 0.569767752318188 | 0.64851614084997 | NA | abundance | 26 | 20 |
| **Candidatus_Gastranaerophilale** | group | patient | grouppatient | -0.850991881548737 | 0 | 0.91263569520397 | 0.35110133919456 | 0.558570312354982 | 0.416179555749692 | 0.554906074332923 | NA | prevalence | 26 | 9 |
| **GGB3175** | group | patient | grouppatient | -0.850991881548737 | 0 | 0.91263569520397 | 0.35110133919456 | 0.558570312354982 | 0.578930528004906 | 0.653631241295862 | NA | prevalence | 26 | 9 |
| **Phascolarctobacterium** | group | patient | grouppatient | -0.850991881548737 | 0 | 0.91263569520397 | 0.35110133919456 | 0.558570312354982 | 0.471334315657667 | 0.610988927704383 | NA | prevalence | 26 | 9 |
| **Intestinibacter** | group | patient | grouppatient | 2.05865219655839 | 0.420724300002535 | 1.75527616728363 | 0.366433201768119 | 0.571822512313307 | 0.0576739954980455 | 0.269663801493871 | NA | abundance | 26 | 19 |
| **Roseburia** | group | patient | grouppatient | 1.00113097812666 | 0.420724300002535 | 0.57380129353191 | 0.364814293078328 | 0.571822512313307 | 0.531792821213394 | 0.631975142887608 | NA | abundance | 26 | 25 |
| **Wujia** | group | patient | grouppatient | 1.78907298049032 | 0.420724300002535 | 1.26599373315929 | 0.365230259907002 | 0.571822512313307 | 0.597067377062267 | 0.66871546230974 | NA | abundance | 26 | 5 |
| **Barnesiella** | group | patient | grouppatient | -1.03706016400567 | 0.420724300002535 | 1.58393872065397 | 0.374761621671733 | 0.577462876160846 | 0.119517999522172 | 0.302769653414133 | NA | abundance | 26 | 18 |
| **Candidatus_Cibionibacter** | group | patient | grouppatient | -0.527734473933467 | 0.420724300002535 | 1.01142966042713 | 0.374344328323403 | 0.577462876160846 | 0.00368133667723923 | 0.151338487744683 | NA | abundance | 26 | 18 |
| **GGB3345** | group | patient | grouppatient | 1.70158956626409 | 0.420724300002535 | 1.30389293538672 | 0.37786271798279 | 0.578602286911147 | 0.612945202324238 | 0.681050224804709 | NA | abundance | 26 | 7 |
| **Anaerotignum** | group | patient | grouppatient | -0.794596974652409 | 0 | 0.920609508276334 | 0.388071196609035 | 0.590543125274618 | 0.569767752318188 | 0.64851614084997 | NA | prevalence | 26 | 20 |
| **GGB4599** | group | patient | grouppatient | 1.55219961700297 | 0.420724300002535 | 1.1842075174309 | 0.391419821707106 | 0.591962076038524 | 0.629630166588989 | 0.694080498602035 | NA | abundance | 26 | 7 |
| **GGB9770** | group | patient | grouppatient | -1.95029225444452 | 0.420724300002535 | 2.80617232475632 | 0.420701257719704 | 0.632342381235138 | 0.0739330881663291 | 0.269663801493871 | NA | abundance | 26 | 11 |
| **Lawsonibacter** | group | patient | grouppatient | -0.438832689475941 | 0.420724300002535 | 1.0358835479042 | 0.426123709546837 | 0.636587249018141 | 0.155891884612755 | 0.321978487862176 | NA | abundance | 26 | 23 |
| **Bacteroides** | group | patient | grouppatient | 1.01299505329844 | 0.420724300002535 | 0.69944413365188 | 0.432389037700174 | 0.642032207494197 | 0.432389037700174 | 0.571079861113437 | NA | abundance | 26 | 26 |
| **GGB4456** | group | patient | grouppatient | 1.95702362099776 | 0.420724300002535 | 1.91208810593072 | 0.446954869022299 | 0.651809183990853 | 0.694141083101858 | 0.75333140801752 | NA | abundance | 26 | 10 |
| **Holdemanella** | group | patient | grouppatient | -0.627645969004433 | 0.420724300002535 | 1.1715550171762 | 0.44419473914582 | 0.651809183990853 | 0.691080512006817 | 0.75333140801752 | NA | abundance | 26 | 5 |
| **Segatella** | group | patient | grouppatient | 1.92698782737413 | 0.420724300002535 | 1.86311416337925 | 0.445052876393469 | 0.651809183990853 | 0.414197979239516 | 0.554906074332923 | NA | abundance | 26 | 10 |
| **GGB5980** | group | patient | grouppatient | 3.2039348916618 | 0.420724300002535 | 3.00878228761304 | 0.453202191828281 | 0.657009094662301 | 0.701012156978604 | 0.754936169053881 | NA | abundance | 26 | 4 |
| **GGB9758** | group | patient | grouppatient | -1.9735856725323 | 0.420724300002535 | 3.1788703506248 | 0.470356135074572 | 0.666690053713385 | 0.0739330881663286 | 0.269663801493871 | NA | abundance | 26 | 11 |
| **Acidaminococcus** | group | patient | grouppatient | -0.816285707635357 | 0 | 1.13178849160409 | 0.47076481343843 | 0.666690053713385 | 0.559849899241996 | 0.644068642287184 | NA | prevalence | 26 | 5 |
| **GGB4266** | group | patient | grouppatient | -0.816285707635356 | 0 | 1.13178849160409 | 0.470764813438431 | 0.666690053713385 | 0.719910117305141 | 0.769369591013128 | NA | prevalence | 26 | 5 |
| **Holdemanella** | group | patient | grouppatient | -0.816285707635355 | 0 | 1.13178849160409 | 0.470764813438431 | 0.666690053713385 | 0.691080512006817 | 0.75333140801752 | NA | prevalence | 26 | 5 |
| **Butyricimonas** | group | patient | grouppatient | 1.24824309523476 | 0.420724300002535 | 1.17110559105058 | 0.499876683476233 | 0.699827356866726 | 0.156400857945543 | 0.321978487862176 | NA | abundance | 26 | 15 |
| **Vescimonas** | group | patient | grouppatient | -0.738834779995396 | 0.420724300002535 | 1.66220764938176 | 0.497943398926554 | 0.699827356866726 | 0.49556365506418 | 0.625035240621489 | NA | abundance | 26 | 18 |
| **Slackia** | group | patient | grouppatient | -0.936372548971961 | 0.420724300002535 | 1.9750575292809 | 0.508884883424308 | 0.702903524991588 | 0.175669562429156 | 0.322309761189383 | NA | abundance | 26 | 12 |
| **Dialister** | group | patient | grouppatient | 0.552084486294378 | 0 | 0.839317616406454 | 0.510680928361236 | 0.702903524991588 | 0.387988863233349 | 0.532533733849695 | NA | prevalence | 26 | 15 |
| **Paraprevotella** | group | patient | grouppatient | -0.6118221680447 | 0 | 0.919742695725005 | 0.505916095330552 | 0.702903524991588 | 0.172048618089217 | 0.322309761189383 | NA | prevalence | 26 | 8 |
| **GGB3256** | group | patient | grouppatient | 1.61177147358266 | 0.420724300002535 | 1.75530134080815 | 0.518619489909101 | 0.709842318590668 | 0.768272804504625 | 0.814834792656421 | NA | abundance | 26 | 10 |
| **Bifidobacterium** | group | patient | grouppatient | -0.199591422154018 | 0.420724300002535 | 0.976131207234789 | 0.54169297961795 | 0.733230828764628 | 0.54169297961795 | 0.631975142887608 | NA | abundance | 26 | 26 |
| **Blautia** | group | patient | grouppatient | 0.114382100862472 | 0.420724300002535 | 0.418812463802323 | 0.54133024046455 | 0.733230828764628 | 0.531792821213394 | 0.631975142887608 | NA | abundance | 26 | 25 |
| **GGB34797** | group | patient | grouppatient | -0.760839573185165 | 0.420724300002535 | 2.0264237381132 | 0.571284764749359 | 0.769037183316445 | 0.0574025692383668 | 0.269663801493871 | NA | abundance | 26 | 14 |
| **Escherichia** | group | patient | grouppatient | -0.679589072246972 | 0.420724300002535 | 1.95263751974222 | 0.584652947530087 | 0.78273208822334 | 0.175669562429156 | 0.322309761189383 | NA | abundance | 26 | 14 |
| **Eubacteriaceae_unclassified** | group | patient | grouppatient | -0.102675883729064 | 0.420724300002535 | 0.966208702585613 | 0.604295059445262 | 0.800282646292374 | 0.029219484622571 | 0.269663801493871 | NA | abundance | 26 | 20 |
| **Parasutterella** | group | patient | grouppatient | -0.459433947755229 | 0 | 0.885037899167607 | 0.603682519424122 | 0.800282646292374 | 0.043556551164222 | 0.269663801493871 | NA | prevalence | 26 | 19 |
| **Simiaoa** | group | patient | grouppatient | -0.703935678684179 | 0.420724300002535 | 2.23710367223316 | 0.622769181816313 | 0.820314244865574 | 0.0140253635134454 | 0.218172321320262 | NA | abundance | 26 | 16 |
| **Faecalibacillus** | group | patient | grouppatient | 1.04528173362626 | 0.420724300002535 | 1.6572493425593 | 0.712282504596615 | 0.829121845527177 | 0.0576739954980458 | 0.269663801493871 | NA | abundance | 26 | 19 |
| **GGB3267** | group | patient | grouppatient | 2.15487728182128 | 0.420724300002535 | 3.60406505049224 | 0.714060038392793 | 0.829121845527177 | 0.918238338356069 | 0.945245348307718 | NA | abundance | 26 | 3 |

| | | | | | | | | | | | | | | |
|---|---|---|---|---|---|---|---|---|---|---|---|---|---|---|
| **GGB36331** | group | patient | grouppatient | 1.45001910316709 | 0.420724300002535 | 2.36388301163274 | 0.685735511203016 | 0.829121845527177 | 0.90123783108117 | 0.945245348307718 | NA | abundance | 26 | 6 |
| **GGB4566** | group | patient | grouppatient | 1.83395791717125 | 0.420724300002535 | 3.50266214756691 | 0.706800451760536 | 0.829121845527177 | 0.541642016912938 | 0.631975142887608 | NA | abundance | 26 | 6 |
| **GGB9627** | group | patient | grouppatient | 1.52998815262004 | 0.420724300002535 | 2.68888749035772 | 0.689935899006845 | 0.829121845527177 | 0.278251630097814 | 0.393487153673676 | NA | abundance | 26 | 11 |
| **GGB9640** | group | patient | grouppatient | -0.098347791038885 | 0.420724300002535 | 1.3275728686124 | 0.709612742220925 | 0.829121845527177 | 0.915675240519549 | 0.945245348307718 | NA | abundance | 26 | 9 |
| **Mediterraneibacter** | group | patient | grouppatient | 0.075856197449091 | 0.420724300002535 | 0.721726368057224 | 0.653852502226173 | 0.829121845527177 | 0.531792821213394 | 0.631975142887608 | NA | abundance | 26 | 25 |
| **Odoribacter** | group | patient | grouppatient | 0.808994952491514 | 0.420724300002535 | 0.96960975286881 | 0.700873766495806 | 0.829121845527177 | 0.0287063816892257 | 0.269663801493871 | NA | abundance | 26 | 20 |
| **Oliverpabstia** | group | patient | grouppatient | 0.819063096111971 | 0.420724300002535 | 0.849420493741544 | 0.657792545936558 | 0.829121845527177 | 0.0294147567563192 | 0.269663801493871 | NA | abundance | 26 | 15 |
| **Parabacteroides** | group | patient | grouppatient | 0.199701546019173 | 0.420724300002535 | 0.52387905221046 | 0.707869906103256 | 0.829121845527177 | 0.465033774474719 | 0.608455405854772 | NA | abundance | 26 | 23 |
| **Turicibacter** | group | patient | grouppatient | -0.636980832330281 | 0.420724300002535 | 2.51942091937162 | 0.68342932990719 | 0.829121845527177 | 0.175669562429156 | 0.322309761189383 | NA | abundance | 26 | 12 |
| **Akkermansia** | group | patient | grouppatient | -0.36755440682482 | 0 | 0.82418432662414 | 0.655625119420826 | 0.829121845527177 | 0.404609397504086 | 0.54995452087934 | NA | prevalence | 26 | 16 |
| **Dorea** | group | patient | grouppatient | -0.568737376960043 | 0 | 1.33778991366626 | 0.670740410510825 | 0.829121845527177 | 0.56125981685026 | 0.644068642287184 | NA | prevalence | 26 | 24 |
| **Eggerthella** | group | patient | grouppatient | 0.355178650198049 | 0 | 0.931898824816763 | 0.703103562102573 | 0.829121845527177 | 0.0178444302179786 | 0.245352627347026 | NA | prevalence | 26 | 19 |
| **Flavonifractor** | group | patient | grouppatient | 0.481122680738174 | 0 | 1.15981678445056 | 0.678268954427243 | 0.829121845527177 | 0.103600708099664 | 0.290081982679058 | NA | prevalence | 26 | 22 |
| **GGB3256** | group | patient | grouppatient | -0.322296929847208 | 0 | 0.842889311311601 | 0.702185756923298 | 0.829121845527177 | 0.768272804504625 | 0.814834792656421 | NA | prevalence | 26 | 10 |
| **GGB3345** | group | patient | grouppatient | -0.355178650198048 | 0 | 0.931898824816763 | 0.703103562102573 | 0.829121845527177 | 0.612945202324238 | 0.681050224804709 | NA | prevalence | 26 | 7 |
| **GGB3612** | group | patient | grouppatient | 0.328225188365468 | 0 | 0.812553720433775 | 0.686254815214486 | 0.829121845527177 | 0.00551808973956618 | 0.151338487744683 | NA | prevalence | 26 | 13 |
| **GGB3740** | group | patient | grouppatient | -0.355178650198048 | 0 | 0.931898824816763 | 0.703103562102573 | 0.829121845527177 | 0.0772070112916873 | 0.269663801493871 | NA | prevalence | 26 | 7 |
| **GGB4456** | group | patient | grouppatient | 0.36755440682482 | 0 | 0.82418432662414 | 0.655625119420826 | 0.829121845527177 | 0.694141083101858 | 0.75333140801752 | NA | prevalence | 26 | 10 |
| **GGB4599** | group | patient | grouppatient | -0.355178650198048 | 0 | 0.931898824816763 | 0.703103562102573 | 0.829121845527177 | 0.629630166588989 | 0.694080498602035 | NA | prevalence | 26 | 7 |
| **GGB5980** | group | patient | grouppatient | -0.481122680738175 | 0 | 1.15981678445056 | 0.678268954427243 | 0.829121845527177 | 0.701012156978604 | 0.754936169053881 | NA | prevalence | 26 | 4 |
| **Lachnoclostridium** | group | patient | grouppatient | 0.355178650198048 | 0 | 0.931898824816763 | 0.703103562102573 | 0.829121845527177 | 0.911852505163819 | 0.945245348307718 | NA | prevalence | 26 | 19 |
| **Lacrimispora** | group | patient | grouppatient | -0.568737376960044 | 0 | 1.33778991366626 | 0.670740410510824 | 0.829121845527177 | 0.00044835586185466 | 0.0313849103298262 | NA | prevalence | 26 | 24 |
| **Prevotella** | group | patient | grouppatient | -0.328225188365467 | 0 | 0.812553720433775 | 0.686254815214486 | 0.829121845527177 | 0.0341655570686616 | 0.269663801493871 | NA | prevalence | 26 | 13 |
| **Anaerobutyricum** | group | patient | grouppatient | 0.259162261886886 | 0.420724300002535 | 0.413531316170304 | 0.743212962627634 | 0.848055935324798 | 0.277157984878331 | 0.393487153673676 | NA | abundance | 26 | 24 |
| **Faecalibacterium** | group | patient | grouppatient | 0.293711890623087 | 0.420724300002535 | 0.278660951832513 | 0.744212351407476 | 0.848055935324798 | 0.531792821213394 | 0.631975142887608 | NA | abundance | 26 | 25 |
| **GGB3277** | group | patient | grouppatient | -0.257728368715934 | 0.420724300002535 | 1.87064819996318 | 0.741491211293094 | 0.848055935324798 | 0.933173206161288 | 0.953607655931243 | NA | abundance | 26 | 5 |
| **Lachnoclostridium** | group | patient | grouppatient | 0.959744355394497 | 0.420724300002535 | 1.61566433946198 | 0.743499605970105 | 0.848055935324798 | 0.911852505163819 | 0.945245348307718 | NA | abundance | 26 | 19 |
| **GGB3653** | group | patient | grouppatient | 0.747393501457367 | 0.420724300002535 | 1.16468194529642 | 0.78666664563682 | 0.857457973609279 | 0.156400857945543 | 0.321978487862176 | NA | abundance | 26 | 15 |
| **GGB6612** | group | patient | grouppatient | -1.0564260269738 | 0.420724300002535 | 5.01954190067717 | 0.787461404335053 | 0.857457973609279 | 0.954703272105258 | 0.954703272105258 | NA | abundance | 26 | 5 |
| **Mogibacterium** | group | patient | grouppatient | 1.66682947194205 | 0.420724300002535 | 4.25611573280612 | 0.777773288742496 | 0.857457973609279 | 0.181874793814009 | 0.322309761189383 | NA | abundance | 26 | 9 |
| **Phocaeicola** | group | patient | grouppatient | 0.632943744501574 | 0.420724300002535 | 0.689906845335248 | 0.774157640791922 | 0.857457973609279 | 0.27715798487833 | 0.393487153673676 | NA | abundance | 26 | 24 |
| **GGB1420** | group | patient | grouppatient | 0.265902022950879 | 0 | 0.984867192958462 | 0.787169720446687 | 0.857457973609279 | 0.954703272105258 | 0.954703272105258 | NA | prevalence | 26 | 5 |
| **GGB3277** | group | patient | grouppatient | 0.265902022950878 | 0 | 0.984867192958462 | 0.787169720446687 | 0.857457973609279 | 0.933173206161288 | 0.953607655931243 | NA | prevalence | 26 | 5 |
| **GGB51647** | group | patient | grouppatient | 0.265902022950878 | 0 | 0.984867192958462 | 0.787169720446687 | 0.857457973609279 | 0.485935311658459 | 0.621432461557186 | NA | prevalence | 26 | 5 |
| **GGB6612** | group | patient | grouppatient | 0.265902022950879 | 0 | 0.984867192958461 | 0.787169720446687 | 0.857457973609279 | 0.954703272105258 | 0.954703272105258 | NA | prevalence | 26 | 5 |
| **GGB6613** | group | patient | grouppatient | 0.265902022950879 | 0 | 0.984867192958462 | 0.787169720446686 | 0.857457973609279 | 0.954703272105258 | 0.954703272105258 | NA | prevalence | 26 | 5 |
| **Wujia** | group | patient | grouppatient | 0.265902022950878 | 0 | 0.984867192958462 | 0.787169720446687 | 0.857457973609279 | 0.597067377062267 | 0.66871546230974 | NA | prevalence | 26 | 5 |
| **Sutterella** | group | patient | grouppatient | 0.729761791329831 | 0.420724300002535 | 1.12213753001956 | 0.791997100913607 | 0.858580927981565 | 0.0442793242754595 | 0.269663801493871 | NA | abundance | 26 | 12 |

| | | | | | | | | | | | | | | |
|---|---|---|---|---|---|---|---|---|---|---|---|---|---|---|
| **GGB3363** | group | patient | grouppatient | -0.442171087683903 | 0.420724300002535 | 3.28855878938047 | 0.805763392726408 | 0.869656525189294 | 0.54164201691294 | 0.631975142887608 | NA | abundance | 26 | 6 |
| **GGB6613** | group | patient | grouppatient | 0.75290313861016 | 0.420724300002535 | 1.38825430507103 | 0.827577808164476 | 0.889283171053932 | 0.954703272105258 | 0.954703272105258 | NA | abundance | 26 | 5 |
| **Enterocloster** | group | patient | grouppatient | -0.173516301503482 | 0 | 0.863709383217398 | 0.840779405477203 | 0.899523818086964 | 0.200440051030393 | 0.342214721271403 | NA | prevalence | 26 | 18 |
| **Fusicatenibacter** | group | patient | grouppatient | 0.519892059700215 | 0.420724300002535 | 0.487782868097634 | 0.857790271181336 | 0.910517363842832 | 0.531792821213394 | 0.631975142887608 | NA | abundance | 26 | 25 |
| **GGB4266** | group | patient | grouppatient | 0.618559553063091 | 0.420724300002535 | 0.999516141898508 | 0.85848780019467 | 0.910517363842832 | 0.719910117305141 | 0.769369591013128 | NA | abundance | 26 | 5 |
| **GGB1420** | group | patient | grouppatient | 0.667758531273613 | 0.420724300002535 | 1.41645389641163 | 0.873616556241682 | 0.922569208100052 | 0.954703272105258 | 0.954703272105258 | NA | abundance | 26 | 5 |
| **Hydrogenoanaerobacterium** | group | patient | grouppatient | 0.326142183828554 | 0.420724300002535 | 0.598024217099753 | 0.885319977604872 | 0.930915856279801 | 0.224090367675199 | 0.360605189362389 | NA | abundance | 26 | 17 |
| **GGB9730** | group | patient | grouppatient | 0.557879991758053 | 0.420724300002535 | 0.980780229530186 | 0.89413477787253 | 0.936166754610128 | 0.0442793242754595 | 0.269663801493871 | NA | abundance | 26 | 12 |
| **Eubacterium** | group | patient | grouppatient | 0.34038737904212 | 0.420724300002535 | 0.63425992170021 | 0.906633495652586 | 0.94521364440376 | 0.078235349465694 | 0.269663801493871 | NA | abundance | 26 | 21 |
| **GGB9640** | group | patient | grouppatient | -0.083127216343171 | 0 | 0.850579276318447 | 0.922146596241812 | 0.95731320372561 | 0.915675240519549 | 0.945245348307718 | NA | prevalence | 26 | 9 |
| **Gemmiger** | group | patient | grouppatient | 0.357268180699221 | 0.420724300002535 | 0.817410839283264 | 0.941065245221875 | 0.969689742713478 | 0.0576739954980453 | 0.269663801493871 | NA | abundance | 26 | 19 |
| **GGB36331** | group | patient | grouppatient | -0.069261003946302 | 0 | 0.951700375118188 | 0.941984321493093 | 0.969689742713478 | 0.90123783108117 | 0.945245348307718 | NA | prevalence | 26 | 6 |
| **Eubacteriales_unclassified** | group | patient | grouppatient | 0.374723938624595 | 0.420724300002535 | 0.628768728653603 | 0.946019073248526 | 0.969768506049745 | 0.277157984878331 | 0.393487153673676 | NA | abundance | 26 | 24 |
| **Dysosmobacter** | group | patient | grouppatient | 0.465408564259477 | 0.420724300002535 | 1.01387664496678 | 0.966129978242582 | 0.97810679615468 | 0.0103963639731613 | 0.198700175522074 | NA | abundance | 26 | 19 |
| **GGB3267** | group | patient | grouppatient | -0.056151171450771 | 0 | 1.21234981226173 | 0.9630584016004 | 0.97810679615468 | 0.918238338356069 | 0.945245348307718 | NA | prevalence | 26 | 3 |
| **GGB9480** | group | patient | grouppatient | -0.056151171450771 | 0 | 1.21234981226173 | 0.9630584016004 | 0.97810679615468 | 0.242515400596504 | 0.380598595198154 | NA | prevalence | 26 | 3 |
| **Anaerostipes** | group | patient | grouppatient | 0.441898216223467 | 0.420724300002535 | 0.795404238260437 | 0.97981277076953 | 0.987877073409608 | 0.531792821213394 | 0.631975142887608 | NA | abundance | 26 | 25 |
| **GGB9699** | group | patient | grouppatient | 0.406595887613334 | 0.420724300002535 | 1.69818391270453 | 0.993492252861645 | 0.997563942422553 | 0.00648593518905785 | 0.151338487744683 | NA | abundance | 26 | 17 |
| **GGB3175** | group | patient | grouppatient | 0.420724300002535 | 0.420724300002535 | 1.90543126779882 | 1 | 1 | 0.578930528004906 | 0.653631241295862 | NA | abundance | 26 | 9 |
| **GGB9760** | group | patient | grouppatient | -3.06614490160455 | 0 | 1.13178849160409 | 0.00674630015197699 | 0.172417391959094 | 0.00019690833284358 | 0.0275671665981014 | Prevale | prevalence | 26 | 14 |
| **Ellagibacter** | group | patient | grouppatient | NA | NA | NA | NA | NA | 0.11876207683137 | 0.302769653414133 | contra | abundance | 26 | 8 |
| **GGB13489** | group | patient | grouppatient | NA | NA | NA | NA | NA | 0.142558905011677 | 0.321907204865078 | contra | abundance | 26 | 7 |
| **GGB1495** | group | patient | grouppatient | NA | NA | NA | NA | NA | 0.252826209667345 | 0.380598595198154 | contra | abundance | 26 | 4 |
| **GGB3005** | group | patient | grouppatient | NA | NA | NA | NA | NA | 0.0985889051946244 | 0.281682586270357 | contra | abundance | 26 | 9 |
| **GGB3033** | group | patient | grouppatient | NA | NA | NA | NA | NA | 0.0985889051946245 | 0.281682586270357 | contra | abundance | 26 | 9 |
| **GGB3304** | group | patient | grouppatient | NA | NA | NA | NA | NA | 0.118762076831369 | 0.302769653414133 | contra | abundance | 26 | 8 |
| **GGB33469** | group | patient | grouppatient | NA | NA | NA | NA | NA | 0.0811907139073936 | 0.269663801493871 | contra | abundance | 26 | 10 |
| **GGB33512** | group | patient | grouppatient | NA | NA | NA | NA | NA | 0.0811907139073935 | 0.269663801493871 | contra | abundance | 26 | 10 |
| **GGB4567** | group | patient | grouppatient | NA | NA | NA | NA | NA | 0.171254616937499 | 0.322309761189383 | contra | abundance | 26 | 6 |
| **GGB6544** | group | patient | grouppatient | NA | NA | NA | NA | NA | 0.252826209667345 | 0.380598595198154 | contra | abundance | 26 | 4 |
| **GGB6606** | group | patient | grouppatient | NA | NA | NA | NA | NA | 0.1712546169375 | 0.322309761189383 | contra | abundance | 26 | 6 |
| **GGB87445** | group | patient | grouppatient | NA | NA | NA | NA | NA | 0.142558905011677 | 0.321907204865078 | contra | abundance | 26 | 7 |
| **GGB9176** | group | patient | grouppatient | NA | NA | NA | NA | NA | 0.171254616937499 | 0.322309761189383 | contra | abundance | 26 | 6 |
| **GGB9261** | group | patient | grouppatient | NA | NA | NA | NA | NA | 0.206867593166369 | 0.344779321943948 | contra | abundance | 26 | 5 |
| **GGB9342** | group | patient | grouppatient | NA | NA | NA | NA | NA | 0.142558905011677 | 0.321907204865078 | contra | abundance | 26 | 7 |
| **GGB9345** | group | patient | grouppatient | NA | NA | NA | NA | NA | 0.0985889051946245 | 0.281682586270357 | contra | abundance | 26 | 9 |
| **GGB9453** | group | patient | grouppatient | NA | NA | NA | NA | NA | 0.0985889051946243 | 0.281682586270357 | contra | abundance | 26 | 9 |
| **GGB9509** | group | patient | grouppatient | NA | NA | NA | NA | NA | 0.0659810329776004 | 0.269663801493871 | contra | abundance | 26 | 11 |

| | | | | | | | | | | | | | | |
|---|---|---|---|---|---|---|---|---|---|---|---|---|---|---|
| **GGB9522** | group | patient | grouppatient | NA | NA | NA | NA | NA | 0.0659810329776006 | 0.269663801493871 | contra | abundance | 26 | 11 |
| **GGB9524** | group | patient | grouppatient | NA | NA | NA | NA | NA | 0.206867593166369 | 0.344779321943948 | contra | abundance | 26 | 5 |
| **GGB9602** | group | patient | grouppatient | NA | NA | NA | NA | NA | 0.0985889051946248 | 0.281682586270357 | contra | abundance | 26 | 9 |
| **GGB9608** | group | patient | grouppatient | NA | NA | NA | NA | NA | 0.0659810329776004 | 0.269663801493871 | contra | abundance | 26 | 11 |
| **GGB9634** | group | patient | grouppatient | NA | NA | NA | NA | NA | 0.142558905011677 | 0.321907204865078 | contra | abundance | 26 | 7 |
| **GGB9708** | group | patient | grouppatient | NA | NA | NA | NA | NA | 0.0659810329776003 | 0.269663801493871 | contra | abundance | 26 | 11 |
| **GGB9712** | group | patient | grouppatient | NA | NA | NA | NA | NA | 0.252826209667345 | 0.380598595198154 | contra | abundance | 26 | 4 |
| **GGB9737** | group | patient | grouppatient | NA | NA | NA | NA | NA | 0.252826209667345 | 0.380598595198154 | contra | abundance | 26 | 4 |
| **GGB9747** | group | patient | grouppatient | NA | NA | NA | NA | NA | 0.142558905011677 | 0.321907204865078 | contra | abundance | 26 | 7 |
| **Methanobrevibacter** | group | patient | grouppatient | NA | NA | NA | NA | NA | 0.11876207683137 | 0.302769653414133 | contra | abundance | 26 | 8 |
| **Monoglobus** | group | patient | grouppatient | NA | NA | NA | NA | NA | 0.11876207683137 | 0.302769653414133 | contra | abundance | 26 | 8 |
| **Parolsenella** | group | patient | grouppatient | NA | NA | NA | NA | NA | 0.252826209667345 | 0.380598595198154 | contra | abundance | 26 | 4 |
| **Senegalimassilia** | group | patient | grouppatient | NA | NA | NA | NA | NA | 0.0985889051946242 | 0.281682586270357 | contra | abundance | 26 | 9 |
| **Bacteroides** | group | patient | grouppatient | NA | 0 | NA | NA | NA | 0.432389037700174 | 0.571079861113437 | All logi | prevalence | 26 | 26 |
| **Bifidobacterium** | group | patient | grouppatient | NA | 0 | NA | NA | NA | 0.54169297961795 | 0.631975142887608 | All logi | prevalence | 26 | 26 |
| **Clostridium** | group | patient | grouppatient | NA | 0 | NA | NA | NA | 0.286946691101012 | 0.401725367541417 | All logi | prevalence | 26 | 26 |
| **Streptococcus** | group | patient | grouppatient | NA | 0 | NA | NA | NA | 0.026359833512628 | 0.269663801493871 | All logi | prevalence | 26 | 26 |

### Supplementary Table 2: Differential Pathway Abundance Analysis (MaAsLin3), Healthy controls vs. Patients at baseline

| feature | metadata | value | name | coef | null_hypothesis | stderr | pval_individual | qval_individual | pval_joint | qval_joint | error | model | N | N_not_zero |
|---|---|---|---|---|---|---|---|---|---|---|---|---|---|---|
| ARGSYN-PWY: L-arginine biosynthesis I (via L-ornithine) | group | patient | grouppatient | -0.339832163977929 | 0.0482358175877709 | 0.070783791157612 | 7.09537826051321e-05 | 0.00799791848342011 | 7.09537826051321e-05 | 0.00734183923282705 | NA | abundance | 26 | 26 |
| ARGSYNBSUB-PWY: L-arginine biosynthesis II (acetyl cycle) | group | patient | grouppatient | -0.353868605128583 | 0.0482358175877709 | 0.0807173623383707 | 0.000156209345379299 | 0.00799791848342011 | 0.000156209345379299 | 0.00734183923282705 | NA | abundance | 26 | 26 |
| HISTSYN-PWY: L-histidine biosynthesis | group | patient | grouppatient | -0.383209900482643 | 0.0482358175877709 | 0.0878160282784228 | 0.00015558987309916 | 0.00799791848342011 | 0.00015558987309916 | 0.00734183923282705 | NA | abundance | 26 | 26 |
| NONMEVIPP-PWY: methylerythritol phosphate pathway I | group | patient | grouppatient | -0.481384203469132 | 0.0482358175877709 | 0.105538272084725 | 8.61377185228385e-05 | 0.00799791848342011 | 8.61377185228385e-05 | 0.00734183923282705 | NA | abundance | 26 | 26 |
| PWY-7238: sucrose biosynthesis II | group | patient | grouppatient | -0.328709661660223 | 0.0482358175877709 | 0.0731113727549431 | 0.000130359272577207 | 0.00799791848342011 | 0.000130359272577207 | 0.00734183923282705 | NA | abundance | 26 | 26 |
| GLUTORN-PWY: L-ornithine biosynthesis I | group | patient | grouppatient | -0.401613984756696 | 0.0482358175877709 | 0.104964648566585 | 0.000493851056067918 | 0.0210709783922312 | 0.000493851056067918 | 0.0193424996959934 | NA | abundance | 26 | 26 |
| PWY-6163: chorismate biosynthesis from 3-dehydroquinate | group | patient | grouppatient | -0.405369430081666 | 0.0482358175877709 | 0.107666566653356 | 0.000601791008974395 | 0.022008356899635 | 0.000601791008974395 | 0.0202029838727118 | NA | abundance | 26 | 26 |
| ARO-PWY: chorismate biosynthesis I | group | patient | grouppatient | -0.340380904876378 | 0.0482358175877709 | 0.0964696184689661 | 0.00104569783115105 | 0.0274834506389336 | 0.00104569783115105 | 0.0252289488287085 | NA | abundance | 26 | 26 |
| COMPLETE-ARO-PWY: superpathway of aromatic amino acid biosynthesis | group | patient | grouppatient | -0.331818293619228 | 0.0482358175877709 | 0.0970715825489265 | 0.00129281963359129 | 0.0274834506389336 | 0.00129281963359129 | 0.0252289488287085 | NA | abundance | 26 | 26 |
| GLYCOGENSYNTH-PWY: glycogen biosynthesis I (from ADP-D-Glucose) | group | patient | grouppatient | -0.27903169586976 | 0.0482358175877709 | 0.0770452546118564 | 0.000880147684649168 | 0.0274834506389336 | 0.000880147684649168 | 0.0252289488287085 | NA | abundance | 26 | 26 |
| PWY-7953: UDP-N-acetylmuramoyl-pentapeptide biosynthesis III (meso-diaminopimelate containing | group | patient | grouppatient | -0.242422571624568 | 0.0482358175877709 | 0.0693576737162474 | 0.0012745107934875 | 0.0274834506389336 | 0.0012745107934875 | 0.0252289488287085 | NA | abundance | 26 | 26 |
| PWY0-1297: superpathway of purine deoxyribonucleosides degradation | group | patient | grouppatient | 1.14180158384212 | 0.0482358175877709 | 0.302692318533899 | 0.00150300120681668 | 0.0274834506389336 | 0.00150300120681668 | 0.0252289488287085 | NA | abundance | 26 | 26 |
| TRNA-CHARGING-PWY: tRNA charging | group | patient | grouppatient | -0.239505575166486 | 0.0482358175877709 | 0.0698823351576745 | 0.0014445790484896 | 0.0274834506389336 | 0.0014445790484896 | 0.0252289488287085 | NA | abundance | 26 | 26 |
| VALSYN-PWY: L-valine biosynthesis | group | patient | grouppatient | -0.206782269083099 | 0.0482358175877709 | 0.0570015992835323 | 0.00118761348409757 | 0.0274834506389336 | 0.00118761348409757 | 0.0252289488287085 | NA | abundance | 26 | 26 |
| FERMENTATION-PWY: mixed acid fermentation | group | patient | grouppatient | -0.890822125954581 | 0.0482358175877709 | 0.269252074902098 | 0.00208031964149091 | 0.0355041218814449 | 0.00208031964149091 | 0.0325916743833576 | NA | abundance | 26 | 26 |
| PEPTIDOGLYCANSYN-PWY: peptidoglycan biosynthesis I (meso-diaminopimelate containing) | group | patient | grouppatient | -0.208268278599191 | 0.0482358175877709 | 0.0658152885237999 | 0.00254872201340461 | 0.0385207654396993 | 0.00254872201340461 | 0.035360858899724 | NA | abundance | 26 | 26 |
| PWY-6151: S-adenosyl-L-methionine salvage I | group | patient | grouppatient | -0.212907904434171 | 0.0482358175877709 | 0.06750711259193 | 0.00270849131997886 | 0.0385207654396993 | 0.00270849131997886 | 0.035360858899724 | NA | abundance | 26 | 26 |
| PWY-6387: UDP-N-acetylmuramoyl-pentapeptide biosynthesis I (meso-diaminopimelate containing) | group | patient | grouppatient | -0.198154677275127 | 0.0482358175877709 | 0.0620108679357628 | 0.00258453722588281 | 0.0385207654396993 | 0.00258453722588281 | 0.035360858899724 | NA | abundance | 26 | 26 |
| COA-PWY-1: superpathway of coenzyme A biosynthesis III (mammals) | group | patient | grouppatient | -0.16424392611843 | 0.0482358175877709 | 0.0523028822417023 | 0.00319503579138125 | 0.0395134280268156 | 0.00319503579138125 | 0.0362720921339909 | NA | abundance | 26 | 26 |
| PWY-6386: UDP-N-acetylmuramoyl-pentapeptide biosynthesis II (lysine-containing) | group | patient | grouppatient | -0.185701437493486 | 0.0482358175877709 | 0.0602389541115944 | 0.00324133589282471 | 0.0395134280268156 | 0.00324133589282471 | 0.0362720921339909 | NA | abundance | 26 | 26 |
| PWY-6969: TCA cycle V (2-oxoglutarate synthase) | group | patient | grouppatient | -0.748929413882537 | 0.0482358175877709 | 0.240125987718082 | 0.00319226844644505 | 0.0395134280268156 | 0.00319226844644505 | 0.0362720921339909 | NA | abundance | 26 | 26 |
| PWY-5686: UMP biosynthesis I | group | patient | grouppatient | -0.170873405230639 | 0.0482358175877709 | 0.0561069596598754 | 0.00377358921222959 | 0.0395253057603577 | 0.00377358921222959 | 0.0362829955222034 | NA | abundance | 26 | 26 |
| PWY-6700: queuosine biosynthesis I (de novo) | group | patient | grouppatient | -0.648424265258134 | 0.0482358175877709 | 0.211182255654775 | 0.00343399775092545 | 0.0395253057603577 | 0.00343399775092545 | 0.0362829955222034 | NA | abundance | 26 | 26 |
| PWY-7790: UMP biosynthesis II | group | patient | grouppatient | -0.170873405230639 | 0.0482358175877709 | 0.0561069596598754 | 0.00385989314065993 | 0.0395253057603577 | 0.00385989314065993 | 0.0362829955222034 | NA | abundance | 26 | 26 |
| PWY-7791: UMP biosynthesis III | group | patient | grouppatient | -0.170873405230639 | 0.0482358175877709 | 0.0561069596598754 | 0.00359264854958341 | 0.0395253057603577 | 0.00359264854958341 | 0.0362829955222034 | NA | abundance | 26 | 26 |
| COA-PWY: coenzyme A biosynthesis I (prokaryotic) | group | patient | grouppatient | -0.162966457222797 | 0.0482358175877709 | 0.0540702929635543 | 0.00407965550692002 | 0.0401689157604433 | 0.00407965550692002 | 0.0368738093894694 | NA | abundance | 26 | 26 |
| PWY-7851: coenzyme A biosynthesis II (eukaryotic) | group | patient | grouppatient | -0.159443568224428 | 0.0482358175877709 | 0.0548162326189487 | 0.00493931980602857 | 0.0468320692719746 | 0.00493931980602857 | 0.042990376089508 | NA | abundance | 26 | 26 |
| PWY-5188: tetrapyrrole biosynthesis I (from glutamate) | group | patient | grouppatient | -0.49270440415082 | 0.0482358175877709 | 0.173913147233986 | 0.00573066506640418 | 0.0523946520356954 | 0.00573066506640418 | 0.0480966532358923 | NA | abundance | 26 | 26 |
| PWY-6629: superpathway of L-tryptophan biosynthesis | group | patient | grouppatient | -0.278638166882894 | 0.0482358175877709 | 0.102503024542552 | 0.00660847736350645 | 0.0583369036226776 | 0.00660847736350645 | 0.0535514544973798 | NA | abundance | 26 | 26 |
| PWY-6385: peptidoglycan biosynthesis III (mycobacteria) | group | patient | grouppatient | -0.167974498714341 | 0.0482358175877709 | 0.0628937188338618 | 0.00736961686233495 | 0.0628873972252582 | 0.00736961686233495 | 0.0577286654216238 | NA | abundance | 26 | 26 |
| BRANCHED-CHAIN-AA-SYN-PWY: superpathway of branched chain amino acid biosynthesis | group | patient | grouppatient | -0.167305610796582 | 0.0482358175877709 | 0.0666787927825685 | 0.0100152600160031 | 0.0801220801280245 | 0.0100152600160031 | 0.0735495657425225 | NA | abundance | 26 | 26 |
| PWY66-409: superpathway of purine nucleotide salvage | group | patient | grouppatient | 0.812007174677121 | 0.0482358175877709 | 0.269498869826028 | 0.00988779992311084 | 0.0801220801280245 | 0.00988779992311084 | 0.0735495657425225 | NA | abundance | 26 | 26 |
| PYRIDNUCSYN-PWY: NAD de novo biosynthesis I (from aspartate) | group | patient | grouppatient | -0.216495495071546 | 0.0482358175877709 | 0.0895775228469991 | 0.0118678620949859 | 0.092065839282315 | 0.0118678620949859 | 0.0845135634036876 | NA | abundance | 26 | 26 |
| PWY-6549: L-glutamine biosynthesis III | group | patient | grouppatient | -0.637359446592335 | 0.0482358175877709 | 0.251119341494277 | 0.0125241043702746 | 0.0942991387879501 | 0.0125241043702746 | 0.0865636625592511 | NA | abundance | 26 | 26 |
| PWY-724: superpathway of L-lysine, L-threonine and L-methionine biosynthesis II | group | patient | grouppatient | -0.0864101811826729 | 0.0482358175877709 | 0.0325980178747144 | 0.0138078243384523 | 0.100994372304108 | 0.0138078243384523 | 0.0927096777010368 | NA | abundance | 26 | 26 |
| CALVIN-PWY: Calvin-Benson-Bassham cycle | group | patient | grouppatient | -0.296365543977019 | 0.0482358175877709 | 0.12745332566122 | 0.0160951816426669 | 0.104447024271941 | 0.0160951816426669 | 0.0958791043121336 | NA | abundance | 26 | 26 |
| ILEUSYN-PWY: L-isoleucine biosynthesis I (from threonine) | group | patient | grouppatient | -0.149484098526364 | 0.0482358175877709 | 0.0652095888564301 | 0.0153889532577307 | 0.104447024271941 | 0.0153889532577307 | 0.0958791043121336 | NA | abundance | 26 | 26 |
| PWY-1042: glycolysis IV | group | patient | grouppatient | -0.187854098197027 | 0.0482358175877709 | 0.0818296686830031 | 0.0153459218005365 | 0.104447024271941 | 0.0153459218005365 | 0.0958791043121336 | NA | abundance | 26 | 26 |
| PWY-6609: adenine and adenosine salvage III | group | patient | grouppatient | -0.357945732366715 | 0.0482358175877709 | 0.15250785915987 | 0.0163198475424908 | 0.104447024271941 | 0.0163198475424908 | 0.0958791043121336 | NA | abundance | 26 | 26 |
| PWY-6612: superpathway of tetrahydrofolate biosynthesis | group | patient | grouppatient | 1.03536462973129 | 0.0482358175877709 | 0.379298656513569 | 0.0161000364453403 | 0.104447024271941 | 0.0161000364453403 | 0.0958791043121336 | NA | abundance | 26 | 26 |
| FOLSYN-PWY: superpathway of tetrahydrofolate biosynthesis and salvage | group | patient | grouppatient | 0.98364366646844 | 0.0482358175877709 | 0.366733210452459 | 0.01806960191275 | 0.112824831455219 | 0.01806960191275 | 0.103569669499908 | NA | abundance | 26 | 26 |
| GLUCOSE1PMETAB-PWY: glucose and glucose-1-phosphate degradation | group | patient | grouppatient | 0.509874801396151 | 0.0482358175877709 | 0.178811374014202 | 0.0207883346083562 | 0.122359870616842 | 0.0411445143609233 | 0.163880692793508 | NA | abundance | 26 | 21 |
| POLYISOPRENSYN-PWY: polyisoprenoid biosynthesis (E. coli) | group | patient | grouppatient | 0.903636783057334 | 0.0482358175877709 | 0.341483750845541 | 0.0202245267909783 | 0.122359870616842 | 0.0202245267909783 | 0.113161042759045 | NA | abundance | 26 | 26 |
| PWY-3841: folate transformations II (plants) | group | patient | grouppatient | -0.141785240227419 | 0.0482358175877709 | 0.0663245317810116 | 0.0210306027622696 | 0.122359870616842 | 0.0210306027622696 | 0.114934689514729 | NA | abundance | 26 | 26 |
| PWY4FS-7: phosphatidylglycerol biosynthesis I (plastidic) | group | patient | grouppatient | 0.578062052662308 | 0.0482358175877709 | 0.213000610382358 | 0.0220419847255684 | 0.123498076044799 | 0.0220419847255684 | 0.11588664948301 | NA | abundance | 26 | 26 |
| PWY4FS-8: phosphatidylglycerol biosynthesis II (non-plastidic) | group | patient | grouppatient | 0.578047447408897 | 0.0482358175877709 | 0.212999392042262 | 0.0221910605392998 | 0.123498076044799 | 0.0221910605392998 | 0.11588664948301 | NA | abundance | 26 | 26 |

| | | | | | | | | | | | | | | |
|---|---|---|---|---|---|---|---|---|---|---|---|---|---|---|
| **HSERMETANA-PWY: L-methionine biosynthesis III** | group | patient | grouppatient | -0.16179167939545 | 0.0482358175877709 | 0.0775600699593 | 0.023418327647841 | 0.126306070064167 | 0.023418327647841 | 0.118411940685157 | NA | abundance | 26 | 26 |
| **LACTOSECAT-PWY: lactose and galactose degradation I** | group | patient | grouppatient | -0.795994942689485 | 0.0482358175877709 | 0.346975679698398 | 0.0236823881370314 | 0.126306070064167 | 0.0236823881370314 | 0.118411940685157 | NA | abundance | 26 | 26 |
| **PWY-5981: CDP-diacylglycerol biosynthesis III** | group | patient | grouppatient | 2.21011091329745 | 0.0482358175877709 | 0.897525124139424 | 0.0245328101206838 | 0.128171416140715 | 0.04846376146895 | 0.177952874143801 | NA | abundance | 26 | 25 |
| **PWY-7221: guanosine ribonucleotides de novo biosynthesis** | group | patient | grouppatient | -0.126587902031459 | 0.0482358175877709 | 0.0628283485298073 | 0.0268865389508887 | 0.13765907942855 | 0.0268865389508887 | 0.131632013613726 | NA | abundance | 26 | 26 |
| **1CMET2-PWY: folate transformations III (E. coli)** | group | patient | grouppatient | -0.316999858216466 | 0.0482358175877709 | 0.151295029620028 | 0.0276953127956533 | 0.139019609327201 | 0.0276953127956533 | 0.132824459326092 | NA | abundance | 26 | 26 |
| **CENTFERM-PWY: pyruvate fermentation to butanoate** | group | patient | grouppatient | -0.771622058003535 | 0.0482358175877709 | 0.359216746843147 | 0.0323664875534683 | 0.144808461872142 | 0.0323664875534683 | 0.1376771299747 | NA | abundance | 26 | 26 |
| **OANTIGEN-PWY: O-antigen building blocks biosynthesis (E. coli)** | group | patient | grouppatient | -0.22314428896367 | 0.0482358175877709 | 0.112955692616725 | 0.0317308748182228 | 0.144808461872142 | 0.0317308748182228 | 0.1376771299747 | NA | abundance | 26 | 26 |
| **P42-PWY: incomplete reductive TCA cycle** | group | patient | grouppatient | -1.37135221303118 | 0.0482358175877709 | 0.6248148007558 | 0.0325801166888491 | 0.144808461872142 | 0.0325801166888491 | 0.1376771299747 | NA | abundance | 26 | 26 |
| **PWY-2942: L-lysine biosynthesis III** | group | patient | grouppatient | -0.239061071782445 | 0.0482358175877709 | 0.11991028951951 | 0.0314157601397214 | 0.144808461872142 | 0.0314157601397214 | 0.1376771299747 | NA | abundance | 26 | 26 |
| **PWY-6121: 5-aminoimidazole ribonucleotide biosynthesis I** | group | patient | grouppatient | -0.138971816968447 | 0.0482358175877709 | 0.072861174423226 | 0.0328081671429072 | 0.144808461872142 | 0.0328081671429072 | 0.1376771299747 | NA | abundance | 26 | 26 |
| **PWY-6590: superpathway of Clostridium acetobutylicum acidogenic fermentation** | group | patient | grouppatient | -0.750210873767313 | 0.0482358175877709 | 0.349377528145562 | 0.0321463711385879 | 0.144808461872142 | 0.0321463711385879 | 0.1376771299747 | NA | abundance | 26 | 26 |
| **TRPSYN-PWY: L-tryptophan biosynthesis** | group | patient | grouppatient | -0.232349161971718 | 0.0482358175877709 | 0.11587603479452 | 0.0306444478216408 | 0.144808461872142 | 0.0306444478216408 | 0.1376771299747 | NA | abundance | 26 | 26 |
| **PANTOSYN-PWY: superpathway of coenzyme A biosynthesis I (bacteria)** | group | patient | grouppatient | -0.336983930240823 | 0.0482358175877709 | 0.172723066075896 | 0.0394179269766175 | 0.170359625342928 | 0.0394179269766175 | 0.161777392061913 | NA | abundance | 26 | 26 |
| **THISYNARA-PWY: superpathway of thiamine diphosphate biosynthesis III (eukaryotes)** | group | patient | grouppatient | 0.27978231614664 | 0.0482358175877709 | 0.0993789491090878 | 0.0399280371897488 | 0.170359625342928 | 0.0399280371897488 | 0.161777392061913 | NA | abundance | 26 | 26 |
| **PWY-4041: γ-glutamyl cycle** | group | patient | grouppatient | 0.531134408403761 | 0.0482358175877709 | 0.226720975666015 | 0.0464611862370106 | 0.189296091351386 | 0.0464611862370106 | 0.176570604363487 | NA | abundance | 26 | 26 |
| **PWY-6122: 5-aminoimidazole ribonucleotide biosynthesis II** | group | patient | grouppatient | -0.148016518471972 | 0.0482358175877709 | 0.0853525987634363 | 0.046584584981005 | 0.189296091351386 | 0.046584584981005 | 0.176570604363487 | NA | abundance | 26 | 26 |
| **PWY-6277: superpathway of 5-aminoimidazole ribonucleotide biosynthesis** | group | patient | grouppatient | -0.148016518471972 | 0.0482358175877709 | 0.0853525987634363 | 0.0463551170903638 | 0.189296091351386 | 0.0463551170903638 | 0.176570604363487 | NA | abundance | 26 | 26 |
| **PWY-5103: L-isoleucine biosynthesis III** | group | patient | grouppatient | -0.127336478959832 | 0.0482358175877709 | 0.075480629091893 | 0.0483650775962827 | 0.193460310385131 | 0.0483650775962827 | 0.177952874143801 | NA | abundance | 26 | 26 |
| **PWY-5097: L-lysine biosynthesis VI** | group | patient | grouppatient | -0.101056995787941 | 0.0482358175877709 | 0.0619060510667243 | 0.0506580532933352 | 0.196491843077179 | 0.0506580532933352 | 0.180373371574754 | NA | abundance | 26 | 26 |
| **SALVADEHYPOX-PWY: adenosine nucleotides degradation II** | group | patient | grouppatient | 0.555126486040545 | 0.0482358175877709 | 0.243085415071302 | 0.0506191190283587 | 0.196491843077179 | 0.0506191190283587 | 0.180373371574754 | NA | abundance | 26 | 26 |
| **PWY-6630: superpathway of L-tyrosine biosynthesis** | group | patient | grouppatient | 0.498941533211211 | 0.0482358175877709 | 0.218708793371917 | 0.0536375977019599 | 0.20494365689107 | 0.0536375977019599 | 0.188131872536725 | NA | abundance | 26 | 26 |
| **PWY-2941: L-lysine biosynthesis II** | group | patient | grouppatient | 0.600999436836605 | 0.0482358175877709 | 0.271599775996761 | 0.0551581578796274 | 0.207654241429186 | 0.0551581578796274 | 0.190620104436948 | NA | abundance | 26 | 26 |
| **P461-PWY: hexitol fermentation to lactate, formate, ethanol and acetate** | group | patient | grouppatient | 0.730527861436365 | 0.0482358175877709 | 0.339987692858764 | 0.0575031097192609 | 0.209215747663459 | 0.0575031097192609 | 0.19205351836294 | NA | abundance | 26 | 26 |
| **PWY-5121: superpathway of geranylgeranyl diphosphate biosynthesis II (via MEP)** | group | patient | grouppatient | 0.433131876829546 | 0.0482358175877709 | 0.189084622834514 | 0.0574216449570701 | 0.209215747663459 | 0.0574216449570701 | 0.19205351836294 | NA | abundance | 26 | 26 |
| **PWY-7199: pyrimidine deoxyribonucleosides salvage** | group | patient | grouppatient | -0.264210413013887 | 0.0482358175877709 | 0.152816632978229 | 0.0580246800160373 | 0.209215747663459 | 0.0580246800160373 | 0.19205351836294 | NA | abundance | 26 | 26 |
| **POLYAMSYN-PWY: superpathway of polyamine biosynthesis I** | group | patient | grouppatient | 0.710974816181916 | 0.0482358175877709 | 0.334814161475394 | 0.0607465629005262 | 0.210150271655874 | 0.0607465629005262 | 0.192911382184103 | NA | abundance | 26 | 26 |
| **PWY-5667: CDP-diacylglycerol biosynthesis I** | group | patient | grouppatient | -0.309019406712154 | 0.0482358175877709 | 0.177191276358105 | 0.0594954905707814 | 0.210150271655874 | 0.0594954905707814 | 0.192911382184103 | NA | abundance | 26 | 26 |
| **PWY0-1319: CDP-diacylglycerol biosynthesis II** | group | patient | grouppatient | -0.308968529996388 | 0.0482358175877709 | 0.177195469944907 | 0.0600841597774608 | 0.210150271655874 | 0.0600841597774608 | 0.192911382184103 | NA | abundance | 26 | 26 |
| **PWY-5695: inosine 5'-phosphate degradation** | group | patient | grouppatient | -0.170898743446551 | 0.0482358175877709 | 0.105720882080791 | 0.0626874119698198 | 0.213973032856985 | 0.0626874119698198 | 0.196420557505435 | NA | abundance | 26 | 26 |
| **ARG+POLYAMINE-SYN: superpathway of arginine and polyamine biosynthesis** | group | patient | grouppatient | 0.639900858492605 | 0.0482358175877709 | 0.306921242426054 | 0.0679150519494542 | 0.224054416318275 | 0.0679150519494542 | 0.205872735384621 | NA | abundance | 26 | 26 |
| **ASPASN-PWY: superpathway of L-aspartate and L-asparagine biosynthesis** | group | patient | grouppatient | 0.291773231592312 | 0.0482358175877709 | 0.121138866089437 | 0.0683322270638316 | 0.224054416318275 | 0.0683322270638316 | 0.205872735384621 | NA | abundance | 26 | 26 |
| **PWY-5840: superpathway of menaquinol-7 biosynthesis** | group | patient | grouppatient | 1.22863151883094 | 0.0482358175877709 | 0.614251037195214 | 0.0694130824665904 | 0.224054416318275 | 0.134007988915667 | 0.324658529847235 | NA | abundance | 26 | 22 |
| **PWY-6527: stachyose degradation** | group | patient | grouppatient | 0.248885489805925 | 0.0482358175877709 | 0.0971729825713498 | 0.0674664704941699 | 0.224054416318275 | 0.0674664704941699 | 0.205872735384621 | NA | abundance | 26 | 26 |
| **PYRIDNUCSAL-PWY: NAD salvage pathway I (PNC VI cycle)** | group | patient | grouppatient | 1.39839060973821 | 0.0482358175877709 | 0.711354047495525 | 0.0700170050994608 | 0.224054416318275 | 0.0700170050994608 | 0.208278432890801 | NA | abundance | 26 | 26 |
| **PWY0-1296: purine ribonucleosides degradation** | group | patient | grouppatient | -0.273826756047759 | 0.0482358175877709 | 0.167691974399887 | 0.072612716910922 | 0.229492043570321 | 0.072612716910922 | 0.213299855925833 | NA | abundance | 26 | 26 |
| **PWY-5845: superpathway of menaquinol-9 biosynthesis** | group | patient | grouppatient | 0.980114333836812 | 0.0482358175877709 | 0.519561019441475 | 0.0911841592363792 | 0.270823130761446 | 0.174053767577113 | 0.38567274579689 | NA | abundance | 26 | 19 |
| **PWY-622: starch biosynthesis** | group | patient | grouppatient | 1.20288637680295 | 0.0482358175877709 | 0.654765323324633 | 0.0920375483447102 | 0.270823130761446 | 0.175604186384116 | 0.38567274579689 | NA | abundance | 26 | 24 |
| **PWY-6606: guanosine nucleotides degradation II** | group | patient | grouppatient | 0.392003877509668 | 0.0482358175877709 | 0.19184080334899 | 0.0915157682727008 | 0.270823130761446 | 0.0915157682727008 | 0.256444928744021 | NA | abundance | 26 | 26 |
| **PWY-6823: molybdopterin biosynthesis** | group | patient | grouppatient | -0.356844592299352 | 0.0482358175877709 | 0.227423248273073 | 0.0912630689572964 | 0.270823130761446 | 0.0912630689572964 | 0.256444928744021 | NA | abundance | 26 | 26 |
| **PWY0-1298: superpathway of pyrimidine deoxyribonucleosides degradation** | group | patient | grouppatient | 0.750527117380566 | 0.0482358175877709 | 0.397892082494378 | 0.0916654213382883 | 0.270823130761446 | 0.0916654213382883 | 0.256444928744021 | NA | abundance | 26 | 26 |
| **RHAMCAT-PWY: L-rhamnose degradation I** | group | patient | grouppatient | -0.407064085711797 | 0.0482358175877709 | 0.252600678063546 | 0.087037025217356 | 0.270823130761446 | 0.087037025217356 | 0.252514826247885 | NA | abundance | 26 | 26 |
| **DTDPRHAMSYN-PWY: dTDP-β-L-rhamnose biosynthesis** | group | patient | grouppatient | -0.254427777251259 | 0.0482358175877709 | 0.174156344020458 | 0.101366996146688 | 0.293119938495187 | 0.101366996146688 | 0.278461278780288 | NA | abundance | 26 | 26 |
| **RIBOSYN2-PWY: flavin biosynthesis I (bacteria and plants)** | group | patient | grouppatient | -0.131472393654953 | 0.0482358175877709 | 0.0987967833456165 | 0.101904978617467 | 0.293119938495187 | 0.101904978617467 | 0.278461278780288 | NA | abundance | 26 | 26 |
| **FASYN-INITIAL-PWY: superpathway of fatty acid biosynthesis initiation (E. coli)** | group | patient | grouppatient | -1.84445703248172 | 0 | 1.13666845661122 | 0.104656349736653 | 0.297689172584258 | 0.198359747933105 | 0.421561701686495 | NA | prevalence | 26 | 22 |
| **ANAGLYCOLYSIS-PWY: glycolysis III (from glucose)** | group | patient | grouppatient | -0.0649565297968491 | 0.0482358175877709 | 0.0578625155791326 | 0.113632574596847 | 0.314029564526522 | 0.113632574596847 | 0.300940506011959 | NA | abundance | 26 | 26 |
| **METSYN-PWY: superpathway of L-homoserine and L-methionine biosynthesis** | group | patient | grouppatient | 1.15207276497931 | 0.0482358175877709 | 0.675546469563998 | 0.115602597027463 | 0.314029564526522 | 0.115602597027463 | 0.300940506011959 | NA | abundance | 26 | 26 |
| **PANTO-PWY: phosphopantothenate biosynthesis I** | group | patient | grouppatient | -0.291640009210719 | 0.0482358175877709 | 0.205669250409385 | 0.116534408711014 | 0.314029564526522 | 0.116534408711014 | 0.300940506011959 | NA | abundance | 26 | 26 |
| **PWY-5347: superpathway of L-methionine biosynthesis (transsulfuration)** | group | patient | grouppatient | 1.1371098538235 | 0.0482358175877709 | 0.666983524939765 | 0.115874214849785 | 0.314029564526522 | 0.115874214849785 | 0.300940506011959 | NA | abundance | 26 | 26 |

| | | | | | | | | | | | | | | |
|---|---|---|---|---|---|---|---|---|---|---|---|---|---|---|
| PWY-7456: β-(1,4)-mannan degradation | group | patient | grouppatient | -0.536733904951361 | 0.0482358175877709 | 0.357193458592475 | 0.115893671813868 | 0.314029564526522 | 0.115893671813868 | 0.300940506011959 | NA | abundance | 26 | 26 |
| MET-SAM-PWY: superpathway of S-adenosyl-L-methionine biosynthesis | group | patient | grouppatient | 1.12545604285748 | 0.0482358175877709 | 0.667889079924864 | 0.120145566946195 | 0.317085207610576 | 0.120145566946195 | 0.303593636907051 | NA | abundance | 26 | 26 |
| PWY3O-4107: NAD salvage pathway V (PNC V cycle) | group | patient | grouppatient | 1.396099133303 | 0.0482358175877709 | 0.835526169336345 | 0.119856574074604 | 0.317085207610576 | 0.119856574074604 | 0.303593636907051 | NA | abundance | 26 | 26 |
| PWY-7977: L-methionine biosynthesis IV | group | patient | grouppatient | -0.318532245713247 | 0.0482358175877709 | 0.22848269966005 | 0.125398150838717 | 0.327570679741954 | 0.125398150838717 | 0.313495377096792 | NA | abundance | 26 | 26 |
| PWY-5676: acetyl-CoA fermentation to butanoate II | group | patient | grouppatient | 0.454537452763902 | 0.0482358175877709 | 0.258422718709691 | 0.132945362408033 | 0.33118628764971 | 0.132945362408033 | 0.324658529847235 | NA | abundance | 26 | 26 |
| PWY-5897: superpathway of menaquinol-11 biosynthesis | group | patient | grouppatient | 1.0211063339262 | 0.0482358175877709 | 0.618894130643006 | 0.132048506150435 | 0.33118628764971 | 0.246660204324308 | 0.47349023097295 | NA | abundance | 26 | 22 |
| PWY-5898: superpathway of menaquinol-12 biosynthesis | group | patient | grouppatient | 1.0211063339262 | 0.0482358175877709 | 0.618894130643006 | 0.13235974751887 | 0.33118628764971 | 0.247200392274481 | 0.47349023097295 | NA | abundance | 26 | 22 |
| PWY-5899: superpathway of menaquinol-13 biosynthesis | group | patient | grouppatient | 1.0211063339262 | 0.0482358175877709 | 0.618894130643006 | 0.132169929110235 | 0.33118628764971 | 0.246870968059465 | 0.47349023097295 | NA | abundance | 26 | 22 |
| PWY-7282: 4-amino-2-methyl-5-diphosphomethylpyrimidine biosynthesis II | group | patient | grouppatient | 0.471863259391568 | 0.0482358175877709 | 0.269966690819914 | 0.133250732921563 | 0.33118628764971 | 0.133250732921563 | 0.324658529847235 | NA | abundance | 26 | 26 |
| PENTOSE-P-PWY: pentose phosphate pathway | group | patient | grouppatient | 0.385002940075403 | 0.0482358175877709 | 0.217347910886661 | 0.139171126408718 | 0.342575080390691 | 0.139171126408718 | 0.333726680673967 | NA | abundance | 26 | 26 |
| PWY-5941: glycogen degradation II | group | patient | grouppatient | -0.0693092044153886 | 0.0482358175877709 | 0.0686001273105279 | 0.146128812781883 | 0.349616598805252 | 0.146128812781883 | 0.339856215863755 | NA | abundance | 26 | 26 |
| PWY-7383: anaerobic energy metabolism (invertebrates, cytosol) | group | patient | grouppatient | -0.73301067328915 | 0.0482358175877709 | 0.518102457705772 | 0.145515302717343 | 0.349616598805252 | 0.145515302717343 | 0.339856215863755 | NA | abundance | 26 | 26 |
| PWY66-429: fatty acid biosynthesis initiation (mitochondria) | group | patient | grouppatient | -0.199600261502717 | 0.0482358175877709 | 0.161094329714222 | 0.14593752492409 | 0.349616598805252 | 0.14593752492409 | 0.339856215863755 | NA | abundance | 26 | 26 |
| PWY-6353: purine nucleotides degradation II (aerobic) | group | patient | grouppatient | 0.347545935190711 | 0.0482358175877709 | 0.196529465674491 | 0.147512059651502 | 0.349658215470227 | 0.147512059651502 | 0.339856215863755 | NA | abundance | 26 | 26 |
| SER-GLYSYN-PWY: superpathway of L-serine and glycine biosynthesis I | group | patient | grouppatient | -0.0811601266450573 | 0.0482358175877709 | 0.0795618884543448 | 0.15126902122566 | 0.355274031502471 | 0.15126902122566 | 0.345128349398354 | NA | abundance | 26 | 26 |
| PWY-5838: superpathway of menaquinol-8 biosynthesis I | group | patient | grouppatient | 0.92251669016063 | 0.0482358175877709 | 0.593008437674783 | 0.156474450728491 | 0.364158721695397 | 0.288464647726199 | 0.526811382008272 | NA | abundance | 26 | 22 |
| P124-PWY: Bifidobacterium shunt | group | patient | grouppatient | -1.04905459861537 | 0.0482358175877709 | 0.771183720864696 | 0.167612236693643 | 0.386565158500653 | 0.167612236693643 | 0.37873918868275 | NA | abundance | 26 | 26 |
| HISDEG-PWY: L-histidine degradation I | group | patient | grouppatient | -0.471712844647278 | 0.0482358175877709 | 0.370196620066072 | 0.17445920147889 | 0.398763889094605 | 0.17445920147889 | 0.38567274579689 | NA | abundance | 26 | 26 |
| PWY-5005: biotin biosynthesis II | group | patient | grouppatient | 0.638755010794127 | 0.0482358175877709 | 0.424840324065974 | 0.178710920862994 | 0.404867218946252 | 0.325484248490288 | 0.562417635258954 | NA | abundance | 26 | 25 |
| PWY-7117: C4 photosynthetic carbon assimilation cycle, PEPCK type | group | patient | grouppatient | 0.526844867118522 | 0.0482358175877709 | 0.357234514519161 | 0.194817538765151 | 0.433680781946771 | 0.194817538765151 | 0.420019464310188 | NA | abundance | 26 | 26 |
| PWY-8187: L-arginine degradation XIII (reductive Stickland reaction) | group | patient | grouppatient | -0.318720686571376 | 0.0482358175877709 | 0.272529347126042 | 0.193980604934943 | 0.433680781946771 | 0.193980604934943 | 0.420019464310188 | NA | abundance | 26 | 26 |
| PWY-5384: sucrose degradation IV (sucrose phosphorylase) | group | patient | grouppatient | 0.36844135066381 | 0.0482358175877709 | 0.241073324063826 | 0.200914513144202 | 0.439607823631758 | 0.200914513144202 | 0.421561701686495 | NA | abundance | 26 | 26 |
| PWY-7400: L-arginine biosynthesis IV (archaebacteria) | group | patient | grouppatient | -0.131774161154519 | 0.0482358175877709 | 0.131321865142276 | 0.199315708486388 | 0.439607823631758 | 0.199315708486388 | 0.421561701686495 | NA | abundance | 26 | 26 |
| GLUCOSE1PMETAB-PWY: glucose and glucose-1-phosphate degradation | group | patient | grouppatient | 3.13127993055332 | 0 | 2.48076076833311 | 0.206867593166368 | 0.448797490259239 | 0.0411445143609233 | 0.163880692793508 | NA | prevalence | 26 | 21 |
| PHOSLIPSYN-PWY: superpathway of phospholipid biosynthesis I (bacteria) | group | patient | grouppatient | 0.285080289371811 | 0.0482358175877709 | 0.180853367581447 | 0.210931986425977 | 0.45376965147101 | 0.210931986425977 | 0.438663865576148 | NA | abundance | 26 | 26 |
| PWY-7345: superpathway of anaerobic sucrose degradation | group | patient | grouppatient | 0.267019172401117 | 0.0482358175877709 | 0.169830738103456 | 0.219204991400463 | 0.459971129496055 | 0.219204991400463 | 0.447940634600947 | NA | abundance | 26 | 26 |
| PWY0-1477: ethanolamine utilization | group | patient | grouppatient | 0.600464786031488 | 0.0482358175877709 | 0.43388796386579 | 0.216385601523096 | 0.459971129496055 | 0.216385601523096 | 0.446058038227435 | NA | abundance | 26 | 26 |
| PWY-5845: superpathway of menaquinol-9 biosynthesis | group | patient | grouppatient | 1.35884687173375 | 0 | 1.1050512259922 | 0.218821203426062 | 0.459971129496055 | 0.174053767577113 | 0.38567274579689 | NA | prevalence | 26 | 19 |
| PWY-5989: stearate biosynthesis II (bacteria and plants) | group | patient | grouppatient | 0.500728394166949 | 0.0482358175877709 | 0.361700644760301 | 0.224836935167403 | 0.464179479055283 | 0.224836935167403 | 0.451595553541365 | NA | abundance | 26 | 26 |
| PWY-8178: pentose phosphate pathway (non-oxidative branch) II | group | patient | grouppatient | -0.143580753529226 | 0.0482358175877709 | 0.150119284792802 | 0.224670667011089 | 0.464179479055283 | 0.224670667011089 | 0.451595553541365 | NA | abundance | 26 | 26 |
| PWY-1861: formaldehyde assimilation II (assimilatory RuMP Cycle) | group | patient | grouppatient | -0.529341610776806 | 0.0482358175877709 | 0.475204221018644 | 0.236148199850129 | 0.481299041876344 | 0.236148199850129 | 0.470295143769324 | NA | abundance | 26 | 26 |
| PWY-5971: palmitate biosynthesis (type II fatty acid synthase) | group | patient | grouppatient | 0.462822964037162 | 0.0482358175877709 | 0.33926727203713 | 0.236889372173513 | 0.481299041876344 | 0.417662169698265 | 0.637566226374194 | NA | abundance | 26 | 25 |
| COLANSYN-PWY: colanic acid building blocks biosynthesis | group | patient | grouppatient | -0.270924531750346 | 0.0482358175877709 | 0.265250892882356 | 0.243607152098397 | 0.486642929886018 | 0.243607152098397 | 0.47349023097295 | NA | abundance | 26 | 26 |
| PWY-6507: 4-deoxy-L-threo-hex-4-enopyranuronate degradation | group | patient | grouppatient | 0.362517265985888 | 0.0482358175877709 | 0.262991480707409 | 0.247826801743289 | 0.486642929886018 | 0.247826801743289 | 0.47349023097295 | NA | abundance | 26 | 26 |
| PWY-5838: superpathway of menaquinol-8 biosynthesis I | group | patient | grouppatient | 2.84536228430117 | 0 | 2.48826692682863 | 0.252826209667345 | 0.486642929886018 | 0.288464647726199 | 0.526811382008272 | NA | prevalence | 26 | 22 |
| PWY-5840: superpathway of menaquinol-7 biosynthesis | group | patient | grouppatient | 2.84536228430117 | 0 | 2.48826692682863 | 0.252826209667345 | 0.486642929886018 | 0.134007988915667 | 0.324658529847235 | NA | prevalence | 26 | 22 |
| PWY-5897: superpathway of menaquinol-11 biosynthesis | group | patient | grouppatient | 2.84536228430117 | 0 | 2.48826692682863 | 0.252826209667345 | 0.486642929886018 | 0.246660204324308 | 0.47349023097295 | NA | prevalence | 26 | 22 |
| PWY-5898: superpathway of menaquinol-12 biosynthesis | group | patient | grouppatient | 2.84536228430117 | 0 | 2.48826692682863 | 0.252826209667345 | 0.486642929886018 | 0.247200392274481 | 0.47349023097295 | NA | prevalence | 26 | 22 |
| PWY-5899: superpathway of menaquinol-13 biosynthesis | group | patient | grouppatient | 2.84536228430117 | 0 | 2.48826692682863 | 0.252826209667345 | 0.486642929886018 | 0.246870968059465 | 0.47349023097295 | NA | prevalence | 26 | 22 |
| PWY0-1061: superpathway of L-alanine biosynthesis | group | patient | grouppatient | -0.715402813976503 | 0.0482358175877709 | 0.665297595819188 | 0.262236721762759 | 0.500989557994525 | 0.262236721762759 | 0.496980883985874 | NA | abundance | 26 | 26 |
| PWY0-845: superpathway of pyridoxal 5'-phosphate biosynthesis and salvage | group | patient | grouppatient | 0.458083685945221 | 0.0482358175877709 | 0.371258237082103 | 0.282049499270006 | 0.534849420837938 | 0.282049499270006 | 0.526811382008272 | NA | abundance | 26 | 26 |
| PWY-702: L-methionine biosynthesis II | group | patient | grouppatient | 0.344481155796164 | 0.0482358175877709 | 0.268544464103824 | 0.284415165436337 | 0.535369723174281 | 0.284415165436337 | 0.526811382008272 | NA | abundance | 26 | 26 |
| COBALSYN-PWY: superpathway of adenosylcobalamin salvage from cobinamide I | group | patient | grouppatient | -0.114511928374492 | 0.0482358175877709 | 0.146232397257834 | 0.289185822464116 | 0.536460656165316 | 0.289185822464116 | 0.526811382008272 | NA | abundance | 26 | 26 |
| PWY66-399: gluconeogenesis III | group | patient | grouppatient | -0.417804292805462 | 0.0482358175877709 | 0.428742885459365 | 0.288289326865537 | 0.536460656165316 | 0.288289326865537 | 0.526811382008272 | NA | abundance | 26 | 26 |
| PWY-6859: all-trans-farnesol biosynthesis | group | patient | grouppatient | 0.718904415949158 | 0.0482358175877709 | 0.631199673554159 | 0.298677835621593 | 0.546153756565199 | 0.298677835621593 | 0.535796117336446 | NA | abundance | 26 | 26 |
| PWY-6901: superpathway of glucose and xylose degradation | group | patient | grouppatient | 0.227499605136677 | 0.0482358175877709 | 0.164414436827085 | 0.297065174972515 | 0.546153756565199 | 0.297065174972515 | 0.535796117336446 | NA | abundance | 26 | 26 |
| PWY-6282: palmitoleate biosynthesis I (from (5Z)-dodec-5-enoate) | group | patient | grouppatient | 0.432643779587552 | 0.0482358175877709 | 0.363911217282989 | 0.302814060274976 | 0.549790066882226 | 0.302814060274976 | 0.539100789125904 | NA | abundance | 26 | 26 |
| METH-ACETATE-PWY: methanogenesis from acetate | group | patient | grouppatient | -0.24794737739891 | 0.0482358175877709 | 0.282501092835354 | 0.307888254622837 | 0.553632155204375 | 0.520981331910977 | 0.724441497035974 | NA | abundance | 26 | 25 |

| | | | | | | | | | | | | | | |
|---|---|---|---|---|---|---|---|---|---|---|---|---|---|---|
| **PWY-5505: L-glutamate and L-glutamine biosynthesis** | group | patient | grouppatient | -0.434166369495915 | 0.0482358175877709 | 0.467888031426574 | 0.312553132891737 | 0.553632155204375 | 0.312553132891737 | 0.552255535560588 | NA | abundance | 26 | 26 |
| **METH-ACETATE-PWY: methanogenesis from acetate** | group | patient | grouppatient | -2.02907657653555 | 0 | 2.02251633158107 | 0.315743338514994 | 0.553632155204375 | 0.520981331910977 | 0.724441497035974 | NA | prevalence | 26 | 25 |
| **P164-PWY: purine nucleobases degradation I (anaerobic)** | group | patient | grouppatient | -2.02907657653555 | 0 | 2.02251633158107 | 0.315743338514994 | 0.553632155204375 | 0.531792821213394 | 0.735125370500868 | NA | prevalence | 26 | 25 |
| **PWY-5971: palmitate biosynthesis (type II fatty acid synthase)** | group | patient | grouppatient | -2.02907657653555 | 0 | 2.02251633158107 | 0.315743338514995 | 0.553632155204375 | 0.417662169698265 | 0.637566226374194 | NA | prevalence | 26 | 25 |
| **GALACT-GLUCUROCAT-PWY: superpathway of hexuronide and hexuronate degradation** | group | patient | grouppatient | 0.38404684525917 | 0.0482358175877709 | 0.332992930629766 | 0.324868856866529 | 0.558163941998869 | 0.324868856866529 | 0.562417635258954 | NA | abundance | 26 | 26 |
| **PWY-6285: superpathway of fatty acids biosynthesis (E. coli)** | group | patient | grouppatient | 0.795072698922361 | 0.0482358175877709 | 0.73337641895094 | 0.324727309174509 | 0.558163941998869 | 0.5440067930253 | 0.743265095121777 | NA | abundance | 26 | 17 |
| **PWY-7237: myo-, chiro- and scyllo-inositol degradation** | group | patient | grouppatient | -0.165679739798071 | 0.0482358175877709 | 0.209371181195377 | 0.321164588822173 | 0.558163941998869 | 0.321164588822173 | 0.562417635258954 | NA | abundance | 26 | 26 |
| **PWY-6936: seleno-amino acid biosynthesis (plants)** | group | patient | grouppatient | 0.204420532375053 | 0.0482358175877709 | 0.154294789645077 | 0.333762278098648 | 0.569620954621693 | 0.333762278098648 | 0.572511936884543 | NA | abundance | 26 | 26 |
| **FASYN-ELONG-PWY: fatty acid elongation -- saturated** | group | patient | grouppatient | 0.379020778600038 | 0.0482358175877709 | 0.345081915967973 | 0.349055312904823 | 0.578297890252821 | 0.349055312904823 | 0.582470721166646 | NA | abundance | 26 | 26 |
| **PWY-3001: superpathway of L-isoleucine biosynthesis I** | group | patient | grouppatient | -0.0162623262329096 | 0.0482358175877709 | 0.0575023670899984 | 0.351533283348308 | 0.578297890252821 | 0.351533283348308 | 0.582470721166646 | NA | abundance | 26 | 26 |
| **PWY-5659: GDP-mannose biosynthesis** | group | patient | grouppatient | -0.205317623640107 | 0.0482358175877709 | 0.268566735421686 | 0.356918229140413 | 0.578297890252821 | 0.356918229140413 | 0.582470721166646 | NA | abundance | 26 | 26 |
| **PWY-7115: C4 photosynthetic carbon assimilation cycle, NAD-ME type** | group | patient | grouppatient | -0.457257964753989 | 0.0482358175877709 | 0.53661927293986 | 0.355716039034448 | 0.578297890252821 | 0.355716039034448 | 0.582470721166646 | NA | abundance | 26 | 26 |
| **PWY-7664: oleate biosynthesis IV (anaerobic)** | group | patient | grouppatient | 0.387866251665144 | 0.0482358175877709 | 0.357910687684891 | 0.353377360654344 | 0.578297890252821 | 0.353377360654344 | 0.582470721166646 | NA | abundance | 26 | 26 |
| **PWY0-862: (5Z)-dodecenoate biosynthesis I** | group | patient | grouppatient | 0.403866177638858 | 0.0482358175877709 | 0.372403469608119 | 0.350800541623694 | 0.578297890252821 | 0.350800541623694 | 0.582470721166646 | NA | abundance | 26 | 26 |
| **THISYN-PWY: superpathway of thiamine diphosphate biosynthesis I** | group | patient | grouppatient | -0.142942748797549 | 0.0482358175877709 | 0.198728162339688 | 0.351282766619543 | 0.578297890252821 | 0.351282766619543 | 0.582470721166646 | NA | abundance | 26 | 26 |
| **PWY-6285: superpathway of fatty acids biosynthesis (E. coli)** | group | patient | grouppatient | 0.850991881548738 | 0 | 0.91263569520397 | 0.35110133919456 | 0.578297890252821 | 0.5440067930253 | 0.743265095121777 | NA | prevalence | 26 | 17 |
| **PWY-7392: taxadiene biosynthesis (engineered)** | group | patient | grouppatient | 0.566298416266173 | 0.0482358175877709 | 0.557815056247547 | 0.362445053348153 | 0.583559331176901 | 0.362445053348153 | 0.587410948529766 | NA | abundance | 26 | 26 |
| **PWY-6519: 8-amino-7-oxononanoate biosynthesis I** | group | patient | grouppatient | 0.342159032202197 | 0.0482358175877709 | 0.321020874722832 | 0.371139775366626 | 0.593280507378887 | 0.371139775366626 | 0.596480867258721 | NA | abundance | 26 | 26 |
| **PYRIDOXSYN-PWY: pyridoxal 5'-phosphate biosynthesis I** | group | patient | grouppatient | 0.41120577766589 | 0.0482358175877709 | 0.399234258476799 | 0.373117819093753 | 0.593280507378887 | 0.373117819093753 | 0.596480867258721 | NA | abundance | 26 | 26 |
| **PWY-8004: Entner-Doudoroff pathway I** | group | patient | grouppatient | 0.258686556690719 | 0.0482358175877709 | 0.232028874391721 | 0.37792567639198 | 0.597215883681153 | 0.37792567639198 | 0.600084688865643 | NA | abundance | 26 | 26 |
| **BIOTIN-BIOSYNTHESIS-PWY: biotin biosynthesis I** | group | patient | grouppatient | 0.304709206035175 | 0.0482358175877709 | 0.293578720669396 | 0.392960796319436 | 0.609670423084086 | 0.392960796319436 | 0.615253601110053 | NA | abundance | 26 | 26 |
| **FASYN-INITIAL-PWY: superpathway of fatty acid biosynthesis initiation (E. coli)** | group | patient | grouppatient | 0.766223241458099 | 0.0482358175877709 | 0.81485223931004 | 0.388383120817889 | 0.609670423084086 | 0.198359747933105 | 0.421561701686495 | NA | abundance | 26 | 22 |
| **PWY-6608: guanosine nucleotides degradation III** | group | patient | grouppatient | 0.244599636600908 | 0.0482358175877709 | 0.223639085547189 | 0.393800582560331 | 0.609670423084086 | 0.393800582560331 | 0.615253601110053 | NA | abundance | 26 | 26 |
| **PWY-7111: pyruvate fermentation to isobutanol (engineered)** | group | patient | grouppatient | -0.0197216994789593 | 0.0482358175877709 | 0.0699633502630989 | 0.395333164968587 | 0.609670423084086 | 0.395333164968587 | 0.615253601110053 | NA | abundance | 26 | 26 |
| **PWY-7234: inosine-5'-phosphate biosynthesis III** | group | patient | grouppatient | 0.362610137967051 | 0.0482358175877709 | 0.368619026356311 | 0.403433667760898 | 0.618437239202335 | 0.403433667760898 | 0.623729683709283 | NA | abundance | 26 | 26 |
| **P164-PWY: purine nucleobases degradation I (anaerobic)** | group | patient | grouppatient | -0.113431527738707 | 0.0482358175877709 | 0.19261373676295 | 0.415006070958658 | 0.632390203365573 | 0.531792821213394 | 0.735125370500868 | NA | abundance | 26 | 25 |
| **GLYCOLYSIS: glycolysis I (from glucose 6-phosphate)** | group | patient | grouppatient | 0.203702199318872 | 0.0482358175877709 | 0.187060159659236 | 0.420522404629788 | 0.633257268148386 | 0.420522404629788 | 0.637566226374194 | NA | abundance | 26 | 26 |
| **PWY-6897: thiamine diphosphate salvage II** | group | patient | grouppatient | 0.141202586009138 | 0.0482358175877709 | 0.106950394143454 | 0.419957044288525 | 0.633257268148386 | 0.419957044288525 | 0.637566226374194 | NA | abundance | 26 | 26 |
| **ANAEROFRUCAT-PWY: homolactic fermentation** | group | patient | grouppatient | 0.173862890017932 | 0.0482358175877709 | 0.157133696204169 | 0.442916920662147 | 0.648008852709846 | 0.442916920662147 | 0.650272858794624 | NA | abundance | 26 | 26 |
| **GLYCOLYSIS-E-D: superpathway of glycolysis and the Entner-Doudoroff pathway** | group | patient | grouppatient | 0.211019084472428 | 0.0482358175877709 | 0.206939251910224 | 0.445415463245393 | 0.648008852709846 | 0.445415463245393 | 0.650272858794624 | NA | abundance | 26 | 26 |
| **NONOXIPENT-PWY: pentose phosphate pathway (non-oxidative branch) I** | group | patient | grouppatient | -0.0574389477694091 | 0.0482358175877709 | 0.130131341392468 | 0.437434354999422 | 0.648008852709846 | 0.437434354999422 | 0.650272858794624 | NA | abundance | 26 | 26 |
| **P4-PWY: superpathway of L-lysine, L-threonine and L-methionine biosynthesis I** | group | patient | grouppatient | 0.49164605541307 | 0.0482358175877709 | 0.570922946485844 | 0.444975090595048 | 0.648008852709846 | 0.444975090595048 | 0.650272858794624 | NA | abundance | 26 | 26 |
| **PWY-6892: thiazole component of thiamine diphosphate biosynthesis I** | group | patient | grouppatient | -0.123882001414553 | 0.0482358175877709 | 0.220537437547816 | 0.445506086238019 | 0.648008852709846 | 0.445506086238019 | 0.650272858794624 | NA | abundance | 26 | 26 |
| **PWY0-1261: anhydromuropeptides recycling I** | group | patient | grouppatient | -0.199866637458552 | 0.0482358175877709 | 0.317007761620565 | 0.442753339702805 | 0.648008852709846 | 0.442753339702805 | 0.650272858794624 | NA | abundance | 26 | 26 |
| **PWY-6895: superpathway of thiamine diphosphate biosynthesis II** | group | patient | grouppatient | 0.231664140389391 | 0.0482358175877709 | 0.236382344763151 | 0.448978763388504 | 0.64937041484439 | 0.448978763388504 | 0.651296354298139 | NA | abundance | 26 | 26 |
| **PWY-6147: 6-hydroxymethyl-dihydropterin diphosphate biosynthesis I** | group | patient | grouppatient | 0.213273518071734 | 0.0482358175877709 | 0.214361031217912 | 0.453727753100781 | 0.652552274122471 | 0.453727753100781 | 0.654147374102352 | NA | abundance | 26 | 26 |
| **PWY0-781: aspartate superpathway** | group | patient | grouppatient | 0.475714595758472 | 0.0482358175877709 | 0.564833903403426 | 0.456594124055243 | 0.653006121553867 | 0.456594124055243 | 0.654265970445013 | NA | abundance | 26 | 26 |
| **PWY-7357: thiamine phosphate formation from pyrithiamine and oxythiamine (yeast)** | group | patient | grouppatient | -0.0237030874825036 | 0.0482358175877709 | 0.0905377877324706 | 0.465941728119073 | 0.662672679991571 | 0.465941728119073 | 0.663613976412014 | NA | abundance | 26 | 26 |
| **PWY-5484: glycolysis II (from fructose 6-phosphate)** | group | patient | grouppatient | 0.187715384904276 | 0.0482358175877709 | 0.188499258952063 | 0.473581985313119 | 0.669817614586511 | 0.473581985313119 | 0.670432328605922 | NA | abundance | 26 | 26 |
| **GLYCOCAT-PWY: glycogen degradation I** | group | patient | grouppatient | 0.337921031628628 | 0.0482358175877709 | 0.415828109971567 | 0.493223624986773 | 0.689974032768382 | 0.493223624986773 | 0.694057196837675 | NA | abundance | 26 | 26 |
| **PWY-6168: flavin biosynthesis III (fungi)** | group | patient | grouppatient | 0.512980844337478 | 0.0482358175877709 | 0.664921064222185 | 0.492059435564312 | 0.689974032768382 | 0.741996383000755 | 0.858961330074765 | NA | abundance | 26 | 23 |
| **PWY-7761: NAD salvage pathway II (PNC IV cycle)** | group | patient | grouppatient | 0.301214379023713 | 0.0482358175877709 | 0.376669202757369 | 0.509125626742101 | 0.708348698075966 | 0.509125626742101 | 0.712169775502344 | NA | abundance | 26 | 26 |
| **FUC-RHAMCAT-PWY: superpathway of fucose and rhamnose degradation** | group | patient | grouppatient | -0.20444444722982 | 0.0482358175877709 | 0.443666571877448 | 0.574394824712488 | 0.749360595700482 | 0.574394824712488 | 0.765617248807243 | NA | abundance | 26 | 26 |
| **P441-PWY: superpathway of N-acetylneuraminate degradation** | group | patient | grouppatient | 0.199702581939154 | 0.0482358175877709 | 0.261407290125491 | 0.569025557940628 | 0.749360595700482 | 0.569025557940628 | 0.764120034948844 | NA | abundance | 26 | 26 |
| **PWY-1269: CMP-3-deoxy-D-manno-octulosonate biosynthesis** | group | patient | grouppatient | -0.120689369449675 | 0.0482358175877709 | 0.297942014576599 | 0.576656395910136 | 0.749360595700482 | 0.576656395910136 | 0.765617248807243 | NA | abundance | 26 | 26 |
| **PWY-6124: inosine-5'-phosphate biosynthesis II** | group | patient | grouppatient | 0.106115143555163 | 0.0482358175877709 | 0.0890250115403336 | 0.553175724032831 | 0.749360595700482 | 0.553175724032831 | 0.747105144527099 | NA | abundance | 26 | 26 |
| **PWY-621: sucrose degradation III (sucrose invertase)** | group | patient | grouppatient | 0.153159624185842 | 0.0482358175877709 | 0.166918163190718 | 0.543976580606623 | 0.749360595700482 | 0.543976580606623 | 0.743265095121777 | NA | abundance | 26 | 26 |
| **PWY0-1586: peptidoglycan maturation (meso-diaminopimelate containing)** | group | patient | grouppatient | 0.0984347248432019 | 0.0482358175877709 | 0.0744698412023128 | 0.553148770026826 | 0.749360595700482 | 0.553148770026826 | 0.747105144527099 | NA | abundance | 26 | 26 |

| | | | | | | | | | | | | | | |
|---|---|---|---|---|---|---|---|---|---|---|---|---|---|---|
| **PWY-5005: biotin biosynthesis II** | group | patient | grouppatient | 1.46033919957257 | 0 | 2.58847384168459 | 0.572638465549701 | 0.749360595700482 | 0.325484248490288 | 0.562417635258954 | NA | prevalence | 26 | 25 |
| **PWY-5981: CDP-diacylglycerol biosynthesis III** | group | patient | grouppatient | 1.46033919957257 | 0 | 2.58847384168459 | 0.572638465549701 | 0.749360595700482 | 0.04846376146895 | 0.177952874143801 | NA | prevalence | 26 | 25 |
| **PWY-6545: pyrimidine deoxyribonucleotides de novo biosynthesis III** | group | patient | grouppatient | 1.46033919957257 | 0 | 2.58847384168459 | 0.572638465549701 | 0.749360595700482 | 0.817362118872286 | 0.898283823082983 | NA | prevalence | 26 | 25 |
| **PWY-7184: pyrimidine deoxyribonucleotides de novo biosynthesis I** | group | patient | grouppatient | 1.46033919957257 | 0 | 2.58847384168459 | 0.572638465549701 | 0.749360595700482 | 0.817362118872286 | 0.898283823082983 | NA | prevalence | 26 | 25 |
| **PWY-7185: UTP and CTP dephosphorylation I** | group | patient | grouppatient | 1.46033919957257 | 0 | 2.58847384168459 | 0.572638465549701 | 0.749360595700482 | 0.817362118872286 | 0.898283823082983 | NA | prevalence | 26 | 25 |
| **PWY-7210: pyrimidine deoxyribonucleotides biosynthesis from CTP** | group | patient | grouppatient | 1.46033919957257 | 0 | 2.58847384168459 | 0.572638465549701 | 0.749360595700482 | 0.817362118872286 | 0.898283823082983 | NA | prevalence | 26 | 25 |
| **PWY-7211: superpathway of pyrimidine deoxyribonucleotides de novo biosynthesis** | group | patient | grouppatient | 1.46033919957257 | 0 | 2.58847384168459 | 0.572638465549701 | 0.749360595700482 | 0.817362118872286 | 0.898283823082983 | NA | prevalence | 26 | 25 |
| **GALACTUROCAT-PWY: D-galacturonate degradation I** | group | patient | grouppatient | 0.202414956117818 | 0.0482358175877709 | 0.288726456238718 | 0.599709668723803 | 0.768652312632075 | 0.599709668723803 | 0.783998669568306 | NA | abundance | 26 | 26 |
| **PWY-6703: preQ0 biosynthesis** | group | patient | grouppatient | -0.0404443379147391 | 0.0482358175877709 | 0.163699741181038 | 0.598224673155841 | 0.768652312632075 | 0.598224673155841 | 0.783998669568306 | NA | abundance | 26 | 26 |
| **PWY-I9: L-cysteine biosynthesis VI (from L-methionine)** | group | patient | grouppatient | 0.213249642800838 | 0.0482358175877709 | 0.310049302720633 | 0.600509619243809 | 0.768652312632075 | 0.600509619243809 | 0.783998669568306 | NA | abundance | 26 | 26 |
| **PWY-6902: chitin degradation II (Vibrio)** | group | patient | grouppatient | 0.231749006894837 | 0.0482358175877709 | 0.352915375868752 | 0.608623918907542 | 0.7751628021907 | 0.608623918907542 | 0.790202325653439 | NA | abundance | 26 | 26 |
| **PWY-6628: superpathway of L-phenylalanine biosynthesis** | group | patient | grouppatient | 0.120195897385079 | 0.0482358175877709 | 0.137223677888974 | 0.614972305005135 | 0.779370841986706 | 0.614972305005135 | 0.793698989626825 | NA | abundance | 26 | 26 |
| **PWY-5497: purine nucleobases degradation II (anaerobic)** | group | patient | grouppatient | 0.251141417304275 | 0.0482358175877709 | 0.401511925054217 | 0.618071979156208 | 0.779440525438371 | 0.618071979156208 | 0.793698989626825 | NA | abundance | 26 | 26 |
| **ARGININE-SYN4-PWY: L-ornithine biosynthesis II** | group | patient | grouppatient | 0.255556807565672 | 0.0482358175877709 | 0.455145381437409 | 0.653136049134701 | 0.809584822648586 | 0.653136049134701 | 0.82708026078924 | NA | abundance | 26 | 26 |
| **PWY-7210: pyrimidine deoxyribonucleotides biosynthesis from CTP** | group | patient | grouppatient | -0.373671759138194 | 0.0482358175877709 | 0.93163316475672 | 0.654384200514092 | 0.809584822648586 | 0.817362118872286 | 0.898283823082983 | NA | abundance | 26 | 25 |
| **PWY-7220: adenosine deoxyribonucleotides de novo biosynthesis II** | group | patient | grouppatient | 0.147753965640738 | 0.0482358175877709 | 0.217506065534293 | 0.654172511141769 | 0.809584822648586 | 0.654172511141769 | 0.82708026078924 | NA | abundance | 26 | 26 |
| **PWY-7222: guanosine deoxyribonucleotides de novo biosynthesis II** | group | patient | grouppatient | 0.147753965640738 | 0.0482358175877709 | 0.217506065534293 | 0.654625227688505 | 0.809584822648586 | 0.654625227688505 | 0.82708026078924 | NA | abundance | 26 | 26 |
| **CITRULBIO-PWY: L-citrulline biosynthesis** | group | patient | grouppatient | 0.207656039673303 | 0.0482358175877709 | 0.368594052798331 | 0.669315056249369 | 0.809699599763794 | 0.669315056249369 | 0.828892207130643 | NA | abundance | 26 | 26 |
| **DAPLYSINESYN-PWY: L-lysine biosynthesis I** | group | patient | grouppatient | -0.142381837745413 | 0.0482358175877709 | 0.444036004307863 | 0.671717583141082 | 0.809699599763794 | 0.671717583141082 | 0.828892207130643 | NA | abundance | 26 | 26 |
| **PWY-4984: urea cycle** | group | patient | grouppatient | 0.21364448778755 | 0.0482358175877709 | 0.387866015322307 | 0.673695368348735 | 0.809699599763794 | 0.673695368348735 | 0.828892207130643 | NA | abundance | 26 | 26 |
| **PWY-6317: D-galactose degradation I (Leloir pathway)** | group | patient | grouppatient | 0.0823716612997123 | 0.0482358175877709 | 0.0677324795579928 | 0.66335010483376 | 0.809699599763794 | 0.66335010483376 | 0.828892207130643 | NA | abundance | 26 | 26 |
| **PWY-7560: methylerythritol phosphate pathway II** | group | patient | grouppatient | 0.097815879235421 | 0.0482358175877709 | 0.106609915058364 | 0.662297189187871 | 0.809699599763794 | 0.662297189187871 | 0.828892207130643 | NA | abundance | 26 | 26 |
| **PWY-622: starch biosynthesis** | group | patient | grouppatient | -0.568737376960044 | 0 | 1.33778991366626 | 0.670740410510824 | 0.809699599763794 | 0.175604186384116 | 0.38567274579689 | NA | prevalence | 26 | 24 |
| **PWY-5030: L-histidine degradation III** | group | patient | grouppatient | -0.180789999246395 | 0.0482358175877709 | 0.55221585519099 | 0.681821886632165 | 0.815637397092683 | 0.681821886632165 | 0.834521579992494 | NA | abundance | 26 | 26 |
| **PWY-7208: superpathway of pyrimidine nucleobases salvage** | group | patient | grouppatient | 0.12131229876324 | 0.0482358175877709 | 0.175996457900715 | 0.686380915392172 | 0.817272159722772 | 0.686380915392172 | 0.835748782990468 | NA | abundance | 26 | 26 |
| **NAGLIPASYN-PWY: lipid IVA biosynthesis (E. coli)** | group | patient | grouppatient | 0.218765893759444 | 0.0482358175877709 | 0.4328806692253 | 0.697053107274864 | 0.818557777350299 | 0.697053107274864 | 0.840038360049195 | NA | abundance | 26 | 26 |
| **PWY-7211: superpathway of pyrimidine deoxyribonucleotides de novo biosynthesis** | group | patient | grouppatient | -0.264038456603496 | 0.0482358175877709 | 0.790054335988786 | 0.695992981663738 | 0.818557777350299 | 0.817362118872286 | 0.898283823082983 | NA | abundance | 26 | 25 |
| **PWY-8073: lipid IVA biosynthesis (P. putida)** | group | patient | grouppatient | 0.218765893759444 | 0.0482358175877709 | 0.4328806692253 | 0.696983546521302 | 0.818557777350299 | 0.696983546521302 | 0.840038360049195 | NA | abundance | 26 | 26 |
| **GLCMANNANAUT-PWY: superpathway of N-acetylglucosamine, N-acetylmannosamine and N-acetyl** | group | patient | grouppatient | -0.021320086175841 | 0.0482358175877709 | 0.193991313198873 | 0.725962619158407 | 0.838403443212293 | 0.725962619158407 | 0.857259089956876 | NA | abundance | 26 | 26 |
| **PWY-7242: D-fructuronate degradation** | group | patient | grouppatient | 0.140736801283439 | 0.0482358175877709 | 0.270696906148188 | 0.736623387725378 | 0.838403443212293 | 0.736623387725378 | 0.857259089956876 | NA | abundance | 26 | 26 |
| **PWY-7328: superpathway of UDP-glucose-derived O-antigen building blocks biosynthesis** | group | patient | grouppatient | -0.0094271032495489 | 0.0482358175877709 | 0.164220030216853 | 0.73546701350296 | 0.838403443212293 | 0.73546701350296 | 0.857259089956876 | NA | abundance | 26 | 26 |
| **PWY-7356: thiamine diphosphate salvage IV (yeast)** | group | patient | grouppatient | 0.179346379530265 | 0.0482358175877709 | 0.361924911970662 | 0.720426226135731 | 0.838403443212293 | 0.720426226135731 | 0.857259089956876 | NA | abundance | 26 | 26 |
| **PWY-7663: gondoate biosynthesis (anaerobic)** | group | patient | grouppatient | 0.0949450050452542 | 0.0482358175877709 | 0.133462400460965 | 0.736878026260804 | 0.838403443212293 | 0.736878026260804 | 0.857259089956876 | NA | abundance | 26 | 26 |
| **PWY0-162: superpathway of pyrimidine ribonucleotides de novo biosynthesis** | group | patient | grouppatient | 0.0039153260243046 | 0.0482358175877709 | 0.122864792174498 | 0.733823074127899 | 0.838403443212293 | 0.733823074127899 | 0.857259089956876 | NA | abundance | 26 | 26 |
| **UDPNAGSYN-PWY: UDP-N-acetyl-D-glucosamine biosynthesis I** | group | patient | grouppatient | -0.0309296260301664 | 0.0482358175877709 | 0.223988462558673 | 0.728684144826606 | 0.838403443212293 | 0.728684144826606 | 0.857259089956876 | NA | abundance | 26 | 26 |
| **PWY-6123: inosine-5'-phosphate biosynthesis I** | group | patient | grouppatient | 0.0725690568777415 | 0.0482358175877709 | 0.0790326055938895 | 0.781623251426018 | 0.885378550287879 | 0.781623251426018 | 0.898283823082983 | NA | abundance | 26 | 26 |
| **PWY-241: C4 photosynthetic carbon assimilation cycle, NADP-ME type** | group | patient | grouppatient | -0.067528242817846 | 0.0482358175877709 | 0.431272851322762 | 0.790641920238239 | 0.891649037801714 | 0.790641920238239 | 0.898283823082983 | NA | abundance | 26 | 26 |
| **PWY-6545: pyrimidine deoxyribonucleotides de novo biosynthesis III** | group | patient | grouppatient | -0.181642532470819 | 0.0482358175877709 | 0.876549106771768 | 0.795171087089112 | 0.892823676731634 | 0.817362118872286 | 0.898283823082983 | NA | abundance | 26 | 25 |
| **PWY-6731: starch degradation III** | group | patient | grouppatient | -0.0199764620195291 | 0.0482358175877709 | 0.269229463997274 | 0.802743123481785 | 0.897389692625926 | 0.802743123481785 | 0.898283823082983 | NA | abundance | 26 | 26 |
| **PWY-7198: pyrimidine deoxyribonucleotides de novo biosynthesis IV** | group | patient | grouppatient | 0.105826023763179 | 0.0482358175877709 | 0.232335039607236 | 0.807534879206931 | 0.898821430769454 | 0.807534879206931 | 0.898283823082983 | NA | abundance | 26 | 26 |
| **PWY-6270: isoprene biosynthesis I** | group | patient | grouppatient | 0.0262861318183705 | 0.0482358175877709 | 0.0865903526317355 | 0.816016923544824 | 0.902959394073371 | 0.816016923544824 | 0.898283823082983 | NA | abundance | 26 | 26 |
| **PWY-7229: superpathway of adenosine nucleotides de novo biosynthesis I** | group | patient | grouppatient | 0.0151741822165766 | 0.0482358175877709 | 0.139571880437087 | 0.819679562027325 | 0.902959394073371 | 0.819679562027325 | 0.898283823082983 | NA | abundance | 26 | 26 |
| **PWY-841: superpathway of purine nucleotides de novo biosynthesis I** | group | patient | grouppatient | 0.0159521122424989 | 0.0482358175877709 | 0.1380009864048 | 0.821834136012091 | 0.902959394073371 | 0.821834136012091 | 0.898283823082983 | NA | abundance | 26 | 26 |
| **FUCCAT-PWY: fucose degradation** | group | patient | grouppatient | 0.116558432470342 | 0.0482358175877709 | 0.340172993338833 | 0.842840865459421 | 0.922082314348768 | 0.842840865459421 | 0.9169796452915 | NA | abundance | 26 | 26 |
| **PWY-5154: L-arginine biosynthesis III (via N-acetyl-L-citrulline)** | group | patient | grouppatient | 0.106791821619706 | 0.0482358175877709 | 0.304814613509529 | 0.849552189390991 | 0.92546961908125 | 0.849552189390991 | 0.920021956252917 | NA | abundance | 26 | 26 |
| **PWY-6125: superpathway of guanosine nucleotides de novo biosynthesis II** | group | patient | grouppatient | 0.0834128825749981 | 0.0482358175877709 | 0.19724778098012 | 0.861583924941667 | 0.930212719958028 | 0.861583924941667 | 0.927823755660637 | NA | abundance | 26 | 26 |
| **PWY-7184: pyrimidine deoxyribonucleotides de novo biosynthesis I** | group | patient | grouppatient | -0.110953579401978 | 0.0482358175877709 | 0.89465139621856 | 0.860185809097925 | 0.930212719958028 | 0.817362118872286 | 0.898283823082983 | NA | abundance | 26 | 25 |
| **PWY-7323: superpathway of GDP-mannose-derived O-antigen building blocks biosynthesis** | group | patient | grouppatient | -0.0091515203734062 | 0.0482358175877709 | 0.331248862364514 | 0.864807138085979 | 0.930212719958028 | 0.864807138085979 | 0.927823755660637 | NA | abundance | 26 | 26 |

| | | | | | | | | | | | | | | |
|---|---|---|---|---|---|---|---|---|---|---|---|---|---|---|
| **THRESYN-PWY: superpathway of L-threonine biosynthesis** | group | patient | grouppatient | 0.0615258787858898 | 0.0482358175877709 | 0.0710131282075294 | 0.868600962746128 | 0.930384294824305 | 0.868600962746128 | 0.927823755660637 | NA | abundance | 26 | 26 |
| **PWY-6470: peptidoglycan biosynthesis V (β-lactam resistance)** | group | patient | grouppatient | 0.00465398139017357 | 0.0482358175877709 | 0.267168790830905 | 0.873130190968199 | 0.931338870366079 | 0.873130190968199 | 0.928441605780664 | NA | abundance | 26 | 26 |
| **GLUCUROCAT-PWY: superpathway of β-D-glucuronosides degradation** | group | patient | grouppatient | 0.0799320899110569 | 0.0482358175877709 | 0.26483380487789 | 0.906040710118769 | 0.942871633294328 | 0.906040710118769 | 0.937971660255113 | NA | abundance | 26 | 26 |
| **PWY-5345: superpathway of L-methionine biosynthesis (by sulfhydrylation)** | group | patient | grouppatient | 0.125920252817264 | 0.0482358175877709 | 0.65114301637264 | 0.9059100722294 | 0.942871633294328 | 0.9059100722294 | 0.937971660255113 | NA | abundance | 26 | 26 |
| **PWY-6292: superpathway of L-cysteine biosynthesis (mammalian)** | group | patient | grouppatient | 0.0217135573738285 | 0.0482358175877709 | 0.183909938347513 | 0.887903121317938 | 0.942871633294328 | 0.887903121317938 | 0.937971660255113 | NA | abundance | 26 | 26 |
| **PWY-6305: superpathway of putrescine biosynthesis** | group | patient | grouppatient | 0.0956822533992318 | 0.0482358175877709 | 0.359373211659471 | 0.896049372885493 | 0.942871633294328 | 0.896049372885493 | 0.937971660255113 | NA | abundance | 26 | 26 |
| **PWY-7228: superpathway of guanosine nucleotides de novo biosynthesis I** | group | patient | grouppatient | 0.0727410927121824 | 0.0482358175877709 | 0.180861530605082 | 0.894736422091422 | 0.942871633294328 | 0.894736422091422 | 0.937971660255113 | NA | abundance | 26 | 26 |
| **SULFATE-CYS-PWY: superpathway of sulfate assimilation and cysteine biosynthesis** | group | patient | grouppatient | 0.126729019368499 | 0.0482358175877709 | 0.655417815540951 | 0.905521442879331 | 0.942871633294328 | 0.905521442879331 | 0.937971660255113 | NA | abundance | 26 | 26 |
| **GLUCONEO-PWY: gluconeogenesis I** | group | patient | grouppatient | 0.0629382887772096 | 0.0482358175877709 | 0.134026531900318 | 0.916096799358517 | 0.948750606194299 | 0.916096799358517 | 0.943183428783497 | NA | abundance | 26 | 26 |
| **PWY-7197: pyrimidine deoxyribonucleotide phosphorylation** | group | patient | grouppatient | 0.0703304248654548 | 0.0482358175877709 | 0.213464858951027 | 0.919102149750727 | 0.948750606194299 | 0.919102149750727 | 0.943183428783497 | NA | abundance | 26 | 26 |
| **P41-PWY: pyruvate fermentation to acetate and (S)-lactate I** | group | patient | grouppatient | 0.0417413655127867 | 0.0482358175877709 | 0.113949588491633 | 0.956847142114468 | 0.975907842156588 | 0.956847142114468 | 0.973415923796103 | NA | abundance | 26 | 26 |
| **PWY-5100: pyruvate fermentation to acetate and lactate II** | group | patient | grouppatient | 0.0417413655127867 | 0.0482358175877709 | 0.113949588491633 | 0.956827944772579 | 0.975907842156588 | 0.956827944772579 | 0.973415923796103 | NA | abundance | 26 | 26 |
| **PWY-7185: UTP and CTP dephosphorylation I** | group | patient | grouppatient | -0.02086480510376 | 0.0482358175877709 | 1.09869501082136 | 0.950423519030826 | 0.975907842156588 | 0.817362118872286 | 0.898283823082983 | NA | abundance | 26 | 25 |
| **PWY-6168: flavin biosynthesis III (fungi)** | group | patient | grouppatient | 0.0561511714507715 | 0 | 1.21234981226173 | 0.963058401600399 | 0.978345042895644 | 0.741996383000755 | 0.858961330074765 | NA | prevalence | 26 | 23 |
| **PWY-6126: superpathway of adenosine nucleotides de novo biosynthesis II** | group | patient | grouppatient | 0.0444272130666948 | 0.0482358175877709 | 0.176112357696323 | 0.983165222402876 | 0.994823308044017 | 0.983165222402876 | 0.995878565796017 | NA | abundance | 26 | 26 |
| **PWY-5973: cis-vaccenate biosynthesis** | group | patient | grouppatient | 0.0467991400266427 | 0.0482358175877709 | 0.14189686055356 | 0.992190327573233 | 0.996081270034305 | 0.992190327573233 | 0.996430457178247 | NA | abundance | 26 | 26 |
| **PWY0-1479: tRNA processing** | group | patient | grouppatient | 0.0523811167051743 | 0.0482358175877709 | 0.385335887604935 | 0.991512237494714 | 0.996081270034305 | 0.991512237494714 | 0.996430457178247 | NA | abundance | 26 | 26 |
| **PWY-5913: partial TCA cycle (obligate autotrophs)** | group | patient | grouppatient | 0.0496724951488991 | 0.0482358175877709 | 0.328854447325851 | 0.996554402095212 | 0.996554402095212 | 0.996554402095212 | 0.996554402095212 | NA | abundance | 26 | 26 |
| **1CMET2-PWY: folate transformations III (E. coli)** | group | patient | grouppatient | NA | 0 | NA | NA | NA | 0.0276953127956533 | 0.132824459326092 | All logi | prevalence | 26 | 26 |
| **ANAEROFRUCAT-PWY: homolactic fermentation** | group | patient | grouppatient | NA | 0 | NA | NA | NA | 0.442916920662147 | 0.650272858794624 | All logi | prevalence | 26 | 26 |
| **ANAGLYCOLYSIS-PWY: glycolysis III (from glucose)** | group | patient | grouppatient | NA | 0 | NA | NA | NA | 0.113632574596847 | 0.300940506011959 | All logi | prevalence | 26 | 26 |
| **ARG+POLYAMINE-SYN: superpathway of arginine and polyamine biosynthesis** | group | patient | grouppatient | NA | 0 | NA | NA | NA | 0.0679150519494542 | 0.205872735384621 | All logi | prevalence | 26 | 26 |
| **ARGININE-SYN4-PWY: L-ornithine biosynthesis II** | group | patient | grouppatient | NA | 0 | NA | NA | NA | 0.653136049134701 | 0.82708026078924 | All logi | prevalence | 26 | 26 |
| **ARGSYN-PWY: L-arginine biosynthesis I (via L-ornithine)** | group | patient | grouppatient | NA | 0 | NA | NA | NA | 7.09537826051321e-05 | 0.00734183923282705 | All logi | prevalence | 26 | 26 |
| **ARGSYNBSUB-PWY: L-arginine biosynthesis II (acetyl cycle)** | group | patient | grouppatient | NA | 0 | NA | NA | NA | 0.000156209345379299 | 0.00734183923282705 | All logi | prevalence | 26 | 26 |
| **ARO-PWY: chorismate biosynthesis I** | group | patient | grouppatient | NA | 0 | NA | NA | NA | 0.00104569783115105 | 0.0252289488287085 | All logi | prevalence | 26 | 26 |
| **ASPASN-PWY: superpathway of L-aspartate and L-asparagine biosynthesis** | group | patient | grouppatient | NA | 0 | NA | NA | NA | 0.0683322270638316 | 0.205872735384621 | All logi | prevalence | 26 | 26 |
| **BIOTIN-BIOSYNTHESIS-PWY: biotin biosynthesis I** | group | patient | grouppatient | NA | 0 | NA | NA | NA | 0.392960796319436 | 0.615253601110053 | All logi | prevalence | 26 | 26 |
| **BRANCHED-CHAIN-AA-SYN-PWY: superpathway of branched chain amino acid biosynthesis** | group | patient | grouppatient | NA | 0 | NA | NA | NA | 0.0100152600160031 | 0.0735495657425225 | All logi | prevalence | 26 | 26 |
| **CALVIN-PWY: Calvin-Benson-Bassham cycle** | group | patient | grouppatient | NA | 0 | NA | NA | NA | 0.0160951816426669 | 0.0958791043121336 | All logi | prevalence | 26 | 26 |
| **CENTFERM-PWY: pyruvate fermentation to butanoate** | group | patient | grouppatient | NA | 0 | NA | NA | NA | 0.0323664875534683 | 0.1376771299747 | All logi | prevalence | 26 | 26 |
| **CITRULBIO-PWY: L-citrulline biosynthesis** | group | patient | grouppatient | NA | 0 | NA | NA | NA | 0.669315056249369 | 0.828892207130643 | All logi | prevalence | 26 | 26 |
| **COA-PWY-1: superpathway of coenzyme A biosynthesis III (mammals)** | group | patient | grouppatient | NA | 0 | NA | NA | NA | 0.00319503579138125 | 0.0362720921339909 | All logi | prevalence | 26 | 26 |
| **COA-PWY: coenzyme A biosynthesis I (prokaryotic)** | group | patient | grouppatient | NA | 0 | NA | NA | NA | 0.00407965550692002 | 0.0368738093894694 | All logi | prevalence | 26 | 26 |
| **COBALSYN-PWY: superpathway of adenosylcobalamin salvage from cobinamide I** | group | patient | grouppatient | NA | 0 | NA | NA | NA | 0.289185822464116 | 0.526811382008272 | All logi | prevalence | 26 | 26 |
| **COLANSYN-PWY: colanic acid building blocks biosynthesis** | group | patient | grouppatient | NA | 0 | NA | NA | NA | 0.243607152098397 | 0.47349023097295 | All logi | prevalence | 26 | 26 |
| **COMPLETE-ARO-PWY: superpathway of aromatic amino acid biosynthesis** | group | patient | grouppatient | NA | 0 | NA | NA | NA | 0.00129281963359129 | 0.0252289488287085 | All logi | prevalence | 26 | 26 |
| **DAPLYSINESYN-PWY: L-lysine biosynthesis I** | group | patient | grouppatient | NA | 0 | NA | NA | NA | 0.671717583141082 | 0.828892207130643 | All logi | prevalence | 26 | 26 |
| **DTDPRHAMSYN-PWY: dTDP-β-L-rhamnose biosynthesis** | group | patient | grouppatient | NA | 0 | NA | NA | NA | 0.101366996146688 | 0.278461278780288 | All logi | prevalence | 26 | 26 |
| **FASYN-ELONG-PWY: fatty acid elongation -- saturated** | group | patient | grouppatient | NA | 0 | NA | NA | NA | 0.349055312904823 | 0.582470721166646 | All logi | prevalence | 26 | 26 |
| **FERMENTATION-PWY: mixed acid fermentation** | group | patient | grouppatient | NA | 0 | NA | NA | NA | 0.00208031964149091 | 0.0325916743833576 | All logi | prevalence | 26 | 26 |
| **FOLSYN-PWY: superpathway of tetrahydrofolate biosynthesis and salvage** | group | patient | grouppatient | NA | 0 | NA | NA | NA | 0.01806960191275 | 0.103569669499908 | All logi | prevalence | 26 | 26 |
| **FUC-RHAMCAT-PWY: superpathway of fucose and rhamnose degradation** | group | patient | grouppatient | NA | 0 | NA | NA | NA | 0.574394824712488 | 0.765617248807243 | All logi | prevalence | 26 | 26 |
| **FUCCAT-PWY: fucose degradation** | group | patient | grouppatient | NA | 0 | NA | NA | NA | 0.842840865459421 | 0.9169796452915 | All logi | prevalence | 26 | 26 |
| **GALACT-GLUCUROCAT-PWY: superpathway of hexuronide and hexuronate degradation** | group | patient | grouppatient | NA | 0 | NA | NA | NA | 0.324868856866529 | 0.562417635258954 | All logi | prevalence | 26 | 26 |
| **GALACTUROCAT-PWY: D-galacturonate degradation I** | group | patient | grouppatient | NA | 0 | NA | NA | NA | 0.599709668723803 | 0.783998669568306 | All logi | prevalence | 26 | 26 |
| **GLCMANNANAUT-PWY: superpathway of N-acetylglucosamine, N-acetylmannosamine and N-acetyl** | group | patient | grouppatient | NA | 0 | NA | NA | NA | 0.725962619158407 | 0.857259089956876 | All logi | prevalence | 26 | 26 |
| **GLUCONEO-PWY: gluconeogenesis I** | group | patient | grouppatient | NA | 0 | NA | NA | NA | 0.916096799358517 | 0.943183428783497 | All logi | prevalence | 26 | 26 |

| | | | | | | | | | | | | | | |
|---|---|---|---|---|---|---|---|---|---|---|---|---|---|---|
| **GLUCUROCAT-PWY: superpathway of β-D-glucuronosides degradation** | group | patient | grouppatient | NA | 0 | NA | NA | NA | 0.906040710118769 | 0.937971660255113 | All logi | prevalence | 26 | 26 |
| **GLUTORN-PWY: L-ornithine biosynthesis I** | group | patient | grouppatient | NA | 0 | NA | NA | NA | 0.000493851056067918 | 0.0193424996959934 | All logi | prevalence | 26 | 26 |
| **GLYCOCAT-PWY: glycogen degradation I** | group | patient | grouppatient | NA | 0 | NA | NA | NA | 0.493223624986773 | 0.694057196837675 | All logi | prevalence | 26 | 26 |
| **GLYCOGENSYNTH-PWY: glycogen biosynthesis I (from ADP-D-Glucose)** | group | patient | grouppatient | NA | 0 | NA | NA | NA | 0.000880147684649168 | 0.0252289488287085 | All logi | prevalence | 26 | 26 |
| **GLYCOLYSIS-E-D: superpathway of glycolysis and the Entner-Doudoroff pathway** | group | patient | grouppatient | NA | 0 | NA | NA | NA | 0.445415463245393 | 0.650272858794624 | All logi | prevalence | 26 | 26 |
| **GLYCOLYSIS: glycolysis I (from glucose 6-phosphate)** | group | patient | grouppatient | NA | 0 | NA | NA | NA | 0.420522404629788 | 0.637566226374194 | All logi | prevalence | 26 | 26 |
| **HISDEG-PWY: L-histidine degradation I** | group | patient | grouppatient | NA | 0 | NA | NA | NA | 0.17445920147889 | 0.38567274579689 | All logi | prevalence | 26 | 26 |
| **HISTSYN-PWY: L-histidine biosynthesis** | group | patient | grouppatient | NA | 0 | NA | NA | NA | 0.00015558987309916 | 0.00734183923282705 | All logi | prevalence | 26 | 26 |
| **HSERMETANA-PWY: L-methionine biosynthesis III** | group | patient | grouppatient | NA | 0 | NA | NA | NA | 0.023418327647841 | 0.118411940685157 | All logi | prevalence | 26 | 26 |
| **ILEUSYN-PWY: L-isoleucine biosynthesis I (from threonine)** | group | patient | grouppatient | NA | 0 | NA | NA | NA | 0.0153889532577307 | 0.0958791043121336 | All logi | prevalence | 26 | 26 |
| **LACTOSECAT-PWY: lactose and galactose degradation I** | group | patient | grouppatient | NA | 0 | NA | NA | NA | 0.0236823881370314 | 0.118411940685157 | All logi | prevalence | 26 | 26 |
| **MET-SAM-PWY: superpathway of S-adenosyl-L-methionine biosynthesis** | group | patient | grouppatient | NA | 0 | NA | NA | NA | 0.120145566946195 | 0.303593636907051 | All logi | prevalence | 26 | 26 |
| **METSYN-PWY: superpathway of L-homoserine and L-methionine biosynthesis** | group | patient | grouppatient | NA | 0 | NA | NA | NA | 0.115602597027463 | 0.300940506011959 | All logi | prevalence | 26 | 26 |
| **NAGLIPASYN-PWY: lipid IVA biosynthesis (E. coli)** | group | patient | grouppatient | NA | 0 | NA | NA | NA | 0.697053107274864 | 0.840038360049195 | All logi | prevalence | 26 | 26 |
| **NONMEVIPP-PWY: methylerythritol phosphate pathway I** | group | patient | grouppatient | NA | 0 | NA | NA | NA | 8.61377185228385e-05 | 0.00734183923282705 | All logi | prevalence | 26 | 26 |
| **NONOXIPENT-PWY: pentose phosphate pathway (non-oxidative branch) I** | group | patient | grouppatient | NA | 0 | NA | NA | NA | 0.437434354999422 | 0.650272858794624 | All logi | prevalence | 26 | 26 |
| **OANTIGEN-PWY: O-antigen building blocks biosynthesis (E. coli)** | group | patient | grouppatient | NA | 0 | NA | NA | NA | 0.0317308748182228 | 0.1376771299747 | All logi | prevalence | 26 | 26 |
| **P124-PWY: Bifidobacterium shunt** | group | patient | grouppatient | NA | 0 | NA | NA | NA | 0.167612236693643 | 0.37873918868275 | All logi | prevalence | 26 | 26 |
| **P4-PWY: superpathway of L-lysine, L-threonine and L-methionine biosynthesis I** | group | patient | grouppatient | NA | 0 | NA | NA | NA | 0.444975090595048 | 0.650272858794624 | All logi | prevalence | 26 | 26 |
| **P41-PWY: pyruvate fermentation to acetate and (S)-lactate I** | group | patient | grouppatient | NA | 0 | NA | NA | NA | 0.956847142114468 | 0.973415923796103 | All logi | prevalence | 26 | 26 |
| **P42-PWY: incomplete reductive TCA cycle** | group | patient | grouppatient | NA | 0 | NA | NA | NA | 0.0325801166888491 | 0.1376771299747 | All logi | prevalence | 26 | 26 |
| **P441-PWY: superpathway of N-acetylneuraminate degradation** | group | patient | grouppatient | NA | 0 | NA | NA | NA | 0.569025557940628 | 0.764120034948844 | All logi | prevalence | 26 | 26 |
| **P461-PWY: hexitol fermentation to lactate, formate, ethanol and acetate** | group | patient | grouppatient | NA | 0 | NA | NA | NA | 0.0575031097192609 | 0.19205351836294 | All logi | prevalence | 26 | 26 |
| **PANTO-PWY: phosphopantothenate biosynthesis I** | group | patient | grouppatient | NA | 0 | NA | NA | NA | 0.116534408711014 | 0.300940506011959 | All logi | prevalence | 26 | 26 |
| **PANTOSYN-PWY: superpathway of coenzyme A biosynthesis I (bacteria)** | group | patient | grouppatient | NA | 0 | NA | NA | NA | 0.0394179269766175 | 0.161777392061913 | All logi | prevalence | 26 | 26 |
| **PENTOSE-P-PWY: pentose phosphate pathway** | group | patient | grouppatient | NA | 0 | NA | NA | NA | 0.139171126408718 | 0.333726680673967 | All logi | prevalence | 26 | 26 |
| **PEPTIDOGLYCANSYN-PWY: peptidoglycan biosynthesis I (meso-diaminopimelate containing)** | group | patient | grouppatient | NA | 0 | NA | NA | NA | 0.00254872201340461 | 0.035360858899724 | All logi | prevalence | 26 | 26 |
| **PHOSLIPSYN-PWY: superpathway of phospholipid biosynthesis I (bacteria)** | group | patient | grouppatient | NA | 0 | NA | NA | NA | 0.210931986425977 | 0.438663865576148 | All logi | prevalence | 26 | 26 |
| **POLYAMSYN-PWY: superpathway of polyamine biosynthesis I** | group | patient | grouppatient | NA | 0 | NA | NA | NA | 0.0607465629005262 | 0.192911382184103 | All logi | prevalence | 26 | 26 |
| **POLYISOPRENSYN-PWY: polyisoprenoid biosynthesis (E. coli)** | group | patient | grouppatient | NA | 0 | NA | NA | NA | 0.0202245267909783 | 0.113161042759045 | All logi | prevalence | 26 | 26 |
| **PWY-1042: glycolysis IV** | group | patient | grouppatient | NA | 0 | NA | NA | NA | 0.0153459218005365 | 0.0958791043121336 | All logi | prevalence | 26 | 26 |
| **PWY-1269: CMP-3-deoxy-D-manno-octulosonate biosynthesis** | group | patient | grouppatient | NA | 0 | NA | NA | NA | 0.576656395910136 | 0.765617248807243 | All logi | prevalence | 26 | 26 |
| **PWY-1861: formaldehyde assimilation II (assimilatory RuMP Cycle)** | group | patient | grouppatient | NA | 0 | NA | NA | NA | 0.236148199850129 | 0.470295143769324 | All logi | prevalence | 26 | 26 |
| **PWY-241: C4 photosynthetic carbon assimilation cycle, NADP-ME type** | group | patient | grouppatient | NA | 0 | NA | NA | NA | 0.790641920238239 | 0.898283823082983 | All logi | prevalence | 26 | 26 |
| **PWY-2941: L-lysine biosynthesis II** | group | patient | grouppatient | NA | 0 | NA | NA | NA | 0.0551581578796274 | 0.190620104436948 | All logi | prevalence | 26 | 26 |
| **PWY-2942: L-lysine biosynthesis III** | group | patient | grouppatient | NA | 0 | NA | NA | NA | 0.0314157601397214 | 0.1376771299747 | All logi | prevalence | 26 | 26 |
| **PWY-3001: superpathway of L-isoleucine biosynthesis I** | group | patient | grouppatient | NA | 0 | NA | NA | NA | 0.351533283348308 | 0.582470721166646 | All logi | prevalence | 26 | 26 |
| **PWY-3841: folate transformations II (plants)** | group | patient | grouppatient | NA | 0 | NA | NA | NA | 0.0210306027622696 | 0.114934689514729 | All logi | prevalence | 26 | 26 |
| **PWY-4041: γ-glutamyl cycle** | group | patient | grouppatient | NA | 0 | NA | NA | NA | 0.0464611862370106 | 0.176570604363487 | All logi | prevalence | 26 | 26 |
| **PWY-4984: urea cycle** | group | patient | grouppatient | NA | 0 | NA | NA | NA | 0.673695368348735 | 0.828892207130643 | All logi | prevalence | 26 | 26 |
| **PWY-5030: L-histidine degradation III** | group | patient | grouppatient | NA | 0 | NA | NA | NA | 0.681821886632165 | 0.834521579992494 | All logi | prevalence | 26 | 26 |
| **PWY-5097: L-lysine biosynthesis VI** | group | patient | grouppatient | NA | 0 | NA | NA | NA | 0.0506580532933352 | 0.180373371574754 | All logi | prevalence | 26 | 26 |
| **PWY-5100: pyruvate fermentation to acetate and lactate II** | group | patient | grouppatient | NA | 0 | NA | NA | NA | 0.956827944772579 | 0.973415923796103 | All logi | prevalence | 26 | 26 |
| **PWY-5103: L-isoleucine biosynthesis III** | group | patient | grouppatient | NA | 0 | NA | NA | NA | 0.0483650775962827 | 0.177952874143801 | All logi | prevalence | 26 | 26 |
| **PWY-5121: superpathway of geranylgeranyl diphosphate biosynthesis II (via MEP)** | group | patient | grouppatient | NA | 0 | NA | NA | NA | 0.0574216449570701 | 0.19205351836294 | All logi | prevalence | 26 | 26 |
| **PWY-5154: L-arginine biosynthesis III (via N-acetyl-L-citrulline)** | group | patient | grouppatient | NA | 0 | NA | NA | NA | 0.849552189390991 | 0.920021956252917 | All logi | prevalence | 26 | 26 |
| **PWY-5188: tetrapyrrole biosynthesis I (from glutamate)** | group | patient | grouppatient | NA | 0 | NA | NA | NA | 0.00573066506640418 | 0.0480966532358923 | All logi | prevalence | 26 | 26 |
| **PWY-5345: superpathway of L-methionine biosynthesis (by sulfhydrylation)** | group | patient | grouppatient | NA | 0 | NA | NA | NA | 0.9059100722294 | 0.937971660255113 | All logi | prevalence | 26 | 26 |

| | | | | | | | | | | | | | | |
|---|---|---|---|---|---|---|---|---|---|---|---|---|---|---|
| **PWY-5347: superpathway of L-methionine biosynthesis (transsulfuration)** | group | patient | grouppatient | NA | 0 | NA | NA | NA | 0.115874214849785 | 0.300940506011959 | All logi | prevalence | 26 | 26 |
| **PWY-5384: sucrose degradation IV (sucrose phosphorylase)** | group | patient | grouppatient | NA | 0 | NA | NA | NA | 0.200914513144202 | 0.421561701686495 | All logi | prevalence | 26 | 26 |
| **PWY-5484: glycolysis II (from fructose 6-phosphate)** | group | patient | grouppatient | NA | 0 | NA | NA | NA | 0.473581985313119 | 0.670432328605922 | All logi | prevalence | 26 | 26 |
| **PWY-5497: purine nucleobases degradation II (anaerobic)** | group | patient | grouppatient | NA | 0 | NA | NA | NA | 0.618071979156208 | 0.793698989626825 | All logi | prevalence | 26 | 26 |
| **PWY-5505: L-glutamate and L-glutamine biosynthesis** | group | patient | grouppatient | NA | 0 | NA | NA | NA | 0.312553132891737 | 0.552255535560588 | All logi | prevalence | 26 | 26 |
| **PWY-5659: GDP-mannose biosynthesis** | group | patient | grouppatient | NA | 0 | NA | NA | NA | 0.356918229140413 | 0.582470721166646 | All logi | prevalence | 26 | 26 |
| **PWY-5667: CDP-diacylglycerol biosynthesis I** | group | patient | grouppatient | NA | 0 | NA | NA | NA | 0.0594954905707814 | 0.192911382184103 | All logi | prevalence | 26 | 26 |
| **PWY-5676: acetyl-CoA fermentation to butanoate II** | group | patient | grouppatient | NA | 0 | NA | NA | NA | 0.132945362408033 | 0.324658529847235 | All logi | prevalence | 26 | 26 |
| **PWY-5686: UMP biosynthesis I** | group | patient | grouppatient | NA | 0 | NA | NA | NA | 0.00377358921222959 | 0.0362829955222034 | All logi | prevalence | 26 | 26 |
| **PWY-5695: inosine 5'-phosphate degradation** | group | patient | grouppatient | NA | 0 | NA | NA | NA | 0.0626874119698198 | 0.196420557505435 | All logi | prevalence | 26 | 26 |
| **PWY-5913: partial TCA cycle (obligate autotrophs)** | group | patient | grouppatient | NA | 0 | NA | NA | NA | 0.996554402095212 | 0.996554402095212 | All logi | prevalence | 26 | 26 |
| **PWY-5941: glycogen degradation II** | group | patient | grouppatient | NA | 0 | NA | NA | NA | 0.146128812781883 | 0.339856215863755 | All logi | prevalence | 26 | 26 |
| **PWY-5973: cis-vaccenate biosynthesis** | group | patient | grouppatient | NA | 0 | NA | NA | NA | 0.992190327573233 | 0.996430457178247 | All logi | prevalence | 26 | 26 |
| **PWY-5989: stearate biosynthesis II (bacteria and plants)** | group | patient | grouppatient | NA | 0 | NA | NA | NA | 0.224836935167403 | 0.451595553541365 | All logi | prevalence | 26 | 26 |
| **PWY-6121: 5-aminoimidazole ribonucleotide biosynthesis I** | group | patient | grouppatient | NA | 0 | NA | NA | NA | 0.0328081671429072 | 0.1376771299747 | All logi | prevalence | 26 | 26 |
| **PWY-6122: 5-aminoimidazole ribonucleotide biosynthesis II** | group | patient | grouppatient | NA | 0 | NA | NA | NA | 0.046584584981005 | 0.176570604363487 | All logi | prevalence | 26 | 26 |
| **PWY-6123: inosine-5'-phosphate biosynthesis I** | group | patient | grouppatient | NA | 0 | NA | NA | NA | 0.781623251426018 | 0.898283823082983 | All logi | prevalence | 26 | 26 |
| **PWY-6124: inosine-5'-phosphate biosynthesis II** | group | patient | grouppatient | NA | 0 | NA | NA | NA | 0.553175724032831 | 0.747105144527099 | All logi | prevalence | 26 | 26 |
| **PWY-6125: superpathway of guanosine nucleotides de novo biosynthesis II** | group | patient | grouppatient | NA | 0 | NA | NA | NA | 0.861583924941667 | 0.927823755660637 | All logi | prevalence | 26 | 26 |
| **PWY-6126: superpathway of adenosine nucleotides de novo biosynthesis II** | group | patient | grouppatient | NA | 0 | NA | NA | NA | 0.983165222402876 | 0.995878565796017 | All logi | prevalence | 26 | 26 |
| **PWY-6147: 6-hydroxymethyl-dihydropterin diphosphate biosynthesis I** | group | patient | grouppatient | NA | 0 | NA | NA | NA | 0.453727753100781 | 0.654147374102352 | All logi | prevalence | 26 | 26 |
| **PWY-6151: S-adenosyl-L-methionine salvage I** | group | patient | grouppatient | NA | 0 | NA | NA | NA | 0.00270849131997886 | 0.035360858899724 | All logi | prevalence | 26 | 26 |
| **PWY-6163: chorismate biosynthesis from 3-dehydroquinate** | group | patient | grouppatient | NA | 0 | NA | NA | NA | 0.00060179100897439 | 0.0202029838727118 | All logi | prevalence | 26 | 26 |
| **PWY-621: sucrose degradation III (sucrose invertase)** | group | patient | grouppatient | NA | 0 | NA | NA | NA | 0.543976580606623 | 0.743265095121777 | All logi | prevalence | 26 | 26 |
| **PWY-6270: isoprene biosynthesis I** | group | patient | grouppatient | NA | 0 | NA | NA | NA | 0.816016923544824 | 0.898283823082983 | All logi | prevalence | 26 | 26 |
| **PWY-6277: superpathway of 5-aminoimidazole ribonucleotide biosynthesis** | group | patient | grouppatient | NA | 0 | NA | NA | NA | 0.0463551170903638 | 0.176570604363487 | All logi | prevalence | 26 | 26 |
| **PWY-6282: palmitoleate biosynthesis I (from (5Z)-dodec-5-enoate)** | group | patient | grouppatient | NA | 0 | NA | NA | NA | 0.302814060274976 | 0.539100789125904 | All logi | prevalence | 26 | 26 |
| **PWY-6292: superpathway of L-cysteine biosynthesis (mammalian)** | group | patient | grouppatient | NA | 0 | NA | NA | NA | 0.887903121317938 | 0.937971660255113 | All logi | prevalence | 26 | 26 |
| **PWY-6305: superpathway of putrescine biosynthesis** | group | patient | grouppatient | NA | 0 | NA | NA | NA | 0.896049372885493 | 0.937971660255113 | All logi | prevalence | 26 | 26 |
| **PWY-6317: D-galactose degradation I (Leloir pathway)** | group | patient | grouppatient | NA | 0 | NA | NA | NA | 0.66335010483376 | 0.828892207130643 | All logi | prevalence | 26 | 26 |
| **PWY-6353: purine nucleotides degradation II (aerobic)** | group | patient | grouppatient | NA | 0 | NA | NA | NA | 0.147512059651502 | 0.339856215863755 | All logi | prevalence | 26 | 26 |
| **PWY-6385: peptidoglycan biosynthesis III (mycobacteria)** | group | patient | grouppatient | NA | 0 | NA | NA | NA | 0.00736961686233495 | 0.0577286654216238 | All logi | prevalence | 26 | 26 |
| **PWY-6386: UDP-N-acetylmuramoyl-pentapeptide biosynthesis II (lysine-containing)** | group | patient | grouppatient | NA | 0 | NA | NA | NA | 0.00324133589282471 | 0.0362720921339909 | All logi | prevalence | 26 | 26 |
| **PWY-6387: UDP-N-acetylmuramoyl-pentapeptide biosynthesis I (meso-diaminopimelate containing)** | group | patient | grouppatient | NA | 0 | NA | NA | NA | 0.00258453722588281 | 0.035360858899724 | All logi | prevalence | 26 | 26 |
| **PWY-6470: peptidoglycan biosynthesis V (β-lactam resistance)** | group | patient | grouppatient | NA | 0 | NA | NA | NA | 0.873130190968199 | 0.928441605780664 | All logi | prevalence | 26 | 26 |
| **PWY-6507: 4-deoxy-L-threo-hex-4-enopyranuronate degradation** | group | patient | grouppatient | NA | 0 | NA | NA | NA | 0.247826801743289 | 0.47349023097295 | All logi | prevalence | 26 | 26 |
| **PWY-6519: 8-amino-7-oxononanoate biosynthesis I** | group | patient | grouppatient | NA | 0 | NA | NA | NA | 0.371139775366626 | 0.596480867258721 | All logi | prevalence | 26 | 26 |
| **PWY-6527: stachyose degradation** | group | patient | grouppatient | NA | 0 | NA | NA | NA | 0.0674664704941699 | 0.205872735384621 | All logi | prevalence | 26 | 26 |
| **PWY-6549: L-glutamine biosynthesis III** | group | patient | grouppatient | NA | 0 | NA | NA | NA | 0.0125241043702746 | 0.0865636625592511 | All logi | prevalence | 26 | 26 |
| **PWY-6590: superpathway of Clostridium acetobutylicum acidogenic fermentation** | group | patient | grouppatient | NA | 0 | NA | NA | NA | 0.0321463711385879 | 0.1376771299747 | All logi | prevalence | 26 | 26 |
| **PWY-6606: guanosine nucleotides degradation II** | group | patient | grouppatient | NA | 0 | NA | NA | NA | 0.0915157682727008 | 0.256444928744021 | All logi | prevalence | 26 | 26 |
| **PWY-6608: guanosine nucleotides degradation III** | group | patient | grouppatient | NA | 0 | NA | NA | NA | 0.393800582560331 | 0.615253601110053 | All logi | prevalence | 26 | 26 |
| **PWY-6609: adenine and adenosine salvage III** | group | patient | grouppatient | NA | 0 | NA | NA | NA | 0.0163198475424908 | 0.0958791043121336 | All logi | prevalence | 26 | 26 |
| **PWY-6612: superpathway of tetrahydrofolate biosynthesis** | group | patient | grouppatient | NA | 0 | NA | NA | NA | 0.0161000364453403 | 0.0958791043121336 | All logi | prevalence | 26 | 26 |
| **PWY-6628: superpathway of L-phenylalanine biosynthesis** | group | patient | grouppatient | NA | 0 | NA | NA | NA | 0.614972305005135 | 0.793698989626825 | All logi | prevalence | 26 | 26 |
| **PWY-6629: superpathway of L-tryptophan biosynthesis** | group | patient | grouppatient | NA | 0 | NA | NA | NA | 0.00660847736350645 | 0.0535514544973798 | All logi | prevalence | 26 | 26 |
| **PWY-6630: superpathway of L-tyrosine biosynthesis** | group | patient | grouppatient | NA | 0 | NA | NA | NA | 0.0536375977019599 | 0.188131872536725 | All logi | prevalence | 26 | 26 |
| **PWY-6700: queuosine biosynthesis I (de novo)** | group | patient | grouppatient | NA | 0 | NA | NA | NA | 0.00343399775092545 | 0.0362829955222034 | All logi | prevalence | 26 | 26 |

| | | | | | | | | | | | | | | |
|---|---|---|---|---|---|---|---|---|---|---|---|---|---|---|
| **PWY-6703: preQ0 biosynthesis** | group | patient | grouppatient | NA | 0 | NA | NA | NA | 0.598224673155841 | 0.783998669568306 | All logi | prevalence | 26 | 26 |
| **PWY-6731: starch degradation III** | group | patient | grouppatient | NA | 0 | NA | NA | NA | 0.802743123481785 | 0.898283823082983 | All logi | prevalence | 26 | 26 |
| **PWY-6823: molybdopterin biosynthesis** | group | patient | grouppatient | NA | 0 | NA | NA | NA | 0.0912630689572964 | 0.256444928744021 | All logi | prevalence | 26 | 26 |
| **PWY-6859: all-trans-farnesol biosynthesis** | group | patient | grouppatient | NA | 0 | NA | NA | NA | 0.298677835621593 | 0.535796117336446 | All logi | prevalence | 26 | 26 |
| **PWY-6892: thiazole component of thiamine diphosphate biosynthesis I** | group | patient | grouppatient | NA | 0 | NA | NA | NA | 0.445506086238019 | 0.650272858794624 | All logi | prevalence | 26 | 26 |
| **PWY-6895: superpathway of thiamine diphosphate biosynthesis II** | group | patient | grouppatient | NA | 0 | NA | NA | NA | 0.448978763388504 | 0.651296354298139 | All logi | prevalence | 26 | 26 |
| **PWY-6897: thiamine diphosphate salvage II** | group | patient | grouppatient | NA | 0 | NA | NA | NA | 0.419957044288525 | 0.637566226374194 | All logi | prevalence | 26 | 26 |
| **PWY-6901: superpathway of glucose and xylose degradation** | group | patient | grouppatient | NA | 0 | NA | NA | NA | 0.297065174972515 | 0.535796117336446 | All logi | prevalence | 26 | 26 |
| **PWY-6902: chitin degradation II (Vibrio)** | group | patient | grouppatient | NA | 0 | NA | NA | NA | 0.608623918907542 | 0.790202325653439 | All logi | prevalence | 26 | 26 |
| **PWY-6936: seleno-amino acid biosynthesis (plants)** | group | patient | grouppatient | NA | 0 | NA | NA | NA | 0.333762278098648 | 0.572511936884543 | All logi | prevalence | 26 | 26 |
| **PWY-6969: TCA cycle V (2-oxoglutarate synthase)** | group | patient | grouppatient | NA | 0 | NA | NA | NA | 0.00319226844644505 | 0.0362720921339909 | All logi | prevalence | 26 | 26 |
| **PWY-702: L-methionine biosynthesis II** | group | patient | grouppatient | NA | 0 | NA | NA | NA | 0.284415165436337 | 0.526811382008272 | All logi | prevalence | 26 | 26 |
| **PWY-7111: pyruvate fermentation to isobutanol (engineered)** | group | patient | grouppatient | NA | 0 | NA | NA | NA | 0.395333164968587 | 0.615253601110053 | All logi | prevalence | 26 | 26 |
| **PWY-7115: C4 photosynthetic carbon assimilation cycle, NAD-ME type** | group | patient | grouppatient | NA | 0 | NA | NA | NA | 0.355716039034448 | 0.582470721166646 | All logi | prevalence | 26 | 26 |
| **PWY-7117: C4 photosynthetic carbon assimilation cycle, PEPCK type** | group | patient | grouppatient | NA | 0 | NA | NA | NA | 0.194817538765151 | 0.420019464310188 | All logi | prevalence | 26 | 26 |
| **PWY-7197: pyrimidine deoxyribonucleotide phosphorylation** | group | patient | grouppatient | NA | 0 | NA | NA | NA | 0.919102149750727 | 0.943183428783497 | All logi | prevalence | 26 | 26 |
| **PWY-7198: pyrimidine deoxyribonucleotides de novo biosynthesis IV** | group | patient | grouppatient | NA | 0 | NA | NA | NA | 0.807534879206931 | 0.898283823082983 | All logi | prevalence | 26 | 26 |
| **PWY-7199: pyrimidine deoxyribonucleosides salvage** | group | patient | grouppatient | NA | 0 | NA | NA | NA | 0.0580246800160373 | 0.19205351836294 | All logi | prevalence | 26 | 26 |
| **PWY-7208: superpathway of pyrimidine nucleobases salvage** | group | patient | grouppatient | NA | 0 | NA | NA | NA | 0.686380915392172 | 0.835748782990468 | All logi | prevalence | 26 | 26 |
| **PWY-7220: adenosine deoxyribonucleotides de novo biosynthesis II** | group | patient | grouppatient | NA | 0 | NA | NA | NA | 0.654172511141769 | 0.82708026078924 | All logi | prevalence | 26 | 26 |
| **PWY-7221: guanosine ribonucleotides de novo biosynthesis** | group | patient | grouppatient | NA | 0 | NA | NA | NA | 0.0268865389508887 | 0.131632013613726 | All logi | prevalence | 26 | 26 |
| **PWY-7222: guanosine deoxyribonucleotides de novo biosynthesis II** | group | patient | grouppatient | NA | 0 | NA | NA | NA | 0.654625227688505 | 0.82708026078924 | All logi | prevalence | 26 | 26 |
| **PWY-7228: superpathway of guanosine nucleotides de novo biosynthesis I** | group | patient | grouppatient | NA | 0 | NA | NA | NA | 0.894736422091422 | 0.937971660255113 | All logi | prevalence | 26 | 26 |
| **PWY-7229: superpathway of adenosine nucleotides de novo biosynthesis I** | group | patient | grouppatient | NA | 0 | NA | NA | NA | 0.819679562027325 | 0.898283823082983 | All logi | prevalence | 26 | 26 |
| **PWY-7234: inosine-5'-phosphate biosynthesis III** | group | patient | grouppatient | NA | 0 | NA | NA | NA | 0.403433667760898 | 0.623729683709283 | All logi | prevalence | 26 | 26 |
| **PWY-7237: myo-, chiro- and scyllo-inositol degradation** | group | patient | grouppatient | NA | 0 | NA | NA | NA | 0.321164588822173 | 0.562417635258954 | All logi | prevalence | 26 | 26 |
| **PWY-7238: sucrose biosynthesis II** | group | patient | grouppatient | NA | 0 | NA | NA | NA | 0.000130359272577207 | 0.00734183923282705 | All logi | prevalence | 26 | 26 |
| **PWY-724: superpathway of L-lysine, L-threonine and L-methionine biosynthesis II** | group | patient | grouppatient | NA | 0 | NA | NA | NA | 0.0138078243384523 | 0.0927096777010368 | All logi | prevalence | 26 | 26 |
| **PWY-7242: D-fructuronate degradation** | group | patient | grouppatient | NA | 0 | NA | NA | NA | 0.736623387725378 | 0.857259089956876 | All logi | prevalence | 26 | 26 |
| **PWY-7282: 4-amino-2-methyl-5-diphosphomethylpyrimidine biosynthesis II** | group | patient | grouppatient | NA | 0 | NA | NA | NA | 0.133250732921563 | 0.324658529847235 | All logi | prevalence | 26 | 26 |
| **PWY-7323: superpathway of GDP-mannose-derived O-antigen building blocks biosynthesis** | group | patient | grouppatient | NA | 0 | NA | NA | NA | 0.864807138085979 | 0.927823755660637 | All logi | prevalence | 26 | 26 |
| **PWY-7328: superpathway of UDP-glucose-derived O-antigen building blocks biosynthesis** | group | patient | grouppatient | NA | 0 | NA | NA | NA | 0.73546701350296 | 0.857259089956876 | All logi | prevalence | 26 | 26 |
| **PWY-7345: superpathway of anaerobic sucrose degradation** | group | patient | grouppatient | NA | 0 | NA | NA | NA | 0.219204991400463 | 0.447940634600947 | All logi | prevalence | 26 | 26 |
| **PWY-7356: thiamine diphosphate salvage IV (yeast)** | group | patient | grouppatient | NA | 0 | NA | NA | NA | 0.720426226135731 | 0.857259089956876 | All logi | prevalence | 26 | 26 |
| **PWY-7357: thiamine phosphate formation from pyrithiamine and oxythiamine (yeast)** | group | patient | grouppatient | NA | 0 | NA | NA | NA | 0.465941728119073 | 0.663613976412014 | All logi | prevalence | 26 | 26 |
| **PWY-7383: anaerobic energy metabolism (invertebrates, cytosol)** | group | patient | grouppatient | NA | 0 | NA | NA | NA | 0.145515302717343 | 0.339856215863755 | All logi | prevalence | 26 | 26 |
| **PWY-7392: taxadiene biosynthesis (engineered)** | group | patient | grouppatient | NA | 0 | NA | NA | NA | 0.362445053348153 | 0.587410948529766 | All logi | prevalence | 26 | 26 |
| **PWY-7400: L-arginine biosynthesis IV (archaebacteria)** | group | patient | grouppatient | NA | 0 | NA | NA | NA | 0.199315708486388 | 0.421561701686495 | All logi | prevalence | 26 | 26 |
| **PWY-7456: β-(1,4)-mannan degradation** | group | patient | grouppatient | NA | 0 | NA | NA | NA | 0.115893671813868 | 0.300940506011959 | All logi | prevalence | 26 | 26 |
| **PWY-7560: methylerythritol phosphate pathway II** | group | patient | grouppatient | NA | 0 | NA | NA | NA | 0.662297189187871 | 0.828892207130643 | All logi | prevalence | 26 | 26 |
| **PWY-7663: gondoate biosynthesis (anaerobic)** | group | patient | grouppatient | NA | 0 | NA | NA | NA | 0.736878026260804 | 0.857259089956876 | All logi | prevalence | 26 | 26 |
| **PWY-7664: oleate biosynthesis IV (anaerobic)** | group | patient | grouppatient | NA | 0 | NA | NA | NA | 0.353377360654344 | 0.582470721166646 | All logi | prevalence | 26 | 26 |
| **PWY-7761: NAD salvage pathway II (PNC IV cycle)** | group | patient | grouppatient | NA | 0 | NA | NA | NA | 0.509125626742101 | 0.712169775502344 | All logi | prevalence | 26 | 26 |
| **PWY-7790: UMP biosynthesis II** | group | patient | grouppatient | NA | 0 | NA | NA | NA | 0.00385989314065993 | 0.0362829955222034 | All logi | prevalence | 26 | 26 |
| **PWY-7791: UMP biosynthesis III** | group | patient | grouppatient | NA | 0 | NA | NA | NA | 0.00359264854958341 | 0.0362829955222034 | All logi | prevalence | 26 | 26 |
| **PWY-7851: coenzyme A biosynthesis II (eukaryotic)** | group | patient | grouppatient | NA | 0 | NA | NA | NA | 0.00493931980602857 | 0.042990376089508 | All logi | prevalence | 26 | 26 |
| **PWY-7953: UDP-N-acetylmuramoyl-pentapeptide biosynthesis III (meso-diaminopimelate containing** | group | patient | grouppatient | NA | 0 | NA | NA | NA | 0.0012745107934875 | 0.0252289488287085 | All logi | prevalence | 26 | 26 |
| **PWY-7977: L-methionine biosynthesis IV** | group | patient | grouppatient | NA | 0 | NA | NA | NA | 0.125398150838717 | 0.313495377096792 | All logi | prevalence | 26 | 26 |

| | | | | | | | | | | | | | | |
|---|---|---|---|---|---|---|---|---|---|---|---|---|---|---|
| **PWY-8004: Entner-Doudoroff pathway I** | group | patient | grouppatient | NA | 0 | NA | NA | NA | 0.37792567639198 | 0.600084688865643 | All logi | prevalence | 26 | 26 |
| **PWY-8073: lipid IVA biosynthesis (P. putida)** | group | patient | grouppatient | NA | 0 | NA | NA | NA | 0.696983546521302 | 0.840038360049195 | All logi | prevalence | 26 | 26 |
| **PWY-8178: pentose phosphate pathway (non-oxidative branch) II** | group | patient | grouppatient | NA | 0 | NA | NA | NA | 0.224670667011089 | 0.451595553541365 | All logi | prevalence | 26 | 26 |
| **PWY-8187: L-arginine degradation XIII (reductive Stickland reaction)** | group | patient | grouppatient | NA | 0 | NA | NA | NA | 0.193980604934943 | 0.420019464310188 | All logi | prevalence | 26 | 26 |
| **PWY-841: superpathway of purine nucleotides de novo biosynthesis I** | group | patient | grouppatient | NA | 0 | NA | NA | NA | 0.821834136012091 | 0.898283823082983 | All logi | prevalence | 26 | 26 |
| **PWY-I9: L-cysteine biosynthesis VI (from L-methionine)** | group | patient | grouppatient | NA | 0 | NA | NA | NA | 0.600509619243809 | 0.783998669568306 | All logi | prevalence | 26 | 26 |
| **PWY0-1061: superpathway of L-alanine biosynthesis** | group | patient | grouppatient | NA | 0 | NA | NA | NA | 0.262236721762759 | 0.496980883985874 | All logi | prevalence | 26 | 26 |
| **PWY0-1261: anhydromuropeptides recycling I** | group | patient | grouppatient | NA | 0 | NA | NA | NA | 0.442753339702805 | 0.650272858794624 | All logi | prevalence | 26 | 26 |
| **PWY0-1296: purine ribonucleosides degradation** | group | patient | grouppatient | NA | 0 | NA | NA | NA | 0.072612716910922 | 0.213299855925833 | All logi | prevalence | 26 | 26 |
| **PWY0-1297: superpathway of purine deoxyribonucleosides degradation** | group | patient | grouppatient | NA | 0 | NA | NA | NA | 0.00150300120681668 | 0.0252289488287085 | All logi | prevalence | 26 | 26 |
| **PWY0-1298: superpathway of pyrimidine deoxyribonucleosides degradation** | group | patient | grouppatient | NA | 0 | NA | NA | NA | 0.0916654213382883 | 0.256444928744021 | All logi | prevalence | 26 | 26 |
| **PWY0-1319: CDP-diacylglycerol biosynthesis II** | group | patient | grouppatient | NA | 0 | NA | NA | NA | 0.0600841597774608 | 0.192911382184103 | All logi | prevalence | 26 | 26 |
| **PWY0-1477: ethanolamine utilization** | group | patient | grouppatient | NA | 0 | NA | NA | NA | 0.216385601523096 | 0.446058038227435 | All logi | prevalence | 26 | 26 |
| **PWY0-1479: tRNA processing** | group | patient | grouppatient | NA | 0 | NA | NA | NA | 0.991512237494714 | 0.996430457178247 | All logi | prevalence | 26 | 26 |
| **PWY0-1586: peptidoglycan maturation (meso-diaminopimelate containing)** | group | patient | grouppatient | NA | 0 | NA | NA | NA | 0.553148770026826 | 0.747105144527099 | All logi | prevalence | 26 | 26 |
| **PWY0-162: superpathway of pyrimidine ribonucleotides de novo biosynthesis** | group | patient | grouppatient | NA | 0 | NA | NA | NA | 0.733823074127899 | 0.857259089956876 | All logi | prevalence | 26 | 26 |
| **PWY0-781: aspartate superpathway** | group | patient | grouppatient | NA | 0 | NA | NA | NA | 0.456594124055243 | 0.654265970445013 | All logi | prevalence | 26 | 26 |
| **PWY0-845: superpathway of pyridoxal 5'-phosphate biosynthesis and salvage** | group | patient | grouppatient | NA | 0 | NA | NA | NA | 0.282049499270006 | 0.526811382008272 | All logi | prevalence | 26 | 26 |
| **PWY0-862: (5Z)-dodecenoate biosynthesis I** | group | patient | grouppatient | NA | 0 | NA | NA | NA | 0.350800541623694 | 0.582470721166646 | All logi | prevalence | 26 | 26 |
| **PWY3O-4107: NAD salvage pathway V (PNC V cycle)** | group | patient | grouppatient | NA | 0 | NA | NA | NA | 0.119856574074604 | 0.303593636907051 | All logi | prevalence | 26 | 26 |
| **PWY4FS-7: phosphatidylglycerol biosynthesis I (plastidic)** | group | patient | grouppatient | NA | 0 | NA | NA | NA | 0.0220419847255684 | 0.11588664948301 | All logi | prevalence | 26 | 26 |
| **PWY4FS-8: phosphatidylglycerol biosynthesis II (non-plastidic)** | group | patient | grouppatient | NA | 0 | NA | NA | NA | 0.0221910605392998 | 0.11588664948301 | All logi | prevalence | 26 | 26 |
| **PWY66-399: gluconeogenesis III** | group | patient | grouppatient | NA | 0 | NA | NA | NA | 0.288289326865537 | 0.526811382008272 | All logi | prevalence | 26 | 26 |
| **PWY66-409: superpathway of purine nucleotide salvage** | group | patient | grouppatient | NA | 0 | NA | NA | NA | 0.00988779992311084 | 0.0735495657425225 | All logi | prevalence | 26 | 26 |
| **PWY66-429: fatty acid biosynthesis initiation (mitochondria)** | group | patient | grouppatient | NA | 0 | NA | NA | NA | 0.14593752492409 | 0.339856215863755 | All logi | prevalence | 26 | 26 |
| **PYRIDNUCSAL-PWY: NAD salvage pathway I (PNC VI cycle)** | group | patient | grouppatient | NA | 0 | NA | NA | NA | 0.0700170050994608 | 0.208278432890801 | All logi | prevalence | 26 | 26 |
| **PYRIDNUCSYN-PWY: NAD de novo biosynthesis I (from aspartate)** | group | patient | grouppatient | NA | 0 | NA | NA | NA | 0.0118678620949859 | 0.0845135634036876 | All logi | prevalence | 26 | 26 |
| **PYRIDOXSYN-PWY: pyridoxal 5'-phosphate biosynthesis I** | group | patient | grouppatient | NA | 0 | NA | NA | NA | 0.373117819093753 | 0.596480867258721 | All logi | prevalence | 26 | 26 |
| **RHAMCAT-PWY: L-rhamnose degradation I** | group | patient | grouppatient | NA | 0 | NA | NA | NA | 0.087037025217356 | 0.252514826247885 | All logi | prevalence | 26 | 26 |
| **RIBOSYN2-PWY: flavin biosynthesis I (bacteria and plants)** | group | patient | grouppatient | NA | 0 | NA | NA | NA | 0.101904978617467 | 0.278461278780288 | All logi | prevalence | 26 | 26 |
| **SALVADEHYPOX-PWY: adenosine nucleotides degradation II** | group | patient | grouppatient | NA | 0 | NA | NA | NA | 0.0506191190283587 | 0.180373371574754 | All logi | prevalence | 26 | 26 |
| **SER-GLYSYN-PWY: superpathway of L-serine and glycine biosynthesis I** | group | patient | grouppatient | NA | 0 | NA | NA | NA | 0.151269021225661 | 0.345128349398354 | All logi | prevalence | 26 | 26 |
| **SULFATE-CYS-PWY: superpathway of sulfate assimilation and cysteine biosynthesis** | group | patient | grouppatient | NA | 0 | NA | NA | NA | 0.905521442879331 | 0.937971660255113 | All logi | prevalence | 26 | 26 |
| **THISYN-PWY: superpathway of thiamine diphosphate biosynthesis I** | group | patient | grouppatient | NA | 0 | NA | NA | NA | 0.351282766619543 | 0.582470721166646 | All logi | prevalence | 26 | 26 |
| **THISYNARA-PWY: superpathway of thiamine diphosphate biosynthesis III (eukaryotes)** | group | patient | grouppatient | NA | 0 | NA | NA | NA | 0.0399280371897488 | 0.161777392061913 | All logi | prevalence | 26 | 26 |
| **THRESYN-PWY: superpathway of L-threonine biosynthesis** | group | patient | grouppatient | NA | 0 | NA | NA | NA | 0.868600962746128 | 0.927823755660637 | All logi | prevalence | 26 | 26 |
| **TRNA-CHARGING-PWY: tRNA charging** | group | patient | grouppatient | NA | 0 | NA | NA | NA | 0.0014445790484896 | 0.0252289488287085 | All logi | prevalence | 26 | 26 |
| **TRPSYN-PWY: L-tryptophan biosynthesis** | group | patient | grouppatient | NA | 0 | NA | NA | NA | 0.0306444478216408 | 0.1376771299747 | All logi | prevalence | 26 | 26 |
| **UDPNAGSYN-PWY: UDP-N-acetyl-D-glucosamine biosynthesis I** | group | patient | grouppatient | NA | 0 | NA | NA | NA | 0.728684144826606 | 0.857259089956876 | All logi | prevalence | 26 | 26 |
| **VALSYN-PWY: L-valine biosynthesis** | group | patient | grouppatient | NA | 0 | NA | NA | NA | 0.00118761348409757 | 0.0252289488287085 | All logi | prevalence | 26 | 26 |

**Supplementary Table 3: Differential Pathway Abundance Analysis (MaAsLin3), Candidate pathways during abatacept tratment, factorial model**

| feature | metadata | value | name | coef | null_hypothesis | stderr | pval_individual | qval_individual | pval_joint | qval_joint | error | model | N | N_not_zero |
|---|---|---|---|---|---|---|---|---|---|---|---|---|---|---|
| **FOLSYN-PWY: superpathway of tetrahydrofolate biosynthesis and salvage** | date_closest | 12 | date_closest12 | -1.07511607736697 | 0.0373621103475272 | 0.422623451436789 | 0.0209368600135256 | 0.399524912747588 | 0.0209368600135256 | 0.399524912747588 | NA | abundance | 27 | 27 |
| **PWY-6612: superpathway of tetrahydrofolate biosynthesis** | date_closest | 12 | date_closest12 | -1.11464311534615 | 0.0373621103475272 | 0.439965206937408 | 0.021307995346538 | 0.399524912747588 | 0.021307995346538 | 0.399524912747588 | NA | abundance | 27 | 27 |
| **PWY0-1297: superpathway of purine deoxyribonucleosides degradation** | date_closest | 12 | date_closest12 | -1.05956202453685 | 0.0373621103475272 | 0.382670127220344 | 0.0143277910835775 | 0.399524912747588 | 0.0143277910835775 | 0.399524912747588 | NA | abundance | 27 | 27 |
| **PWY0-1297: superpathway of purine deoxyribonucleosides degradation** | date_closest | 6 | date_closest6 | -0.823925570852375 | 0.0934413552518843 | 0.338442505689119 | 0.0184421024377775 | 0.399524912747588 | 0.0184421024377775 | 0.399524912747588 | NA | abundance | 27 | 27 |
| **FOLSYN-PWY: superpathway of tetrahydrofolate biosynthesis and salvage** | date_closest | 6 | date_closest6 | -0.785862698688146 | 0.0934413552518843 | 0.374277378615709 | 0.0352400322508079 | 0.43716804399067 | 0.0352400322508079 | 0.43716804399067 | NA | abundance | 27 | 27 |
| **PWY-6612: superpathway of tetrahydrofolate biosynthesis** | date_closest | 6 | date_closest6 | -0.820235768238008 | 0.0934413552518843 | 0.389612624330749 | 0.0352444498565663 | 0.43716804399067 | 0.0352444498565663 | 0.43716804399067 | NA | abundance | 27 | 27 |
| **PWY66-409: superpathway of purine nucleotide salvage** | date_closest | 12 | date_closest12 | -0.614343175672491 | 0.0373621103475272 | 0.270629099242257 | 0.0408023507724625 | 0.43716804399067 | 0.0408023507724625 | 0.43716804399067 | NA | abundance | 27 | 27 |
| **PWY66-409: superpathway of purine nucleotide salvage** | date_closest | 6 | date_closest6 | -0.415908064539792 | 0.0934413552518843 | 0.239219843300707 | 0.0619191636582739 | 0.580492159296318 | 0.0619191636582739 | 0.580492159296318 | NA | abundance | 27 | 27 |
| **FERMENTATION-PWY: mixed acid fermentation** | date_closest | 3 | date_closest3 | 0.254107865543844 | -0.0572374156074166 | 0.167924759721138 | 0.110147450907228 | 0.827750526003359 | 0.110147450907228 | 0.827750526003359 | NA | abundance | 27 | 27 |
| **FERMENTATION-PWY: mixed acid fermentation** | date_closest | 6 | date_closest6 | 0.383018834494565 | 0.0934413552518843 | 0.158983059282369 | 0.12721210360468 5 | 0.827750526003359 | 0.127212103604685 | 0.827750526003359 | NA | abundance | 27 | 27 |
| **PWY-5676: acetyl-CoA fermentation to butanoate II** | date_closest | 12 | date_closest12 | -0.517373750946077 | 0.0373621103475272 | 0.33582410297509 | 0.132440084160537 | 0.827750526003359 | 0.132440084160537 | 0.827750526003359 | NA | abundance | 27 | 27 |
| **PWY0-1297: superpathway of purine deoxyribonucleosides degradation** | date_closest | 3 | date_closest3 | -0.644107263728101 | -0.0572374156074166 | 0.357249708730896 | 0.125598132020704 | 0.827750526003359 | 0.125598132020704 | 0.827750526003359 | NA | abundance | 27 | 27 |
| **ARGSYN-PWY: L-arginine biosynthesis I (via L-ornithine)** | date_closest | 3 | date_closest3 | -0.00170292939525374 | -0.0572374156074166 | 0.0829252681550574 | 0.631809731230481 | 0.88012155533212 | 0.631809731230481 | 0.88012155533212 | NA | abundance | 27 | 27 |
| **ARGSYN-PWY: L-arginine biosynthesis I (via L-ornithine)** | date_closest | 6 | date_closest6 | 0.122042258944398 | 0.0934413552518843 | 0.0788082687414934 | 0.803587636869462 | 0.88012155533212 | 0.803587636869462 | 0.88012155533212 | NA | abundance | 27 | 27 |
| **ARGSYNBSUB-PWY: L-arginine biosynthesis II (acetyl cycle)** | date_closest | 3 | date_closest3 | 0.00020748912369371 | -0.0572374156074166 | 0.0878866552915053 | 0.631694800110912 | 0.88012155533212 | 0.631694800110912 | 0.88012155533212 | NA | abundance | 27 | 27 |
| **ARGSYNBSUB-PWY: L-arginine biosynthesis II (acetyl cycle)** | date_closest | 6 | date_closest6 | 0.137827961785778 | 0.0934413552518843 | 0.0834412090249699 | 0.709610507628353 | 0.88012155533212 | 0.709610507628353 | 0.88012155533212 | NA | abundance | 27 | 27 |
| **ARO-PWY: chorismate biosynthesis I** | date_closest | 12 | date_closest12 | 0.0739982412354402 | 0.0373621103475272 | 0.102469856314459 | 0.80971183090555 | 0.88012155533212 | 0.80971183090555 | 0.88012155533212 | NA | abundance | 27 | 27 |
| **ARO-PWY: chorismate biosynthesis I** | date_closest | 3 | date_closest3 | 0.0373824615525472 | -0.0572374156074166 | 0.0960534433852208 | 0.437443341501433 | 0.88012155533212 | 0.437443341501433 | 0.88012155533212 | NA | abundance | 27 | 27 |
| **ARO-PWY: chorismate biosynthesis I** | date_closest | 6 | date_closest6 | 0.181059826758345 | 0.0934413552518843 | 0.0912558376804639 | 0.489983246668 | 0.88012155533212 | 0.489983246668 | 0.88012155533212 | NA | abundance | 27 | 27 |
| **BRANCHED-CHAIN-AA-SYN-PWY: superpathway of branched chain amino acid biosynthesis** | date_closest | 12 | date_closest12 | -0.00818504314152224 | 0.0373621103475272 | 0.0776392297837773 | 0.738499517463882 | 0.88012155533212 | 0.738499517463882 | 0.88012155533212 | NA | abundance | 27 | 27 |
| **BRANCHED-CHAIN-AA-SYN-PWY: superpathway of branched chain amino acid biosynthesis** | date_closest | 3 | date_closest3 | -0.0012958994606633 | -0.0572374156074166 | 0.0733621764251692 | 0.608992949829751 | 0.88012155533212 | 0.608992949829751 | 0.88012155533212 | NA | abundance | 27 | 27 |
| **BRANCHED-CHAIN-AA-SYN-PWY: superpathway of branched chain amino acid biosynthesis** | date_closest | 6 | date_closest6 | 0.0648404515593706 | 0.0934413552518843 | 0.0701476576851196 | 0.79313964052178 | 0.88012155533212 | 0.79313964052178 | 0.88012155533212 | NA | abundance | 27 | 27 |
| **CENTFERM-PWY: pyruvate fermentation to butanoate** | date_closest | 12 | date_closest12 | 0.363064526502543 | 0.0373621103475272 | 0.413016898295874 | 0.454077813805704 | 0.88012155533212 | 0.454077813805704 | 0.88012155533212 | NA | abundance | 27 | 27 |
| **CENTFERM-PWY: pyruvate fermentation to butanoate** | date_closest | 3 | date_closest3 | 0.454219589800323 | -0.0572374156074166 | 0.386172082102694 | 0.209718691324272 | 0.88012155533212 | 0.209718691324272 | 0.88012155533212 | NA | abundance | 27 | 27 |
| **CENTFERM-PWY: pyruvate fermentation to butanoate** | date_closest | 6 | date_closest6 | 0.191695184300292 | 0.0934413552518843 | 0.366219617997731 | 0.793935037558904 | 0.88012155533212 | 0.793935037558904 | 0.88012155533212 | NA | abundance | 27 | 27 |
| **COA-PWY-1: superpathway of coenzyme A biosynthesis III (mammals)** | date_closest | 12 | date_closest12 | -0.0183468545389849 | 0.0373621103475272 | 0.0956920547716307 | 0.703287078347031 | 0.88012155533212 | 0.703287078347031 | 0.88012155533212 | NA | abundance | 27 | 27 |
| **COA-PWY-1: superpathway of coenzyme A biosynthesis III (mammals)** | date_closest | 3 | date_closest3 | -0.125848350799938 | -0.0572374156074166 | 0.0904204926323238 | 0.558179927064229 | 0.88012155533212 | 0.558179927064229 | 0.88012155533212 | NA | abundance | 27 | 27 |
| **COA-PWY-1: superpathway of coenzyme A biosynthesis III (mammals)** | date_closest | 6 | date_closest6 | 0.0229216558809668 | 0.0934413552518843 | 0.0864585277314105 | 0.556946650124804 | 0.88012155533212 | 0.556946650124804 | 0.88012155533212 | NA | abundance | 27 | 27 |
| **COA-PWY: coenzyme A biosynthesis I (prokaryotic)** | date_closest | 12 | date_closest12 | -0.0117047413805483 | 0.0373621103475272 | 0.0966597333638194 | 0.738870117995778 | 0.88012155533212 | 0.738870117995778 | 0.88012155533212 | NA | abundance | 27 | 27 |
| **COA-PWY: coenzyme A biosynthesis I (prokaryotic)** | date_closest | 3 | date_closest3 | -0.13564083966247 | -0.0572374156074166 | 0.091334862955171 | 0.506287312118883 | 0.88012155533212 | 0.506287312118883 | 0.88012155533212 | NA | abundance | 27 | 27 |
| **COA-PWY: coenzyme A biosynthesis I (prokaryotic)** | date_closest | 6 | date_closest6 | 0.057206760900983 | 0.0934413552518843 | 0.0873328329869252 | 0.763332878455233 | 0.88012155533212 | 0.763332878455233 | 0.88012155533212 | NA | abundance | 27 | 27 |
| **COMPLETE-ARO-PWY: superpathway of aromatic amino acid biosynthesis** | date_closest | 12 | date_closest12 | 0.0816365262292778 | 0.0373621103475272 | 0.0982056642130953 | 0.767861216945989 | 0.88012155533212 | 0.767861216945989 | 0.88012155533212 | NA | abundance | 27 | 27 |
| **COMPLETE-ARO-PWY: superpathway of aromatic amino acid biosynthesis** | date_closest | 3 | date_closest3 | 0.0327336841932747 | -0.0572374156074166 | 0.091992999827555 | 0.446328069906658 | 0.88012155533212 | 0.446328069906658 | 0.88012155533212 | NA | abundance | 27 | 27 |
| **COMPLETE-ARO-PWY: superpathway of aromatic amino acid biosynthesis** | date_closest | 6 | date_closest6 | 0.178382936044478 | 0.0934413552518843 | 0.0873543816484682 | 0.492807332637846 | 0.88012155533212 | 0.492807332637846 | 0.88012155533212 | NA | abundance | 27 | 27 |
| **FOLSYN-PWY: superpathway of tetrahydrofolate biosynthesis and salvage** | date_closest | 3 | date_closest3 | -0.448447781484001 | -0.0572374156074166 | 0.394865551367823 | 0.33861037445777 | 0.88012155533212 | 0.33861037445777 | 0.88012155533212 | NA | abundance | 27 | 27 |
| **HISTSYN-PWY: L-histidine biosynthesis** | date_closest | 12 | date_closest12 | 0.157647893970105 | 0.0373621103475272 | 0.0896538840621834 | 0.419532892608786 | 0.88012155533212 | 0.419532892608786 | 0.88012155533212 | NA | abundance | 27 | 27 |
| **HISTSYN-PWY: L-histidine biosynthesis** | date_closest | 3 | date_closest3 | 0.103033846140812 | -0.0572374156074166 | 0.0844494577684767 | 0.177811413193298 | 0.88012155533212 | 0.177811413193298 | 0.88012155533212 | NA | abundance | 27 | 27 |
| **HISTSYN-PWY: L-histidine biosynthesis** | date_closest | 6 | date_closest6 | 0.210973742629202 | 0.0934413552518843 | 0.0805348781890463 | 0.333240767013498 | 0.88012155533212 | 0.333240767013498 | 0.88012155533212 | NA | abundance | 27 | 27 |
| **NAGLIPASYN-PWY: lipid IVA biosynthesis (E. coli)** | date_closest | 12 | date_closest12 | 0.801332124231087 | 0.0373621103475272 | 0.637555507737512 | 0.252259650286331 | 0.88012155533212 | 0.252259650286331 | 0.88012155533212 | NA | abundance | 27 | 27 |
| **NAGLIPASYN-PWY: lipid IVA biosynthesis (E. coli)** | date_closest | 3 | date_closest3 | -0.611851512999976 | -0.0572374156074166 | 0.596462309446682 | 0.364224269214988 | 0.88012155533212 | 0.364224269214988 | 0.88012155533212 | NA | abundance | 27 | 27 |
| **NAGLIPASYN-PWY: lipid IVA biosynthesis (E. coli)** | date_closest | 6 | date_closest6 | -0.135695967595182 | 0.0934413552518843 | 0.565872862808472 | 0.69059939183748 | 0.88012155533212 | 0.69059939183748 | 0.88012155533212 | NA | abundance | 27 | 27 |
| **PEPTIDOGLYCANSYN-PWY: peptidoglycan biosynthesis I (meso-diaminopimelate containing** | date_closest | 12 | date_closest12 | -0.0830720504311478 | 0.0373621103475272 | 0.101321647288622 | 0.430659685245749 | 0.88012155533212 | 0.430659685245749 | 0.88012155533212 | NA | abundance | 27 | 27 |
| **PEPTIDOGLYCANSYN-PWY: peptidoglycan biosynthesis I (meso-diaminopimelate containing** | date_closest | 6 | date_closest6 | 0.0229126663584192 | 0.0934413552518843 | 0.0915449090606486 | 0.57035882921506 | 0.88012155533212 | 0.57035882921506 | 0.88012155533212 | NA | abundance | 27 | 27 |
| **PWY-5676: acetyl-CoA fermentation to butanoate II** | date_closest | 3 | date_closest3 | 0.0696334205934607 | -0.0572374156074166 | 0.316298771310109 | 0.694081142803406 | 0.88012155533212 | 0.694081142803406 | 0.88012155533212 | NA | abundance | 27 | 27 |
| **PWY-5676: acetyl-CoA fermentation to butanoate II** | date_closest | 6 | date_closest6 | -0.130364538969629 | 0.0934413552518843 | 0.301613111840782 | 0.47823458190988 | 0.88012155533212 | 0.47823458190988 | 0.88012155533212 | NA | abundance | 27 | 27 |

| | | | | | | | | | | | | | | |
|---|---|---|---|---|---|---|---|---|---|---|---|---|---|---|
| **PWY-5686: UMP biosynthesis I** | date_closest | 12 | date_closest12 | -0.100888072393597 | 0.0373621103475272 | 0.0978799209148726 | 0.361807036290137 | 0.88012155533212 | 0.361807036290137 | 0.88012155533212 | NA | abundance | 27 | 27 |
| **PWY-5686: UMP biosynthesis I** | date_closest | 3 | date_closest3 | -0.115752324999125 | -0.0572374156074166 | 0.0924878318169367 | 0.619933117984289 | 0.88012155533212 | 0.619933117984289 | 0.88012155533212 | NA | abundance | 27 | 27 |
| **PWY-5686: UMP biosynthesis I** | date_closest | 6 | date_closest6 | 0.00152258289626418 | 0.0934413552518843 | 0.0884352820823284 | 0.46652278526528 | 0.88012155533212 | 0.46652278526528 | 0.88012155533212 | NA | abundance | 27 | 27 |
| **PWY-6163: chorismate biosynthesis from 3-dehydroquinate** | date_closest | 12 | date_closest12 | 0.0972465959219921 | 0.0373621103475272 | 0.108925655415935 | 0.704067656843595 | 0.88012155533212 | 0.704067656843595 | 0.88012155533212 | NA | abundance | 27 | 27 |
| **PWY-6163: chorismate biosynthesis from 3-dehydroquinate** | date_closest | 3 | date_closest3 | 0.043039004998909 | -0.0572374156074166 | 0.102127253433021 | 0.429263364960297 | 0.88012155533212 | 0.429263364960297 | 0.88012155533212 | NA | abundance | 27 | 27 |
| **PWY-6163: chorismate biosynthesis from 3-dehydroquinate** | date_closest | 6 | date_closest6 | 0.195044189382807 | 0.0934413552518843 | 0.0970418702699061 | 0.43913474043453 | 0.88012155533212 | 0.43913474043453 | 0.88012155533212 | NA | abundance | 27 | 27 |
| **PWY-6590: superpathway of Clostridium acetobutylicum acidogenic fermentation** | date_closest | 12 | date_closest12 | 0.348330996397843 | 0.0373621103475272 | 0.404990533341072 | 0.465965025844077 | 0.88012155533212 | 0.465965025844077 | 0.88012155533212 | NA | abundance | 27 | 27 |
| **PWY-6590: superpathway of Clostridium acetobutylicum acidogenic fermentation** | date_closest | 3 | date_closest3 | 0.446459943026654 | -0.0572374156074166 | 0.378708168774755 | 0.207822592265658 | 0.88012155533212 | 0.207822592265658 | 0.88012155533212 | NA | abundance | 27 | 27 |
| **PWY-6590: superpathway of Clostridium acetobutylicum acidogenic fermentation** | date_closest | 6 | date_closest6 | 0.191098642432899 | 0.0934413552518843 | 0.359168039115779 | 0.791778050155704 | 0.88012155533212 | 0.791778050155704 | 0.88012155533212 | NA | abundance | 27 | 27 |
| **PWY-6612: superpathway of tetrahydrofolate biosynthesis** | date_closest | 3 | date_closest3 | -0.475970403422939 | -0.0572374156074166 | 0.411053991397781 | 0.325138502519887 | 0.88012155533212 | 0.325138502519887 | 0.88012155533212 | NA | abundance | 27 | 27 |
| **PWY-6629: superpathway of L-tryptophan biosynthesis** | date_closest | 12 | date_closest12 | 0.104447025148364 | 0.0373621103475272 | 0.0850691063522094 | 0.640639114478816 | 0.88012155533212 | 0.640639114478816 | 0.88012155533212 | NA | abundance | 27 | 27 |
| **PWY-6629: superpathway of L-tryptophan biosynthesis** | date_closest | 3 | date_closest3 | 0.0457470288654679 | -0.0572374156074166 | 0.0795235411154728 | 0.353029969424313 | 0.88012155533212 | 0.353029969424313 | 0.88012155533212 | NA | abundance | 27 | 27 |
| **PWY-6629: superpathway of L-tryptophan biosynthesis** | date_closest | 6 | date_closest6 | 0.165012974628322 | 0.0934413552518843 | 0.075404128303343 | 0.53411941060099 | 0.88012155533212 | 0.53411941060099 | 0.88012155533212 | NA | abundance | 27 | 27 |
| **PWY-6969: TCA cycle V (2-oxoglutarate synthase)** | date_closest | 12 | date_closest12 | 0.281730442281884 | 0.0373621103475272 | 0.253416889037547 | 0.386649821697898 | 0.88012155533212 | 0.386649821697898 | 0.88012155533212 | NA | abundance | 27 | 27 |
| **PWY-6969: TCA cycle V (2-oxoglutarate synthase)** | date_closest | 6 | date_closest6 | 0.232666434228853 | 0.0934413552518843 | 0.224186221305276 | 0.560913189645744 | 0.88012155533212 | 0.560913189645744 | 0.88012155533212 | NA | abundance | 27 | 27 |
| **PWY-7851: coenzyme A biosynthesis II (eukaryotic)** | date_closest | 12 | date_closest12 | -0.0136591832348093 | 0.0373621103475272 | 0.0970153738295966 | 0.728748739507872 | 0.88012155533212 | 0.728748739507872 | 0.88012155533212 | NA | abundance | 27 | 27 |
| **PWY-7851: coenzyme A biosynthesis II (eukaryotic)** | date_closest | 3 | date_closest3 | -0.132297967284641 | -0.0572374156074166 | 0.0916709116082416 | 0.526535102657615 | 0.88012155533212 | 0.526535102657615 | 0.88012155533212 | NA | abundance | 27 | 27 |
| **PWY-7851: coenzyme A biosynthesis II (eukaryotic)** | date_closest | 6 | date_closest6 | 0.0258481005359582 | 0.0934413552518843 | 0.0876541569583478 | 0.578303454340907 | 0.88012155533212 | 0.578303454340907 | 0.88012155533212 | NA | abundance | 27 | 27 |
| **PWY66-409: superpathway of purine nucleotide salvage** | date_closest | 3 | date_closest3 | -0.406736940099216 | -0.0572374156074166 | 0.25256761676337 | 0.199669171229678 | 0.88012155533212 | 0.199669171229678 | 0.88012155533212 | NA | abundance | 27 | 27 |
| **TRPSYN-PWY: L-tryptophan biosynthesis** | date_closest | 12 | date_closest12 | 0.129782266239232 | 0.0373621103475272 | 0.0788890732361541 | 0.512979944473576 | 0.88012155533212 | 0.512979944473576 | 0.88012155533212 | NA | abundance | 27 | 27 |
| **TRPSYN-PWY: L-tryptophan biosynthesis** | date_closest | 3 | date_closest3 | 0.0566430183038661 | -0.0572374156074166 | 0.0736629239872013 | 0.292943261258251 | 0.88012155533212 | 0.292943261258251 | 0.88012155533212 | NA | abundance | 27 | 27 |
| **TRPSYN-PWY: L-tryptophan biosynthesis** | date_closest | 6 | date_closest6 | 0.156143500399692 | 0.0934413552518843 | 0.0697939223215085 | 0.573727898506427 | 0.88012155533212 | 0.573727898506427 | 0.88012155533212 | NA | abundance | 27 | 27 |
| **VALSYN-PWY: L-valine biosynthesis** | date_closest | 12 | date_closest12 | -0.0577741471971323 | 0.0373621103475272 | 0.074559023147995 | 0.487837601298499 | 0.88012155533212 | 0.487837601298499 | 0.88012155533212 | NA | abundance | 27 | 27 |
| **VALSYN-PWY: L-valine biosynthesis** | date_closest | 6 | date_closest6 | 0.0509375749429972 | 0.0934413552518843 | 0.066822653128692 | 0.693179231500069 | 0.88012155533212 | 0.693179231500069 | 0.88012155533212 | NA | abundance | 27 | 27 |
| **VALSYN-PWY: L-valine biosynthesis** | date_closest | 3 | date_closest3 | -0.0375198869271014 | -0.0572374156074166 | 0.0701422072912102 | 0.844436497987452 | 0.904753390700841 | 0.844436497987452 | 0.904753390700841 | NA | abundance | 27 | 27 |
| **PEPTIDOGLYCANSYN-PWY: peptidoglycan biosynthesis I (meso-diaminopimelate containing** | date_closest | 3 | date_closest3 | -0.0769549442877317 | -0.0572374156074166 | 0.0957399575546772 | 0.867229345968977 | 0.916087337291173 | 0.867229345968977 | 0.916087337291173 | NA | abundance | 27 | 27 |
| **ARGSYNBSUB-PWY: L-arginine biosynthesis II (acetyl cycle)** | date_closest | 12 | date_closest12 | 0.0496993902323654 | 0.0373621103475272 | 0.0938436388734846 | 0.932663015012955 | 0.954339524415889 | 0.932663015012955 | 0.954339524415889 | NA | abundance | 27 | 27 |
| **FERMENTATION-PWY: mixed acid fermentation** | date_closest | 12 | date_closest12 | 0.0528023014177823 | 0.0373621103475272 | 0.180053816194564 | 0.941614997423677 | 0.954339524415889 | 0.941614997423677 | 0.954339524415889 | NA | abundance | 27 | 27 |
| **PWY-6969: TCA cycle V (2-oxoglutarate synthase)** | date_closest | 3 | date_closest3 | -0.0770002627674954 | -0.0572374156074166 | 0.236619830539017 | 0.935173624657668 | 0.954339524415889 | 0.935173624657668 | 0.954339524415889 | NA | abundance | 27 | 27 |
| **ARGSYN-PWY: L-arginine biosynthesis I (via L-ornithine)** | date_closest | 12 | date_closest12 | 0.0373621103475272 | 0.0373621103475272 | 0.088426909382851 | 1 | 1 | 1 | 1 | NA | abundance | 27 | 27 |
| **ARGSYN-PWY: L-arginine biosynthesis I (via L-ornithine)** | date_closest | 12 | date_closest12 | NA | 0 | NA | NA | NA | 1 | 1 | All logi | prevalence | 27 | 27 |
| **ARGSYN-PWY: L-arginine biosynthesis I (via L-ornithine)** | date_closest | 3 | date_closest3 | NA | 0 | NA | NA | NA | 0.631809731230481 | 0.88012155533212 | All logi | prevalence | 27 | 27 |
| **ARGSYN-PWY: L-arginine biosynthesis I (via L-ornithine)** | date_closest | 6 | date_closest6 | NA | 0 | NA | NA | NA | 0.803587636869462 | 0.88012155533212 | All logi | prevalence | 27 | 27 |
| **ARGSYNBSUB-PWY: L-arginine biosynthesis II (acetyl cycle)** | date_closest | 12 | date_closest12 | NA | 0 | NA | NA | NA | 0.932663015012955 | 0.954339524415889 | All logi | prevalence | 27 | 27 |
| **ARGSYNBSUB-PWY: L-arginine biosynthesis II (acetyl cycle)** | date_closest | 3 | date_closest3 | NA | 0 | NA | NA | NA | 0.631694800110912 | 0.88012155533212 | All logi | prevalence | 27 | 27 |
| **ARGSYNBSUB-PWY: L-arginine biosynthesis II (acetyl cycle)** | date_closest | 6 | date_closest6 | NA | 0 | NA | NA | NA | 0.709610507628353 | 0.88012155533212 | All logi | prevalence | 27 | 27 |
| **ARO-PWY: chorismate biosynthesis I** | date_closest | 12 | date_closest12 | NA | 0 | NA | NA | NA | 0.80971183090555 | 0.88012155533212 | All logi | prevalence | 27 | 27 |
| **ARO-PWY: chorismate biosynthesis I** | date_closest | 3 | date_closest3 | NA | 0 | NA | NA | NA | 0.437443341501433 | 0.88012155533212 | All logi | prevalence | 27 | 27 |
| **ARO-PWY: chorismate biosynthesis I** | date_closest | 6 | date_closest6 | NA | 0 | NA | NA | NA | 0.489983246668 | 0.88012155533212 | All logi | prevalence | 27 | 27 |
| **BRANCHED-CHAIN-AA-SYN-PWY: superpathway of branched chain amino acid biosynthesis** | date_closest | 12 | date_closest12 | NA | 0 | NA | NA | NA | 0.738499517463882 | 0.88012155533212 | All logi | prevalence | 27 | 27 |
| **BRANCHED-CHAIN-AA-SYN-PWY: superpathway of branched chain amino acid biosynthesis** | date_closest | 3 | date_closest3 | NA | 0 | NA | NA | NA | 0.608992949829751 | 0.88012155533212 | All logi | prevalence | 27 | 27 |
| **BRANCHED-CHAIN-AA-SYN-PWY: superpathway of branched chain amino acid biosynthesis** | date_closest | 6 | date_closest6 | NA | 0 | NA | NA | NA | 0.79313964052178 | 0.88012155533212 | All logi | prevalence | 27 | 27 |
| **CENTFERM-PWY: pyruvate fermentation to butanoate** | date_closest | 12 | date_closest12 | NA | 0 | NA | NA | NA | 0.454077813805704 | 0.88012155533212 | All logi | prevalence | 27 | 27 |
| **CENTFERM-PWY: pyruvate fermentation to butanoate** | date_closest | 3 | date_closest3 | NA | 0 | NA | NA | NA | 0.209718691324272 | 0.88012155533212 | All logi | prevalence | 27 | 27 |
| **CENTFERM-PWY: pyruvate fermentation to butanoate** | date_closest | 6 | date_closest6 | NA | 0 | NA | NA | NA | 0.793935037558904 | 0.88012155533212 | All logi | prevalence | 27 | 27 |
| **COA-PWY-1: superpathway of coenzyme A biosynthesis III (mammals)** | date_closest | 12 | date_closest12 | NA | 0 | NA | NA | NA | 0.703287078347031 | 0.88012155533212 | All logi | prevalence | 27 | 27 |
| **COA-PWY-1: superpathway of coenzyme A biosynthesis III (mammals)** | date_closest | 3 | date_closest3 | NA | 0 | NA | NA | NA | 0.558179927064229 | 0.88012155533212 | All logi | prevalence | 27 | 27 |

| | | | | | | | | | | | | | | |
|---|---|---|---|---|---|---|---|---|---|---|---|---|---|---|
| **COA-PWY-1: superpathway of coenzyme A biosynthesis III (mammals)** | date_closest | 6 | date_closest6 | NA | 0 | NA | NA | NA | 0.556946650124804 | 0.88012155533212 | All logi | prevalence | 27 | 27 |
| **COA-PWY: coenzyme A biosynthesis I (prokaryotic)** | date_closest | 12 | date_closest12 | NA | 0 | NA | NA | NA | 0.738870117995778 | 0.88012155533212 | All logi | prevalence | 27 | 27 |
| **COA-PWY: coenzyme A biosynthesis I (prokaryotic)** | date_closest | 3 | date_closest3 | NA | 0 | NA | NA | NA | 0.506287312118883 | 0.88012155533212 | All logi | prevalence | 27 | 27 |
| **COA-PWY: coenzyme A biosynthesis I (prokaryotic)** | date_closest | 6 | date_closest6 | NA | 0 | NA | NA | NA | 0.763332878455233 | 0.88012155533212 | All logi | prevalence | 27 | 27 |
| **COMPLETE-ARO-PWY: superpathway of aromatic amino acid biosynthesis** | date_closest | 12 | date_closest12 | NA | 0 | NA | NA | NA | 0.767861216945989 | 0.88012155533212 | All logi | prevalence | 27 | 27 |
| **COMPLETE-ARO-PWY: superpathway of aromatic amino acid biosynthesis** | date_closest | 3 | date_closest3 | NA | 0 | NA | NA | NA | 0.446328069906658 | 0.88012155533212 | All logi | prevalence | 27 | 27 |
| **COMPLETE-ARO-PWY: superpathway of aromatic amino acid biosynthesis** | date_closest | 6 | date_closest6 | NA | 0 | NA | NA | NA | 0.492807332637846 | 0.88012155533212 | All logi | prevalence | 27 | 27 |
| **FERMENTATION-PWY: mixed acid fermentation** | date_closest | 12 | date_closest12 | NA | 0 | NA | NA | NA | 0.941614997423677 | 0.954339524415889 | All logi | prevalence | 27 | 27 |
| **FERMENTATION-PWY: mixed acid fermentation** | date_closest | 3 | date_closest3 | NA | 0 | NA | NA | NA | 0.110147450907228 | 0.827750526003359 | All logi | prevalence | 27 | 27 |
| **FERMENTATION-PWY: mixed acid fermentation** | date_closest | 6 | date_closest6 | NA | 0 | NA | NA | NA | 0.127212103604685 | 0.827750526003359 | All logi | prevalence | 27 | 27 |
| **FOLSYN-PWY: superpathway of tetrahydrofolate biosynthesis and salvage** | date_closest | 12 | date_closest12 | NA | 0 | NA | NA | NA | 0.0209368600135256 | 0.399524912747588 | All logi | prevalence | 27 | 27 |
| **FOLSYN-PWY: superpathway of tetrahydrofolate biosynthesis and salvage** | date_closest | 3 | date_closest3 | NA | 0 | NA | NA | NA | 0.33861037445777 | 0.88012155533212 | All logi | prevalence | 27 | 27 |
| **FOLSYN-PWY: superpathway of tetrahydrofolate biosynthesis and salvage** | date_closest | 6 | date_closest6 | NA | 0 | NA | NA | NA | 0.0352400322508079 | 0.43716804399067 | All logi | prevalence | 27 | 27 |
| **HISTSYN-PWY: L-histidine biosynthesis** | date_closest | 12 | date_closest12 | NA | 0 | NA | NA | NA | 0.419532892608786 | 0.88012155533212 | All logi | prevalence | 27 | 27 |
| **HISTSYN-PWY: L-histidine biosynthesis** | date_closest | 3 | date_closest3 | NA | 0 | NA | NA | NA | 0.177811413193298 | 0.88012155533212 | All logi | prevalence | 27 | 27 |
| **HISTSYN-PWY: L-histidine biosynthesis** | date_closest | 6 | date_closest6 | NA | 0 | NA | NA | NA | 0.333240767013498 | 0.88012155533212 | All logi | prevalence | 27 | 27 |
| **NAGLIPASYN-PWY: lipid IVA biosynthesis (E. coli)** | date_closest | 12 | date_closest12 | NA | 0 | NA | NA | NA | 0.252259650286331 | 0.88012155533212 | All logi | prevalence | 27 | 27 |
| **NAGLIPASYN-PWY: lipid IVA biosynthesis (E. coli)** | date_closest | 3 | date_closest3 | NA | 0 | NA | NA | NA | 0.364224269214988 | 0.88012155533212 | All logi | prevalence | 27 | 27 |
| **NAGLIPASYN-PWY: lipid IVA biosynthesis (E. coli)** | date_closest | 6 | date_closest6 | NA | 0 | NA | NA | NA | 0.69059939183748 | 0.88012155533212 | All logi | prevalence | 27 | 27 |
| **PEPTIDOGLYCANSYN-PWY: peptidoglycan biosynthesis I (meso-diaminopimelate containing** | date_closest | 12 | date_closest12 | NA | 0 | NA | NA | NA | 0.430659685245749 | 0.88012155533212 | All logi | prevalence | 27 | 27 |
| **PEPTIDOGLYCANSYN-PWY: peptidoglycan biosynthesis I (meso-diaminopimelate containing** | date_closest | 3 | date_closest3 | NA | 0 | NA | NA | NA | 0.867229345968977 | 0.916087337291173 | All logi | prevalence | 27 | 27 |
| **PEPTIDOGLYCANSYN-PWY: peptidoglycan biosynthesis I (meso-diaminopimelate containing** | date_closest | 6 | date_closest6 | NA | 0 | NA | NA | NA | 0.57035882921506 | 0.88012155533212 | All logi | prevalence | 27 | 27 |
| **PWY-5676: acetyl-CoA fermentation to butanoate II** | date_closest | 12 | date_closest12 | NA | 0 | NA | NA | NA | 0.132440084160537 | 0.827750526003359 | All logi | prevalence | 27 | 27 |
| **PWY-5676: acetyl-CoA fermentation to butanoate II** | date_closest | 3 | date_closest3 | NA | 0 | NA | NA | NA | 0.694081142803406 | 0.88012155533212 | All logi | prevalence | 27 | 27 |
| **PWY-5676: acetyl-CoA fermentation to butanoate II** | date_closest | 6 | date_closest6 | NA | 0 | NA | NA | NA | 0.47823458190988 | 0.88012155533212 | All logi | prevalence | 27 | 27 |
| **PWY-5686: UMP biosynthesis I** | date_closest | 12 | date_closest12 | NA | 0 | NA | NA | NA | 0.361807036290137 | 0.88012155533212 | All logi | prevalence | 27 | 27 |
| **PWY-5686: UMP biosynthesis I** | date_closest | 3 | date_closest3 | NA | 0 | NA | NA | NA | 0.619933117984289 | 0.88012155533212 | All logi | prevalence | 27 | 27 |
| **PWY-5686: UMP biosynthesis I** | date_closest | 6 | date_closest6 | NA | 0 | NA | NA | NA | 0.46652278526528 | 0.88012155533212 | All logi | prevalence | 27 | 27 |
| **PWY-6163: chorismate biosynthesis from 3-dehydroquinate** | date_closest | 12 | date_closest12 | NA | 0 | NA | NA | NA | 0.704067656843595 | 0.88012155533212 | All logi | prevalence | 27 | 27 |
| **PWY-6163: chorismate biosynthesis from 3-dehydroquinate** | date_closest | 3 | date_closest3 | NA | 0 | NA | NA | NA | 0.429263364960297 | 0.88012155533212 | All logi | prevalence | 27 | 27 |
| **PWY-6163: chorismate biosynthesis from 3-dehydroquinate** | date_closest | 6 | date_closest6 | NA | 0 | NA | NA | NA | 0.43913474043453 | 0.88012155533212 | All logi | prevalence | 27 | 27 |
| **PWY-6590: superpathway of Clostridium acetobutylicum acidogenic fermentation** | date_closest | 12 | date_closest12 | NA | 0 | NA | NA | NA | 0.465965025844077 | 0.88012155533212 | All logi | prevalence | 27 | 27 |
| **PWY-6590: superpathway of Clostridium acetobutylicum acidogenic fermentation** | date_closest | 3 | date_closest3 | NA | 0 | NA | NA | NA | 0.207822592265658 | 0.88012155533212 | All logi | prevalence | 27 | 27 |
| **PWY-6590: superpathway of Clostridium acetobutylicum acidogenic fermentation** | date_closest | 6 | date_closest6 | NA | 0 | NA | NA | NA | 0.791778050155704 | 0.88012155533212 | All logi | prevalence | 27 | 27 |
| **PWY-6612: superpathway of tetrahydrofolate biosynthesis** | date_closest | 12 | date_closest12 | NA | 0 | NA | NA | NA | 0.021307995346538 | 0.399524912747588 | All logi | prevalence | 27 | 27 |
| **PWY-6612: superpathway of tetrahydrofolate biosynthesis** | date_closest | 3 | date_closest3 | NA | 0 | NA | NA | NA | 0.325138502519887 | 0.88012155533212 | All logi | prevalence | 27 | 27 |
| **PWY-6612: superpathway of tetrahydrofolate biosynthesis** | date_closest | 6 | date_closest6 | NA | 0 | NA | NA | NA | 0.0352444498565663 | 0.43716804399067 | All logi | prevalence | 27 | 27 |
| **PWY-6629: superpathway of L-tryptophan biosynthesis** | date_closest | 12 | date_closest12 | NA | 0 | NA | NA | NA | 0.640639114478816 | 0.88012155533212 | All logi | prevalence | 27 | 27 |
| **PWY-6629: superpathway of L-tryptophan biosynthesis** | date_closest | 3 | date_closest3 | NA | 0 | NA | NA | NA | 0.353029969424313 | 0.88012155533212 | All logi | prevalence | 27 | 27 |
| **PWY-6629: superpathway of L-tryptophan biosynthesis** | date_closest | 6 | date_closest6 | NA | 0 | NA | NA | NA | 0.53411941060099 | 0.88012155533212 | All logi | prevalence | 27 | 27 |
| **PWY-6969: TCA cycle V (2-oxoglutarate synthase)** | date_closest | 12 | date_closest12 | NA | 0 | NA | NA | NA | 0.386649821697898 | 0.88012155533212 | All logi | prevalence | 27 | 27 |
| **PWY-6969: TCA cycle V (2-oxoglutarate synthase)** | date_closest | 3 | date_closest3 | NA | 0 | NA | NA | NA | 0.935173624657668 | 0.954339524415889 | All logi | prevalence | 27 | 27 |
| **PWY-6969: TCA cycle V (2-oxoglutarate synthase)** | date_closest | 6 | date_closest6 | NA | 0 | NA | NA | NA | 0.560913189645744 | 0.88012155533212 | All logi | prevalence | 27 | 27 |
| **PWY-7851: coenzyme A biosynthesis II (eukaryotic)** | date_closest | 12 | date_closest12 | NA | 0 | NA | NA | NA | 0.728748739507872 | 0.88012155533212 | All logi | prevalence | 27 | 27 |
| **PWY-7851: coenzyme A biosynthesis II (eukaryotic)** | date_closest | 3 | date_closest3 | NA | 0 | NA | NA | NA | 0.526535102657615 | 0.88012155533212 | All logi | prevalence | 27 | 27 |
| **PWY-7851: coenzyme A biosynthesis II (eukaryotic)** | date_closest | 6 | date_closest6 | NA | 0 | NA | NA | NA | 0.578303454340907 | 0.88012155533212 | All logi | prevalence | 27 | 27 |
| **PWY0-1297: superpathway of purine deoxyribonucleosides degradation** | date_closest | 12 | date_closest12 | NA | 0 | NA | NA | NA | 0.0143277910835775 | 0.399524912747588 | All logi | prevalence | 27 | 27 |

| | | | | | | | | | | | | | | |
|---|---|---|---|---|---|---|---|---|---|---|---|---|---|---|
| **PWY0-1297: superpathway of purine deoxyribonucleosides degradation** | date_closest | 3 | date_closest3 | NA | 0 | NA | NA | NA | 0.125598132020704 | 0.827750526003359 | All logi | prevalence | 27 | 27 |
| **PWY0-1297: superpathway of purine deoxyribonucleosides degradation** | date_closest | 6 | date_closest6 | NA | 0 | NA | NA | NA | 0.0184421024377775 | 0.399524912747588 | All logi | prevalence | 27 | 27 |
| **PWY66-409: superpathway of purine nucleotide salvage** | date_closest | 12 | date_closest12 | NA | 0 | NA | NA | NA | 0.0408023507724625 | 0.43716804399067 | All logi | prevalence | 27 | 27 |
| **PWY66-409: superpathway of purine nucleotide salvage** | date_closest | 3 | date_closest3 | NA | 0 | NA | NA | NA | 0.199669171229678 | 0.88012155533212 | All logi | prevalence | 27 | 27 |
| **PWY66-409: superpathway of purine nucleotide salvage** | date_closest | 6 | date_closest6 | NA | 0 | NA | NA | NA | 0.0619191636582739 | 0.580492159296318 | All logi | prevalence | 27 | 27 |
| **TRPSYN-PWY: L-tryptophan biosynthesis** | date_closest | 12 | date_closest12 | NA | 0 | NA | NA | NA | 0.512979944473576 | 0.88012155533212 | All logi | prevalence | 27 | 27 |
| **TRPSYN-PWY: L-tryptophan biosynthesis** | date_closest | 3 | date_closest3 | NA | 0 | NA | NA | NA | 0.292943261258251 | 0.88012155533212 | All logi | prevalence | 27 | 27 |
| **TRPSYN-PWY: L-tryptophan biosynthesis** | date_closest | 6 | date_closest6 | NA | 0 | NA | NA | NA | 0.573727898506427 | 0.88012155533212 | All logi | prevalence | 27 | 27 |
| **VALSYN-PWY: L-valine biosynthesis** | date_closest | 12 | date_closest12 | NA | 0 | NA | NA | NA | 0.487837601298499 | 0.88012155533212 | All logi | prevalence | 27 | 27 |
| **VALSYN-PWY: L-valine biosynthesis** | date_closest | 3 | date_closest3 | NA | 0 | NA | NA | NA | 0.844436497987452 | 0.904753390700841 | All logi | prevalence | 27 | 27 |
| **VALSYN-PWY: L-valine biosynthesis** | date_closest | 6 | date_closest6 | NA | 0 | NA | NA | NA | 0.693179231500069 | 0.88012155533212 | All logi | prevalence | 27 | 27 |

#### Supplementary Table 4: Differential Pathway Abundance Analysis (MaAsLin3), Candidate pathways during abatacept tratment, linear model

| feature | metadata | value | name | coef | null_hypothesis | stderr | pval_individual | qval_individual | pval_joint | qval_joint | error | model | N | N_not_zero |
|---|---|---|---|---|---|---|---|---|---|---|---|---|---|---|
| **FOLSYN-PWY: superpathway of tetrahydrofolate biosynthesis and salvage** | diff_to_baseine | diff_to_baseine | diff_to_baseine | -0.077465422169652 | 0.00486976258617576 | 0.0270323183556582 | 0.00844490981944305 | 0.074786057918652 | 0.00844490981944305 | 0.074786057918652 | NA | abundance | 27 | 27 |
| **PWY-6612: superpathway of tetrahydrofolate biosynthesis** | diff_to_baseine | diff_to_baseine | diff_to_baseine | -0.080391283924664 | 0.00486976258617576 | 0.0281574696527548 | 0.00862244897214534 | 0.074786057918652 | 0.00862244897214534 | 0.074786057918652 | NA | abundance | 27 | 27 |
| **PWY0-1297: superpathway of purine deoxyribonucleosides degradation** | diff_to_baseine | diff_to_baseine | diff_to_baseine | -0.0730904398464333 | 0.00486976258617576 | 0.0255667259454162 | 0.00897432695023823 | 0.074786057918652 | 0.00897432695023823 | 0.074786057918652 | NA | abundance | 27 | 27 |
| **PWY66-409: superpathway of purine nucleotide salvage** | diff_to_baseine | diff_to_baseine | diff_to_baseine | -0.0412816479506746 | 0.00486976258617576 | 0.017796032553341 | 0.0284115110850622 | 0.177571944281638 | 0.0284115110850622 | 0.177571944281638 | NA | abundance | 27 | 27 |
| **PWY-5676: acetyl-CoA fermentation to butanoate II** | diff_to_baseine | diff_to_baseine | diff_to_baseine | -0.029394005894114 | 0.00486976258617576 | 0.0217028329836536 | 0.148775579356197 | 0.743877896780986 | 0.148775579356197 | 0.743877896780986 | NA | abundance | 27 | 27 |
| **ARO-PWY: chorismate biosynthesis I** | diff_to_baseine | diff_to_baseine | diff_to_baseine | 0.00796169999531402 | 0.00486976258617576 | 0.00695424135177877 | 0.765677864069327 | 0.925861580484465 | 0.765677864069327 | 0.925861580484465 | NA | abundance | 27 | 27 |
| **BRANCHED-CHAIN-AA-SYN-PWY: superpathway of branched chain amino acid biosynthesis** | diff_to_baseine | diff_to_baseine | diff_to_baseine | 0.000604689250094451 | 0.00486976258617576 | 0.00498139029592231 | 0.64325454100129 | 0.925861580484465 | 0.64325454100129 | 0.925861580484465 | NA | abundance | 27 | 27 |
| **CENTFERM-PWY: pyruvate fermentation to butanoate** | diff_to_baseine | diff_to_baseine | diff_to_baseine | 0.0188601678009767 | 0.00486976258617576 | 0.0273435165848699 | 0.623670817236797 | 0.925861580484465 | 0.623670817236797 | 0.925861580484465 | NA | abundance | 27 | 27 |
| **COA-PWY-1: superpathway of coenzyme A biosynthesis III (mammals)** | diff_to_baseine | diff_to_baseine | diff_to_baseine | 0.00210921118708447 | 0.00486976258617576 | 0.0063264033507329 | 0.779451238021495 | 0.925861580484465 | 0.779451238021495 | 0.925861580484465 | NA | abundance | 27 | 27 |
| **COMPLETE-ARO-PWY: superpathway of aromatic amino acid biosynthesis** | diff_to_baseine | diff_to_baseine | diff_to_baseine | 0.00829588598040132 | 0.00486976258617576 | 0.00670399296798357 | 0.737889157635735 | 0.925861580484465 | 0.737889157635735 | 0.925861580484465 | NA | abundance | 27 | 27 |
| **FERMENTATION-PWY: mixed acid fermentation** | diff_to_baseine | diff_to_baseine | diff_to_baseine | 0.0128622218361892 | 0.00486976258617576 | 0.0134151567133063 | 0.606439414085784 | 0.925861580484465 | 0.606439414085784 | 0.925861580484465 | NA | abundance | 27 | 27 |
| **HISTSYN-PWY: L-histidine biosynthesis** | diff_to_baseine | diff_to_baseine | diff_to_baseine | 0.0124461337174578 | 0.00486976258617576 | 0.00595481277103076 | 0.449317639423979 | 0.925861580484465 | 0.449317639423979 | 0.925861580484465 | NA | abundance | 27 | 27 |
| **NAGLIPASYN-PWY: lipid IVA biosynthesis (E. coli)** | diff_to_baseine | diff_to_baseine | diff_to_baseine | 0.0461578231726279 | 0.00486976258617576 | 0.0434092520659379 | 0.357496237264372 | 0.925861580484465 | 0.357496237264372 | 0.925861580484465 | NA | abundance | 27 | 27 |
| **PEPTIDOGLYCANSYN-PWY: peptidoglycan biosynthesis I (meso-diaminopimelate containing** | diff_to_baseine | diff_to_baseine | diff_to_baseine | -0.00377148554170626 | 0.00486976258617576 | 0.0065173274751329 | 0.397520561139666 | 0.925861580484465 | 0.397520561139666 | 0.925861580484465 | NA | abundance | 27 | 27 |
| **PWY-5686: UMP biosynthesis I** | diff_to_baseine | diff_to_baseine | diff_to_baseine | -0.00525717228845906 | 0.00486976258617576 | 0.00636740806922145 | 0.32092916695806 | 0.925861580484465 | 0.32092916695806 | 0.925861580484465 | NA | abundance | 27 | 27 |
| **PWY-6163: chorismate biosynthesis from 3-dehydroquinate** | diff_to_baseine | diff_to_baseine | diff_to_baseine | 0.00914221898755687 | 0.00486976258617576 | 0.00739869289946606 | 0.690196223167294 | 0.925861580484465 | 0.690196223167294 | 0.925861580484465 | NA | abundance | 27 | 27 |
| **PWY-6590: superpathway of Clostridium acetobutylicum acidogenic fermentation** | diff_to_baseine | diff_to_baseine | diff_to_baseine | 0.0182418711953176 | 0.00486976258617576 | 0.0268068054985364 | 0.632487523215742 | 0.925861580484465 | 0.632487523215742 | 0.925861580484465 | NA | abundance | 27 | 27 |
| **PWY-6629: superpathway of L-tryptophan biosynthesis** | diff_to_baseine | diff_to_baseine | diff_to_baseine | 0.00942092542932678 | 0.00486976258617576 | 0.0057759650260708 | 0.641508408526958 | 0.925861580484465 | 0.641508408526958 | 0.925861580484465 | NA | abundance | 27 | 27 |
| **PWY-6969: TCA cycle V (2-oxoglutarate synthase)** | diff_to_baseine | diff_to_baseine | diff_to_baseine | 0.0165235514309735 | 0.00486976258617576 | 0.0168946269410437 | 0.532558822162461 | 0.925861580484465 | 0.532558822162461 | 0.925861580484465 | NA | abundance | 27 | 27 |
| **PWY-7851: coenzyme A biosynthesis II (eukaryotic)** | diff_to_baseine | diff_to_baseine | diff_to_baseine | 0.00254264916237993 | 0.00486976258617576 | 0.00644172025984745 | 0.814758190826329 | 0.925861580484465 | 0.814758190826329 | 0.925861580484465 | NA | abundance | 27 | 27 |
| **TRPSYN-PWY: L-tryptophan biosynthesis** | diff_to_baseine | diff_to_baseine | diff_to_baseine | 0.0106189358333386 | 0.00486976258617576 | 0.00527848045680571 | 0.546728014186002 | 0.925861580484465 | 0.546728014186002 | 0.925861580484465 | NA | abundance | 27 | 27 |
| **VALSYN-PWY: L-valine biosynthesis** | diff_to_baseine | diff_to_baseine | diff_to_baseine | -0.00133815851764987 | 0.00486976258617576 | 0.00492995002496726 | 0.502498492726803 | 0.925861580484465 | 0.502498492726803 | 0.925861580484465 | NA | abundance | 27 | 27 |
| **COA-PWY: coenzyme A biosynthesis I (prokaryotic)** | diff_to_baseine | diff_to_baseine | diff_to_baseine | 0.00327634236230274 | 0.00486976258617576 | 0.00653941897333436 | 0.873395103402114 | 0.94934250369795 | 0.873395103402114 | 0.94934250369795 | NA | abundance | 27 | 27 |
| **ARGSYNBSUB-PWY: L-arginine biosynthesis II (acetyl cycle)** | diff_to_baseine | diff_to_baseine | diff_to_baseine | 0.00572353113869459 | 0.00486976258617576 | 0.00631517614764713 | 0.931350694781923 | 0.97015697373117 | 0.931350694781923 | 0.97015697373117 | NA | abundance | 27 | 27 |
| **ARGSYN-PWY: L-arginine biosynthesis I (via L-ornithine)** | diff_to_baseine | diff_to_baseine | diff_to_baseine | 0.00486976258617576 | 0.00486976258617576 | 0.0058799728971423 | 1 | 1 | 1 | 1 | NA | abundance | 27 | 27 |
| **ARGSYN-PWY: L-arginine biosynthesis I (via L-ornithine)** | diff_to_baseine | diff_to_baseine | diff_to_baseine | NA | 0 | NA | NA | NA | 1 | 1 | All logistic values are the same | prevalence | 27 | 27 |
| **ARGSYNBSUB-PWY: L-arginine biosynthesis II (acetyl cycle)** | diff_to_baseine | diff_to_baseine | diff_to_baseine | NA | 0 | NA | NA | NA | 0.931350694781923 | 0.97015697373117 | All logistic values are the same | prevalence | 27 | 27 |
| **ARO-PWY: chorismate biosynthesis I** | diff_to_baseine | diff_to_baseine | diff_to_baseine | NA | 0 | NA | NA | NA | 0.765677864069327 | 0.925861580484465 | All logistic values are the same | prevalence | 27 | 27 |
| **BRANCHED-CHAIN-AA-SYN-PWY: superpathway of branched chain amino acid biosynthesis** | diff_to_baseine | diff_to_baseine | diff_to_baseine | NA | 0 | NA | NA | NA | 0.64325454100129 | 0.925861580484465 | All logistic values are the same | prevalence | 27 | 27 |
| **CENTFERM-PWY: pyruvate fermentation to butanoate** | diff_to_baseine | diff_to_baseine | diff_to_baseine | NA | 0 | NA | NA | NA | 0.623670817236797 | 0.925861580484465 | All logistic values are the same | prevalence | 27 | 27 |
| **COA-PWY-1: superpathway of coenzyme A biosynthesis III (mammals)** | diff_to_baseine | diff_to_baseine | diff_to_baseine | NA | 0 | NA | NA | NA | 0.779451238021495 | 0.925861580484465 | All logistic values are the same | prevalence | 27 | 27 |
| **COA-PWY: coenzyme A biosynthesis I (prokaryotic)** | diff_to_baseine | diff_to_baseine | diff_to_baseine | NA | 0 | NA | NA | NA | 0.873395103402114 | 0.94934250369795 | All logistic values are the same | prevalence | 27 | 27 |
| **COMPLETE-ARO-PWY: superpathway of aromatic amino acid biosynthesis** | diff_to_baseine | diff_to_baseine | diff_to_baseine | NA | 0 | NA | NA | NA | 0.737889157635735 | 0.925861580484465 | All logistic values are the same | prevalence | 27 | 27 |
| **FERMENTATION-PWY: mixed acid fermentation** | diff_to_baseine | diff_to_baseine | diff_to_baseine | NA | 0 | NA | NA | NA | 0.606439414085784 | 0.925861580484465 | All logistic values are the same | prevalence | 27 | 27 |
| **FOLSYN-PWY: superpathway of tetrahydrofolate biosynthesis and salvage** | diff_to_baseine | diff_to_baseine | diff_to_baseine | NA | 0 | NA | NA | NA | 0.00844490981944305 | 0.074786057918652 | All logistic values are the same | prevalence | 27 | 27 |
| **HISTSYN-PWY: L-histidine biosynthesis** | diff_to_baseine | diff_to_baseine | diff_to_baseine | NA | 0 | NA | NA | NA | 0.449317639423979 | 0.925861580484465 | All logistic values are the same | prevalence | 27 | 27 |
| **NAGLIPASYN-PWY: lipid IVA biosynthesis (E. coli)** | diff_to_baseine | diff_to_baseine | diff_to_baseine | NA | 0 | NA | NA | NA | 0.357496237264372 | 0.925861580484465 | All logistic values are the same | prevalence | 27 | 27 |
| **PEPTIDOGLYCANSYN-PWY: peptidoglycan biosynthesis I (meso-diaminopimelate containing** | diff_to_baseine | diff_to_baseine | diff_to_baseine | NA | 0 | NA | NA | NA | 0.397520561139666 | 0.925861580484465 | All logistic values are the same | prevalence | 27 | 27 |
| **PWY-5676: acetyl-CoA fermentation to butanoate II** | diff_to_baseine | diff_to_baseine | diff_to_baseine | NA | 0 | NA | NA | NA | 0.148775579356197 | 0.743877896780986 | All logistic values are the same | prevalence | 27 | 27 |
| **PWY-5686: UMP biosynthesis I** | diff_to_baseine | diff_to_baseine | diff_to_baseine | NA | 0 | NA | NA | NA | 0.32092916695806 | 0.925861580484465 | All logistic values are the same | prevalence | 27 | 27 |
| **PWY-6163: chorismate biosynthesis from 3-dehydroquinate** | diff_to_baseine | diff_to_baseine | diff_to_baseine | NA | 0 | NA | NA | NA | 0.690196223167294 | 0.925861580484465 | All logistic values are the same | prevalence | 27 | 27 |
| **PWY-6590: superpathway of Clostridium acetobutylicum acidogenic fermentation** | diff_to_baseine | diff_to_baseine | diff_to_baseine | NA | 0 | NA | NA | NA | 0.632487523215742 | 0.925861580484465 | All logistic values are the same | prevalence | 27 | 27 |
| **PWY-6612: superpathway of tetrahydrofolate biosynthesis** | diff_to_baseine | diff_to_baseine | diff_to_baseine | NA | 0 | NA | NA | NA | 0.00862244897214534 | 0.074786057918652 | All logistic values are the same | prevalence | 27 | 27 |
| **PWY-6629: superpathway of L-tryptophan biosynthesis** | diff_to_baseine | diff_to_baseine | diff_to_baseine | NA | 0 | NA | NA | NA | 0.641508408526958 | 0.925861580484465 | All logistic values are the same | prevalence | 27 | 27 |
| **PWY-6969: TCA cycle V (2-oxoglutarate synthase)** | diff_to_baseine | diff_to_baseine | diff_to_baseine | NA | 0 | NA | NA | NA | 0.532558822162461 | 0.925861580484465 | All logistic values are the same | prevalence | 27 | 27 |
| **PWY-7851: coenzyme A biosynthesis II (eukaryotic)** | diff_to_baseine | diff_to_baseine | diff_to_baseine | NA | 0 | NA | NA | NA | 0.814758190826329 | 0.925861580484465 | All logistic values are the same | prevalence | 27 | 27 |
| **PWY0-1297: superpathway of purine deoxyribonucleosides degradation** | diff_to_baseine | diff_to_baseine | diff_to_baseine | NA | 0 | NA | NA | NA | 0.00897432695023823 | 0.074786057918652 | All logistic values are the same | prevalence | 27 | 27 |
| **PWY66-409: superpathway of purine nucleotide salvage** | diff_to_baseine | diff_to_baseine | diff_to_baseine | NA | 0 | NA | NA | NA | 0.0284115110850622 | 0.177571944281638 | All logistic values are the same | prevalence | 27 | 27 |
| **TRPSYN-PWY: L-tryptophan biosynthesis** | diff_to_baseine | diff_to_baseine | diff_to_baseine | NA | 0 | NA | NA | NA | 0.546728014186002 | 0.925861580484465 | All logistic values are the same | prevalence | 27 | 27 |
| **VALSYN-PWY: L-valine biosynthesis** | diff_to_baseine | diff_to_baseine | diff_to_baseine | NA | 0 | NA | NA | NA | 0.502498492726803 | 0.925861580484465 | All logistic values are the same | prevalence | 27 | 27 |

**Supplementary Table 5: CHAI domain analysis (Maaslin3), significant features, abundance model**

| | feature | metadata | value | name | coef | null_hypothesis | stderr | pval_individual | qval_individual | pval_joint | qval_joint | error | model | N | N_not_zero | model_type | domain | q_global |
|---|---|---|---|---|---|---|---|---|---|---|---|---|---|---|---|---|---|---|
| 1 | Lachnoclostridium | ratio_cyto | ratio_cyto | ratio_cyto | -2.30509994740504 | -0.0526659990278388 | 0.486896761507109 | 0.000267731965048793 | 0.118337528551567 | 0.000267731965048793 | 0.06211381589132 | NA | linear | 27 | 22 | abundance | ratio_cyto | 0.0542968009425943 |
| 2 | Alistipes | ratio_gut | ratio_gut | ratio_gut | -3.11175566118097 | -0.461364713778247 | 0.724252428185172 | 0.00186659498072095 | 0.200167088767341 | 0.00372970578461986 | 0.206073182955149 | NA | linear | 27 | 25 | abundance | ratio_gut | 0.209949559430679 |
| 3 | Escherichia | ratio_gut | ratio_gut | ratio_gut | 1.6825182616622 | -0.461364713778247 | 0.541467307116 | 0.00250365743784456 | 0.200167088767341 | 0.00500104657512304 | 0.206073182955149 | NA | linear | 27 | 21 | abundance | ratio_gut | 0.209949559430679 |
| 4 | Veillonella | ratio_gut | ratio_gut | ratio_gut | 1.85544353465479 | -0.461364713778247 | 0.640527188266963 | 0.00307590694711475 | 0.200167088767341 | 0.00614235269068219 | 0.206073182955149 | NA | linear | 27 | 18 | abundance | ratio_gut | 0.225694672244545 |
| 5 | GGB9715 | ratio_cyto | ratio_cyto | ratio_cyto | 1.75548229466966 | -0.0526659990278388 | 0.498159853814011 | 0.00352441711274116 | 0.519264121277198 | 0.00703641270949774 | 0.544149249534492 | NA | linear | 27 | 16 | abundance | ratio_cyto | 0.229870316131007 |
| 6 | Limosilactobacillus | ratio_gut | ratio_gut | ratio_gut | 1.31278722210406 | -0.461364713778247 | 0.430515769770544 | 0.00465888552740809 | 0.262062310916705 | 0.0092960658404587 | 0.269585909373302 | NA | linear | 27 | 11 | abundance | ratio_gut | 0.273476580458855 |
| 7 | Dialister | ratio_cyto | ratio_cyto | ratio_cyto | -3.87641814070947 | -0.0526659990278388 | 1.21082058168098 | 0.00630614725457498 | 0.632633039318742 | 0.0125725270159536 | 0.6617451745223 | NA | linear | 27 | 19 | abundance | ratio_cyto | 0.307606496827664 |
| 8 | Lactobacillus | ratio_gut | ratio_gut | ratio_gut | 3.6279152308911 | -0.461364713778247 | 0.852423615114717 | 0.00716332912908696 | 0.33013985179119 | 0.0142753449739623 | 0.308329242952018 | NA | linear | 27 | 11 | abundance | ratio_gut | 0.307606496827664 |
| 9 | Bifidobacterium | ratio_lung | ratio_lung | ratio_lung | -1.41144978475208 | 0.0388300276987826 | 0.460270992505239 | 0.00894144889998463 | 0.83828499064691 | 0.00894144889998463 | 0.890108230962996 | NA | linear | 20 | 20 | abundance | ratio_lung | 0.318973059259068 |
| 10 | GGB34797 | ratio_lymph | ratio_lymph | ratio_lymph | -1.16828030017073 | 0.10745163079374 | 0.263284569018175 | 0.00907721215495393 | 1 | 0.0180720285294018 | 0.995817319118829 | NA | linear | 26 | 9 | abundance | ratio_lymph | 0.318973059259068 |
| 11 | GGB4604 | ratio_immun | ratio_immun | ratio_immun | -1.97158942149764 | 0.163312103696631 | 0.51518876052311 | 0.00923772062590144 | 0.876790003664405 | 0.0183901057694407 | 0.856298689617484 | NA | linear | 27 | 8 | abundance | ratio_immun | 0.318973059259068 |
| 12 | Actinomyces | ratio_lymph | ratio_lymph | ratio_lymph | -1.10441840859452 | 0.10745163079374 | 0.449126648372216 | 0.0143246080285288 | 1 | 0.0284440216618866 | 0.995817319118829 | NA | linear | 26 | 25 | abundance | ratio_lymph | 0.39043116612757 |
| 13 | Ruminococcus | ratio_gut | ratio_gut | ratio_gut | -2.68001247782139 | -0.461364713778247 | 0.801956367456726 | 0.0154476585139341 | 0.496531880805024 | 0.030656686874305 | 0.474156756989251 | NA | linear | 27 | 23 | abundance | ratio_gut | 0.394251110768665 |
| 14 | GGB3109 | ratio_immun | ratio_immun | ratio_immun | -1.01805878673731 | 0.163312103696631 | 0.412724207870233 | 0.0201864312969261 | 0.876790003664405 | 0.0399653705853467 | 0.856298689617484 | NA | linear | 27 | 12 | abundance | ratio_immun | 0.455747506588294 |
| 15 | GGB6613 | ratio_cyto | ratio_cyto | ratio_cyto | -1.35041981075823 | -0.0526659990278388 | 0.466421388025063 | 0.0216344479116513 | 0.78193548414832 | 0.0428008464868606 | 0.81141629214839 | NA | linear | 27 | 13 | abundance | ratio_cyto | 0.45760712773886 |
| 16 | GGB1420 | ratio_cyto | ratio_cyto | ratio_cyto | -1.15198088317549 | -0.0526659990278388 | 0.288443698338955 | 0.0229981024749506 | 0.78193548414832 | 0.0454672922324529 | 0.81141629214839 | NA | linear | 27 | 7 | abundance | ratio_cyto | 0.45760712773886 |
| 17 | Gordonibacter | ratio_lymph | ratio_lymph | ratio_lymph | -1.51387762483969 | 0.10745163079374 | 0.655682020650894 | 0.0295437880804054 | 1 | 0.0582147407466709 | 0.995817319118829 | NA | linear | 26 | 16 | abundance | ratio_lymph | 0.502128930846725 |
| 18 | Lachnoclostridium | ratio_gut | ratio_gut | ratio_gut | -1.7228200219439 | -0.461364713778247 | 0.504287942055816 | 0.0328573138326933 | 0.598577945837562 | 0.0646350245930864 | 0.606938078315422 | NA | linear | 27 | 22 | abundance | ratio_gut | 0.529575866213035 |
| 19 | Escherichia | ratio_cyto | ratio_cyto | ratio_cyto | 1.63584737152076 | -0.0526659990278388 | 0.74099136407745 | 0.0382158884553712 | 0.986404523297104 | 0.074971322780309 | 0.96504980487362 | NA | linear | 27 | 21 | abundance | ratio_cyto | 0.575198115982125 |
| 20 | GGB52130 | ratio_gut | ratio_gut | ratio_gut | -3.37458652291773 | -0.461364713778247 | 1.25251699928877 | 0.0418052708476069 | 0.613102250189133 | 0.0818628610245721 | 0.618395605148873 | NA | linear | 27 | 14 | abundance | ratio_gut | 0.612751537257168 |
| 21 | Intestinimonas | ratio_lymph | ratio_lymph | ratio_lymph | 1.15938675419711 | 0.10745163079374 | 0.435904963717434 | 0.0427986593314206 | 1 | 0.0837655934222742 | 0.995817319118829 | NA | linear | 26 | 16 | abundance | ratio_lymph | 0.612751537257168 |
| 22 | Rothia | ratio_lung | ratio_lung | ratio_lung | 1.57462456224748 | 0.0388300276987826 | 0.655251092858015 | 0.0452651826527525 | 0.83828499064691 | 0.0884814285449179 | 0.890108230962996 | NA | linear | 20 | 16 | abundance | ratio_lung | 0.63195467219965 |
| 23 | Lachnoclostridium | ratio_lymph | ratio_lymph | ratio_lymph | -1.31628130690291 | 0.10745163079374 | 0.665708719841498 | 0.0498253820272871 | 1 | 0.097168195360409 | 0.995817319118829 | NA | linear | 26 | 21 | abundance | ratio_lymph | 0.63195467219965 |
| 24 | Oliverpabstia | ratio_lung | ratio_lung | ratio_lung | -3.40329662879709 | 0.0388300276987826 | 0.83221495769008 | 0.051790683182988 | 0.83828499064691 | 0.100899091501415 | 0.890108230962996 | NA | linear | 20 | 6 | abundance | ratio_lung | 0.63195467219965 |
| 25 | Lawsonibacter | ratio_gut | ratio_gut | ratio_gut | -2.59368432227265 | -0.461364713778247 | 1.02519477860356 | 0.0522177864965201 | 0.654679318775824 | 0.101708875766444 | 0.639592224169068 | NA | linear | 27 | 24 | abundance | ratio_gut | 0.63195467219965 |
| 26 | Zhenpiania | ratio_gut | ratio_gut | ratio_gut | -3.98747957331731 | -0.461364713778247 | 1.46684357335579 | 0.0530639589970371 | 0.654679318775824 | 0.103312134249635 | 0.639592224169068 | NA | linear | 27 | 10 | abundance | ratio_gut | 0.63195467219965 |
| 27 | Anaerotruncus | ratio_cyto | ratio_cyto | ratio_cyto | -1.61367696044979 | -0.0526659990278388 | 0.742091506326574 | 0.0538289865309819 | 0.986404523297104 | 0.104760413271011 | 0.96504980487362 | NA | linear | 27 | 19 | abundance | ratio_cyto | 0.63195467219965 |
| 28 | Faecalibacillus | ratio_lung | ratio_lung | ratio_lung | 1.96748936683592 | 0.0388300276987826 | 0.879079795612819 | 0.0621665816709265 | 0.83828499064691 | 0.120468479465205 | 0.890108230962996 | NA | linear | 20 | 14 | abundance | ratio_lung | 0.68852421586479 |
| 29 | Anaerotignum | ratio_gut | ratio_gut | ratio_gut | 1.04613208573784 | -0.461364713778247 | 0.708660036728595 | 0.0687583386781273 | 0.687583386781273 | 0.132788968218478 | 0.669718274493196 | NA | linear | 27 | 15 | abundance | ratio_gut | 0.72919627049146 |
| 30 | GGB34797 | ratio_cyto | ratio_cyto | ratio_cyto | -1.17380543342383 | -0.0526659990278388 | 0.473244360632062 | 0.0689697471599694 | 0.986404523297104 | 0.133182668296629 | 0.96504980487362 | NA | linear | 27 | 9 | abundance | ratio_cyto | 0.72919627049146 |
| 31 | Lachnospiraceae_unclassified | ratio_lung | ratio_lung | ratio_lung | -1.34373380574003 | 0.0388300276987826 | 0.687610415953265 | 0.0736220032316064 | 0.83828499064691 | 0.141823807103378 | 0.890108230962996 | NA | linear | 20 | 15 | abundance | ratio_lung | 0.734471109177799 |
| 32 | GGB4604 | ratio_lymph | ratio_lymph | ratio_lymph | -1.34307117977125 | 0.10745163079374 | 0.641083772552354 | 0.0746070587146945 | 1 | 0.143647904219331 | 0.995817319118829 | NA | linear | 26 | 8 | abundance | ratio_lymph | 0.734471109177799 |
| 33 | Lentihominibacter | ratio_cyto | ratio_cyto | ratio_cyto | 1.24030201242531 | -0.0526659990278388 | 0.539819816551899 | 0.0775761648535324 | 0.986404523297104 | 0.149134268353682 | 0.96504980487362 | NA | linear | 27 | 15 | abundance | ratio_cyto | 0.734471109177799 |
| 34 | Limosilactobacillus | ratio_cyto | ratio_cyto | ratio_cyto | 1.64547967228154 | -0.0526659990278388 | 0.838369096456713 | 0.0792005690629545 | 0.986404523297104 | 0.152128407986013 | 0.96504980487362 | NA | linear | 27 | 11 | abundance | ratio_cyto | 0.734915231494309 |
| 35 | GGB3109 | ratio_gut | ratio_gut | ratio_gut | 1.39221676488438 | -0.461364713778247 | 0.884487013987563 | 0.0803996757692371 | 0.753746960336598 | 0.154335243674676 | 0.730730133316832 | NA | linear | 27 | 12 | abundance | ratio_gut | 0.734915231494309 |
| 36 | Candidatus_Cibionibacter | ratio_lung | ratio_lung | ratio_lung | -1.13257695514539 | 0.0388300276987826 | 0.525584428498771 | 0.0813790290411074 | 0.83828499064691 | 0.156135511714541 | 0.890108230962996 | NA | linear | 20 | 8 | abundance | ratio_lung | 0.734915231494309 |
| 37 | Escherichia | ratio_lymph | ratio_lymph | ratio_lymph | 1.29749953391519 | 0.10745163079374 | 0.648569589082302 | 0.0931348111407062 | 1 | 0.177595529235197 | 0.995817319118829 | NA | linear | 26 | 20 | abundance | ratio_lymph | 0.746482454459885 |
| 38 | Eubacterium | ratio_lung | ratio_lung | ratio_lung | -1.23405062351261 | 0.0388300276987826 | 0.694584548638328 | 0.0958184113224463 | 0.83828499064691 | 0.182455654696535 | 0.890108230962996 | NA | linear | 20 | 15 | abundance | ratio_lung | 0.746482454459885 |

**Supplementary Table 6: CHAI domain analysis (Maaslin3), significant features, prevalence model**

| | feature | metadata | value | name | coef | null_hypothesis | stderr | pval_individual | qval_individual | pval_joint | qval_joint | error | model | N | N_not_zero | model_type | domain | q_global |
|---|---|---|---|---|---|---|---|---|---|---|---|---|---|---|---|---|---|---|
| 1 | Hungatella | ratio_gut | ratio_gut | ratio_gut | -1.59900128875834 | 0 | 0.692612715043396 | 0.0209629352855286 | 0.505878761331246 | 0.0414864259152719 | 0.515460126741942 | NA | logistic | 27 | 18 | prevalence | ratio_gut | 1 |
| 2 | Faecalicatena | ratio_gut | ratio_gut | ratio_gut | -1.50223915941993 | 0 | 0.705608152067859 | 0.033254330324309 | 0.598577945837562 | 0.0654028101632998 | 0.606938078315422 | NA | logistic | 27 | 20 | prevalence | ratio_gut | 1 |
| 3 | Intestinimonas | ratio_gut | ratio_gut | ratio_gut | -1.43858124674297 | 0 | 0.710401064429031 | 0.0428646047934182 | 0.613102250189133 | 0.0838918352427404 | 0.618395605148873 | NA | logistic | 27 | 17 | prevalence | ratio_gut | 1 |
| 4 | Anaerotruncus | ratio_gut | ratio_gut | ratio_gut | -1.39448241944241 | 0 | 0.691045423956314 | 0.0435983822356717 | 0.613102250189133 | 0.0852959455377757 | 0.618395605148873 | NA | logistic | 27 | 19 | prevalence | ratio_gut | 1 |
| 5 | Faecalicatena | ratio_cyto | ratio_cyto | ratio_cyto | -2.10370491096666 | 0 | 1.04592908592101 | 0.044290978832542 | 0.986404523297104 | 0.0866202668591392 | 0.96504980487362 | NA | logistic | 27 | 20 | prevalence | ratio_cyto | 1 |
| 6 | Zhenpiania | ratio_immun | ratio_immun | ratio_immun | 1.47240924502183 | 0 | 0.77159092228878 | 0.0563554182861586 | 0.876790003664405 | 0.109534903402109 | 0.856298689617484 | NA | logistic | 27 | 10 | prevalence | ratio_immun | 1 |
| 7 | Oscillibacter | ratio_gut | ratio_gut | ratio_gut | -1.2476873672681 | 0 | 0.669104506146948 | 0.0622217491597205 | 0.656078113524133 | 0.120571952250946 | 0.640240168498096 | NA | logistic | 27 | 20 | prevalence | ratio_gut | 1 |
| 8 | Eisenbergiella | ratio_gut | ratio_gut | ratio_gut | -1.21933739257146 | 0 | 0.658524904255231 | 0.0640805161010508 | 0.656078113524133 | 0.12405471965832 5 | 0.640240168498096 | NA | logistic | 27 | 13 | prevalence | ratio_gut | 1 |
| 9 | Oscillospiraceae_unclassified | ratio_gut | ratio_gut | ratio_gut | -1.26683717367344 | 0 | 0.684229838194391 | 0.0641006737471171 | 0.656078113524133 | 0.1240924511194 | 0.640240168498096 | NA | logistic | 27 | 22 | prevalence | ratio_gut | 1 |
| 10 | Pseudoflavonifractor | ratio_gut | ratio_gut | ratio_gut | -1.0391465453615 | 0 | 0.594126470074966 | 0.0802853900607624 | 0.753746960336598 | 0.154125036264316 | 0.730730133316832 | NA | logistic | 27 | 12 | prevalence | ratio_gut | 1 |
| 11 | Anaerotruncus | ratio_cyto | ratio_cyto | ratio_cyto | -1.68286401355331 | 0 | 0.962990714512853 | 0.080543853033476 | 0.986404523297104 | 0.104760413271011 | 0.96504980487362 | NA | logistic | 27 | 19 | prevalence | ratio_cyto | 1 |
| 12 | Oscillibacter | ratio_cyto | ratio_cyto | ratio_cyto | -1.5860364332797 | 0 | 0.936008059696082 | 0.0901762397003061 | 0.986404523297104 | 0.172220725194125 | 0.96504980487362 | NA | logistic | 27 | 20 | prevalence | ratio_cyto | 1 |
| 13 | GGB3109 | ratio_cyto | ratio_cyto | ratio_cyto | -1.73117212474206 | 0 | 1.04532512004363 | 0.0976997886217513 | 0.986404523297104 | 0.0976997886217513 | 0.96504980487362 | NA | logistic | 27 | 12 | prevalence | ratio_cyto | 1 |
| 14 | GGB2653 | ratio_gut | ratio_gut | ratio_gut | -1.15097685158137 | 0 | 0.698245921842969 | 0.0992740599260395 | 0.794064526122834 | 0.18869278087788 | 0.777592258298231 | NA | logistic | 27 | 12 | prevalence | ratio_gut | 1 |
